# Topographic organization of bidirectional connections between the cingulate region (infralimbic area and anterior cingulate area, dorsal part) and the interbrain (diencephalon) of the adult male rat

**DOI:** 10.1101/2024.09.29.615708

**Authors:** Kenichiro Negishi, Laura P. Montes, Vanessa I. Navarro, Lidice Soto Arzate, Cindy Oliveros, Arshad M. Khan

**Affiliations:** UTEP Systems Neuroscience Laboratory, Department of Biological Sciences, The University of Texas at El Paso, El Paso, Texas, 79968, USA; Ph.D. Program in Bioscience, Department of Biological Sciences, The University of Texas at El Paso, El Paso, Texas, 79968, USA; Undergraduate Baccalaureate Program in Nursing, College of Nursing, The University of Texas at El Paso, El Paso, Texas, 79968, USA; Border Biomedical Research Center, The University of Texas at El Paso, El Paso, Texas, 79968, USA; Behavioral Neuroscience Branch, IRP/NIDA/NIH, Baltimore, MD 21224, USA

**Keywords:** prefrontal cortex, mapping, brain atlas, feeding, lateral hypothalamic area, anterograde, retrograde

## Abstract

The medial prefrontal cortex [*cingulate region (Brodmann, 1909)* (CNG)] in the rat is a connectionally and functionally diverse structure. It harbors cerebral nuclei that use long-range connections to promote adaptive changes to ongoing behaviors. The CNG is often described across functional and anatomical gradients, a dorsal-ventral gradient being the most prominent. Topographic organization is a general feature of the nervous system, and it is becoming clear that such spatial arrangements can reflect connectional, functional, and cellular differences. Portions of the CNG are known to form reciprocal connections with cortical areas and thalamus; however, these connectional features have not been described in detail or mapped to standardized rat brain atlases. Here, we used co-injected anterograde (*Phaseolus vulgaris* leucoagglutinin) and retrograde (cholera toxin B subunit) tracers throughout the CNG to identify zones of reciprocal connectivity in the diencephalon [or *interbrain (Baer, 1837)* (IB)]. Tracer distributions were observed using a Nissl-based atlas-mapping approach that facilitates description of topographic organization. This draft report describes CNG connections of the *infralimbic area (Rose & Woolsey, 1948)* (ILA) and *the anterior cingulate area, dorsal part (Krettek & Price, 1977)* (ACAd) throughout the IB. We found that corticothalamic connections are predominantly reciprocal, and that ILA and ACAd connections tended to be spatially segregated with minimal overlap. In the *hypothalamus (Kuhlenbeck, 1927)*, we found dense and specific ILA-originating terminals in the following *Brain Maps 4.0 at*las territories: *dorsal region (Swanson, 2004)* (LHAd) and *suprafornical region (Swanson, 2004)* (LHAs) of the *lateral hypothalamic area (Nissl, 1913)*, *parasubthalamic nucleus (Wang & Zhang, 1995)* (PSTN), *tuberal nucleus, terete part (Petrovich et al., 2001)* (TUte), and an ill-defined dorsal cap of the *medial mammillary nucleus (Gudden, 1881)* (MM). We discuss these findings in the context of feeding behaviors.

## 1. Introduction

An area of the medial prefrontal region of the cerebral cortex, which we more precisely define here as the *cingulate region (Brodmann, 1909)* (CNG; see *Section 2.1* for nomenclature used in this study), is critical for making adaptive changes to ongoing behaviors (Miller and Cohen, 2001). These functions leverage excitatory outputs to every major division of the central nervous system (Gabbott et al., 2005). Interestingly, the medial prefrontal cortex is reportedly the last brain region to fully mature (Paus et al., 1999), leaving wide the opportunity for the environment and chance to shape the dynamics of behavior. It is therefore appropriate that considerable attention has been committed towards understanding the medial prefrontal cortex and how it contributes to mental illness, substance use disorders, and aberrations of motivated behaviors (Sapolsky, 2004).

Classical experiments with decerebrate animals indicated that the diencephalon (alternatively, *interbrain (Baer, 1837)* (IB) [Swanson, 2015]) is likely necessary for the expression of motivated behaviors, as no spontaneous or intact behaviors were observed in these preparations (Grill and Norgren, 1978). Decorticate animals, on the other hand, were able to perform motivated behaviors but failed to integrate environmental information as indicated by their expression at inappropriate times, places, and a near-complete lack of anticipatory and social behaviors (Vanderwolf et al., 1978). At a structural level, motivated behaviors are supported by distributed and interconnected brain regions that are often described in a highly schematized manner to better facilitate an understanding of their complex organization (Watts et al., 2022).

Pursuing a comprehensive “wiring diagram” for motivated behaviors is technically challenging due to the well-recognized but poorly addressed phenomenon of topographic organization. That is, connections within discrete gray matter regions often form gradients or compartmental organization (Hintiryan et al., 2016; Eickhoff et al., 2018) that require careful parsing across three dimensions. Moreover, topography is increasingly observed to align connectional, functional, cellular and gene-expression differences within individual brain regions (Mandelbaum et al., 2019; Mickelsen et al., 2020). Even documentation of connectional topography alone will likely identify meaningful spatial patterns when merged with relevant functional and transcriptomic datasets, especially if those datasets are placed within standardized atlases for others to use and interrelate with their own datasets (Khan, 2013; Khan et al., 2018a).

We accordingly examined CNG connections using a Nissl-based atlas-mapping approach that is profitably employed to uncover sub-regional architecture (Simmons and Swanson, 2009a). The CNG is commonly studied along a dorsoventral functional and anatomical gradient. The *infralimbic area (Rose & Woolsey, 1948)* (ILA) and *anterior cingulate area, dorsal part (Krettek & Price, 1977)* (ACAd) are on opposite ends of this gradient, making them excellent starting points for elaborating CNG connections. Bidirectional connections were revealed for these structures by co-injecting the anterograde and retrograde tracers *Phaseolus vulgaris* leucoagglutinin (PHAL) and cholera toxin B subunit (CTB) throughout the perigenual CNG.

Here, we present a draft report of the complete mappings of ILA/ACAd bidirectional connections with the IB with the goal of clarifying their precise spatial arrangements. This work, to our knowledge, is the first detailed representation of reciprocal connectivity between these CNG structures and the IB. Our mapping of ILA and ACAd connections in the present study, portions of which have been presented in preliminary form (Negishi et al., 2015, 2017, 2019; Negishi, 2016; 2023), prompts interpretive refinements to various thalamic and hypothalamic regions, especially regarding CNG connections with the *lateral hypothalamic zone (Nauta & Haymaker, 1969)* (HYl) and *zona incerta (>1840)* (ZI).

## 2. Materials and Methods

### 2.1 Neuroanatomical nomenclature

In this study, we follow the standardized nomenclature from the *Brain Maps 4.0* rat brain atlas (Swanson, 2018), which is derived from the lexicon developed by Swanson (2015). In this system, the formal form *term (author, date)* comprises a *standard term* for a neuroanatomical structure. The citation portion of the formal term refers to the first use of the term as it is defined by Swanson (2015, 2018). Standard terms for which such priority has not yet been established is assigned a value “*(>1840)*” to denote that the term was used approximately after the formulation of cell theory.

### 2.2 Animals

Experiments were performed on adult male Sprague-Dawley rats (Harlan Labs, Indianapolis, IN) weighing 300–400 g. Animals were fed ad libitum and housed in a temperature-controlled vivarium under a 12-hour day/ night cycle (lights on at 0700 h). All methods followed protocols approved by the The University of Texas at El Paso Institutional Animal Care and Use Committee (Protocol # A-201207-1) and were in accordance with the NIH *Guidelines for the Care and Use of Laboratory Animals* (NRC, 2011).

### 2.3 Intracranial tracer injection

Rats were anesthetized with a mixture containing 50% ketamine, 5% xylazine, 10% acepromazine and 35% sterile saline (1 µL/g). Once anesthetized, animals were positioned in a Kopf stereotaxic frame (David Kopf Instruments, Tujunga, California, USA) and maintained on 1.5% isoflurane delivered with pure oxygen for the duration of the surgery. The frontal bone and bregma fiducials were carefully exposed for craniotomy. Glass micropipettes with inner tip diameters ranging from 12 to 20 µm were selected and filled with a cocktail containing 2.5% PHAL (Vector Laboratories, Inc., Newark, California, USA; catalog #L-1110) and 0.25% CTB (List Biological Labs, Inc., Campbell, California, USA; catalog #104) dissolved in 10 mM sodium phosphate solution. Stereotaxic coordinates targeting the ILA (AP +11.20 mm, ML –0.50 mm and DV –4.40 mm) and ACAd (AP +11.20 mm, ML –0.50 mm and DV –2.40 mm) were obtained using the Paxinos and Watson (2014) rat brain atlas. Tracers were ejected iontophoretically at a current of 5 µA through 7 sec on/off cycles for 10–15 min. Micropipettes were retracted slowly after a resting period of 10 min. Following surgery, animals received intramuscular injections of Flunazine (Bimeda-MTC Animal Health, Inc., catalog #200-387) as an analgesic and Gentamicin (Vedco, Inc., St. Joseph, Missouri, USA; catalog #50989-040-12) for its antimicrobial and anti-inflammatory properties. Another injection of Flunazine was given 8 h after surgery if grimaces or other signs of pain were detected. Daily status evaluations were maintained for a survival period of 10–14 d to allow for tracer transport.

### 2.4 Transcardial perfusion and tissue preparation

Animals were sedated with isoflurane for two min and perfused transcardially with 200 mL of phosphate-buffered saline (PBS; pH 7.4 at room temperature), followed by fixation with 350 mL of ice-cold 4% paraformaldehyde (PFA) in 0.05 M PBS (pH 7.4 at room temperature). Brains were carefully dissected and post-fixed overnight in the same PFA solution at 4°C. Brains were transferred to PBS containing 10% (w/v) sucrose until they appeared fully saturated. Fixed brains were then blocked (a coronal cut at the level of the caudal end of the mammillary body) and frozen with dry ice on a custom stage fitted to an OmE sliding microtome (Reichert, Austria; Nr. 15 156). Tissue blocks were cut into coronal sections of 30-µm thickness and collected into six series. Sections were collected into 24-well plates containing cryoprotectant (50% phosphate buffer, 20% glycerol, and 30% ethylene glycol) and stored at –20°C until further processing.

### 2.5 Tissue processing and immunohistochemical detection of tracers

Sections were removed from cryoprotectant and placed in 0.05 M Tris-buffered saline (TBS; pH 7.4 at room temperature) for five washes, each for five min (5 × 5). Endogenous peroxidase activity was suppressed by incubating sections in a TBS solution containing 0.014% phenylhydrazine for 20 min and then rinsed in TBS (5 × 5). Following a 2-h incubation in blocking solution consisting of 2% (v/v) normal donkey serum (MilliporeSigma, Burlington, Massachusetts, USA; Catalog #S30-100ML) and 0.1% Triton X-100 (Sigma-Aldrich, Inc., St. Louis, Missouri, USA; Catalog #T8532-500ML), sections were transferred into primary antiserum for about 48 hours at 4°C. Primary antisera contained antibodies raised against PHAL (species: rabbit; dilution: 1:4,000; Vector Labs, Cat # AS-2224); RRID: AB_2313686) or CTB (species: goat; dilution: 1:10,000; List Biological, Cat # 703; RRID: AB_10013220). Sections were then washed (5 × 5) in TBS and incubated in secondary antibody solution with biotinylated antibodies raised against rabbit (species: donkey; dilution: 1:1,000; Jackson ImmunoResearch Laboratories, Inc., West Grove, Pennsylvania, USA; Cat# 711-065-152; RRID: AB_2340593) or goat (species: donkey; dilution: 1:1,000; Jackson ImmunoResearch Labs, Cat# 705-065-147; RRID:AB_2340397). After a 5-h incubation period at room temperature and washes in TBS (5 × 5), signal amplification was achieved using an avidin-biotin-horseradish peroxidase complex (Vectastain ABC HRP Kit, Vector Labs, 45 µl reagent A and B per 10 ml) in 0.05 M TBS containing 0.1% Triton X-100 (v/v) for 1 h. Reacted tissues were then developed in 0.05% 3,3’-diaminobenzidine (DAB) (Sigma-Aldrich) mixed with 0.015% H_2_O_2_ in 0.05 M TBS for 10–20 min. Sections were then washed in TBS (5 × 5), mounted onto gelatin-coated slides and left to dry overnight at room temperature. Finally, tissue sections were dehydrated with ascending concentrations of ethanol (50–100%), defatted in xylene, and coverslipped with DPX mounting medium (Catalog # 06522; Sigma-Aldrich).

The same overall steps from DAB were used for immunofluorescent labeling. The phenylhydrazine reaction was omitted and secondary antibodies with fluorescent conjugates were used (see **Table 1**). After washing off secondary antiserum, freely-floating sections were mounted onto glass slides and coverslipped in sodium phosphate-buffered glycerol (0.05 M in 50% glycerol).

Dual-label immunohistochemistry was performed using the identical steps as described above for immunoperoxidase staining. Steps and reagents used for nickel-intensified DAB stain were identical to that described above except for the DAB solution. Nickel-intensification was achieved with 0.05% DAB mixed with 0.005% H_2_O_2_, and 0.1% ammonium nickel(II) sulfate dissolved in 0.05 M TBS. The phenylhydrazine step was omitted for the second stain that ended with a brown DAB product.

### 2.6 Antibody validation

Antisera for this study only contained those raised against PHAL and CTB. Specific staining was not observed for PHAL or CTB in brain tissues of animals that did not receive tracer co-injections. Additionally, specific staining was not evident for injection cases that did not effectively transport tracers. The same validation approach and outcome are found in other reports that catalogued the same antibodies (Hahn and Swanson, 2010; 2012).

### 2.7 Nissl staining

Sections were first washed (5 × 5) in 0.05 M TBS (pH 7.4 at room temperature) to remove cryoprotectant. Free-floating sections were mounted onto gelatin-coated glass slides and dried overnight at 60°C. Sections were dehydrated through ascending concentrations of ethanol (50%, 70%, 95%, and 100%) and defatted in xylenes for 25 min. Next, they were rehydrated and stained with a 0.25% thionine solution (thionin acetate, Catalog #T7029; Sigma-Aldrich) and differentiated in 0.4% anhydrous glacial acetic acid. Stained slides were dehydrated again and coverslipped with DPX mounting medium and left to dry overnight.

### 2.8 Photography and post-acquisition image processing

Immunostained and Nissl-stained sections were visualized and photographed under brightfield and darkfield microscopy using a BX63 microscope (Olympus Corporation, Shinjuku, Japan) equipped with a DP74 color CMOS camera (cooled, 20.8 MP pixel-shift, 60 fps). Image acquisition, stitching (15% overlap), and .TIFF image exporting were carried out with cellSens Dimension software (Version 2.3; Olympus). Images were adjusted for brightness and contrast using Adobe® Photoshop® (Version 13.0.1; Adobe Systems, Inc., San Jose, California, USA) and exported to Adobe® Illustrator® (<u>Ai</u>; Version CC 18.0.0; Adobe) for parcellation.

CNG sections stained for immunofluorescence were visualized with epifluorescence illumination using a Zeiss M2 AxioImager microscope equipped with an X-Y-Z motorized stage (Carl Zeiss Corporation, Thornwood, New York, USA) and a 100 W halogen light source. An EXi Blue monochrome camera (Teledyne QImaging, Inc., Surrey, British Columbia, Canada) operated by Volocity Software (Version 6.1.1; Quorum Technologies, Puslinch, Ontario, Canada; installed on an Apple Mac Pro computer) was used to generate widefield mosaic images from multiple channels. The same equipment was used to generate brightfield stitched images of adjacent Nissl-stained CNG-containing sections. Images were adjusted for brightness and contrast using Adobe® Photoshop® (Version 13.0.1; Adobe) and exported to Adobe® Illustrator® (Version CC 18.0.0; Adobe) for parcellation.

**Table 1.** List of antibodies and reagents used in this study.

| Reagent | Antigen/Conjugate | Host | Type | Source | Catalog # | Titer | Incubation |
| --- | --- | --- | --- | --- | --- | --- | --- |
| Primary | anti-PHAL | Rb | poly IgG | Vector Labs | AS-2300 | 1:4,000 | 60 h, 4°C |
|  | anti-CTB | Gt | poly IgG | List Biological | 703 | 1:10,000 | 60 h, 4°C |
| Secondary | anti-rabbit IgG | Dk | biotinylated | Jackson | 711-065-152 | 1:500 | 5 h, 22°C |
|  | anti-goat IgG | Dk | biotinylated | Jackson | 705-065-152 | 1:500 | 5 h, 22°C |
|  | anti-rabbit IgG | Dk | Cy5-conjugated | Jackson | 711-175-152 | 1:500 | 5 h, 22°C |
|  | anti-goat IgG | Dk | Cy3-conjugated | Jackson | 705-165-147 | 1:500 | 5 h, 22°C |
| Conjugate | avidin | – | HRP-conjugated | Vector Labs | PK-6100 | na | 1 h, 22°C |

### 2.9 Mapping of immunohistochemically detected tracer

The atlas-based mapping approach used here follows the cytoarchitectonic approach described by Swanson (1992, 1998, 2004, 2018). Briefly, formal cytoarchitecture-based definitions for gray matter regions were obtained for all areas examined (see *Table C* in *Supplementary Information Folder 1* from Swanson [2018]), and boundaries were drawn with Nissl-stained reference sections and then superimposed on DAB-stained sections to localize tracers. One of two approaches were used for this process. The first involved making camera lucida pencil-on-paper drawings of cytoarchitectonic boundaries via a drawing tube fitted to an Olympus microscope, aligning drawn boundaries to corresponding immunostained sections, and then transcribing histological data to digital atlas templates. The second approach involved placing images of Nissl- and immunostained tissue sections onto <u>Ai</u> files as separate layers. Immunostained images were aligned to Nissl and boundary layers by relying on shared features (*i.e.*, brain surfaces, white matter tracts, background staining, and blood vessels). These overlays informed the final transcription of data onto atlas plates on a different <u>Ai</u> file. It should be emphasized that the transcription process, in our case, was a representation and never an attempt to simply fit or copy-paste information. Matching histological sections to atlas levels involved careful scrutiny of the histological plane-of-section and other forms of non-linear distortion (Simmons and Swanson, 2009a). Once the corresponding atlas levels were identified, maps were created using data from only the best-matched sections. Once histological sections were matched to atlas levels, axons were drawn using the *Pencil* tool in <u>Ai</u> with high accuracy settings and cell bodies were plotted as ellipse objects. The goal was to accurately show spatial information with an attempt to also capture some morphological features such as connectional density, direction, and axonal morphology.

### 2.10 Semi-quantitative analysis

Mapped CTB datasets were used as a starting point to obtain a 0–5 scaling system for connectional strength. CTB-ir neuron counts across each diencephalic region at each atlas level were tabulated within .CSV files. First, data were transformed using the natural logarithm and histograms were generated to confirm linearity. The highest log value was used to divide the dataset into quintiles, and the log values for each division were used to calculate thresholds for CTB connectional scores. Numerical thresholds for each score were calculated separately for each experiment (*see* **Supplemental Figure 1**). To score PHAL axonal density, photomicrographs were examined in bins of 128 × 128 pixels (roughly 120 μm ×120 μm). **Supplemental Figure 1** shows representative snapshots for each 0–5 score category. Scores were determined by obtaining bins from each diencephalic region and comparing them to the representative snapshots.

**Figure 1.**
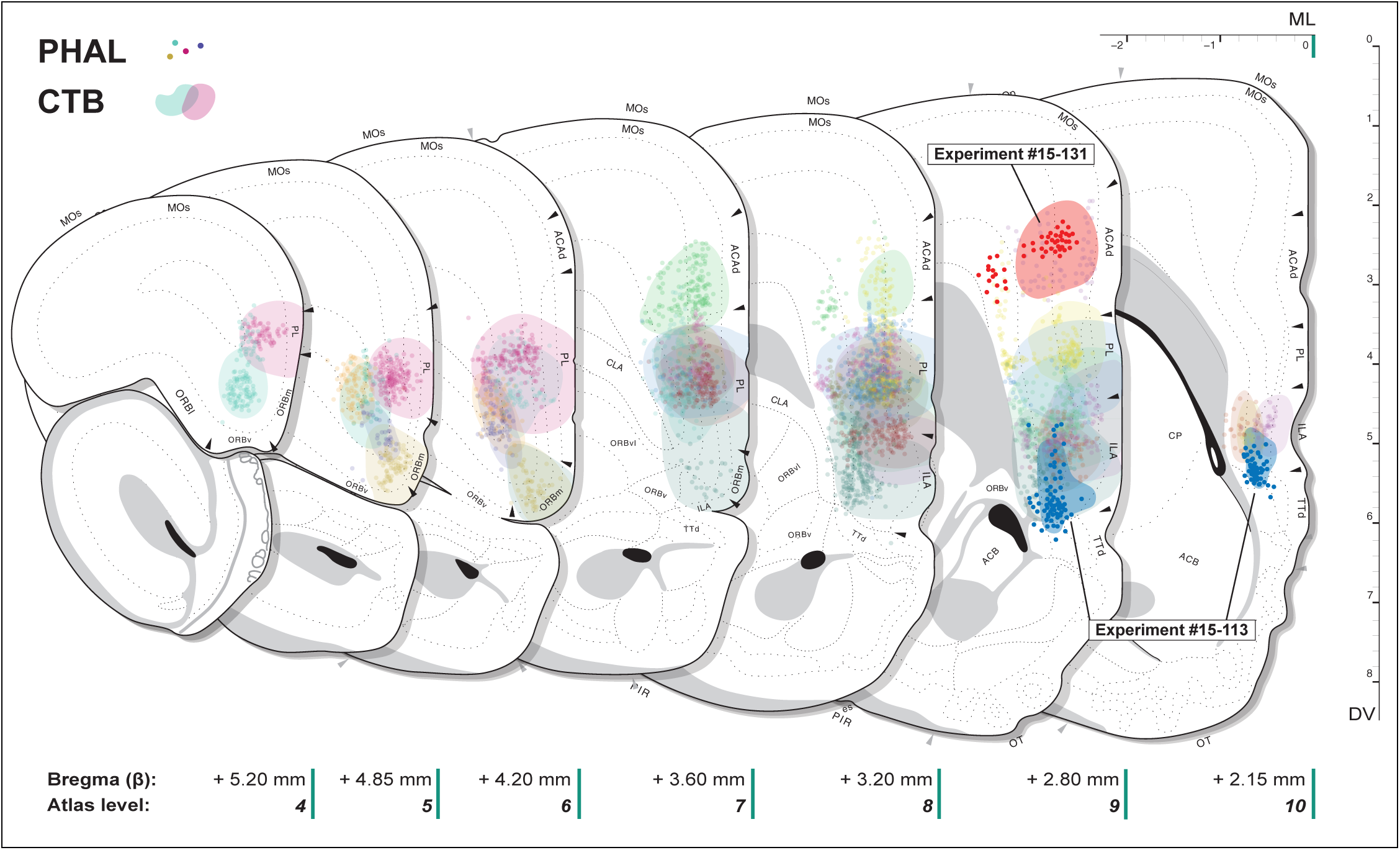
Maps showing PHAL and CTB injection sites in the *cingulate region (Brodmann, 1909)* (CNG). Each experiment is coded with a unique color, with PHAL-ir cells shown as circles and CTB injection spread represented with contours. The scales on the edges indicate mediolateral (*top*), dorsoventral (*right*), and anteroposterior (*bottom*) dimensions based on atlas levels from *Brain Maps 4.0* (*BM4.0*; Swanson, 2018). See the list of abbreviations on page 35 for their explanation.

## 3. Results

### 3.1: Neuroanatomical connections of the CNG

Immunodetected sites for PHAL and CTB injections into the CNG were localized with the aid of an adjacent Nissl-stained reference section. Injection site maps show the extent of coverage in our study (**Figure 1**). Cytoarchitectonic parceling of the CNG followed descriptions (Krettek and Price, 1977; Vogt and Peters, 1981) found in the annotated nomenclature tables for *Brain Maps 4.0* (*BM4.0*; Swanson, 2018). Although delivered from the same micropipettes, co-injected PHAL and CTB were treated as individual experiments because they differ in their diffusion properties and uptake mechanisms (Gerfen and Sawchenko, 1984; Luppi et al., 1990). We found that co-injected tracers largely overlapped in their distributions but marginal differences, such as a typically wider spread for CTB, were observed. From a total of 39 co-injection experiments, five were selected to investigate CNG connectional distributions within the diencephalon. These include two injections centered in the ACAd and three injections centered in the ILA (**Figure 1**). One of our mapped experiments, #15-113, is located in a ventral part of the ILA at atlas levels 9 and 10 in *BM4.0* (**Figure 2**). This roughly corresponds with the so-called “dorsal peduncular cortex” (Paxinos and Watson, 2014; Akhter et al., 2014), which has a slightly cell-sparse layer 2/3. PHAL-filled cell bodies were most abundant in layer 5 and adjacent layers. CTB spread was also concentrated in layer 5 but showed less immunoreactivity in layer 6. The ACAd injection (experiment #15-131) was restricted to level 9 of *BM4.0*; PHAL-immunoreactive (-ir) cells were concentrated in the boundary of layers 3 and 5, with a separate cluster in layer 6. CTB had a circular spread contained mainly inside layers 3 and 5.

Our analysis of CNG connectivity focused on the entire *interbrain (Baer, 1837)* (diencephalon). Connectivity maps showing tracer distributions were drawn from level 16 to level 34 (approx. 4.5 mm, ranging between +0.10 mm and −4.45 mm from bregma; **Figure 3**). Our data were also summarized using semi-quantitative scores (**Table 2**) to facilitate the assembly of connectomes. For simplicity, our account of efferent and afferent ILA and ACAd connections will proceed across major divisions of *thalamus (His, 1893a)* (TH) and *hypothalamus (Kuhlenbeck, 1927)* (HY). And, given that CNG connections have been characterized by many others (see **Discussion**), we will privilege the novel findings that result from our atlas-resolution mapping effort and which display bidirectional connectivity.

**Figure 2.**
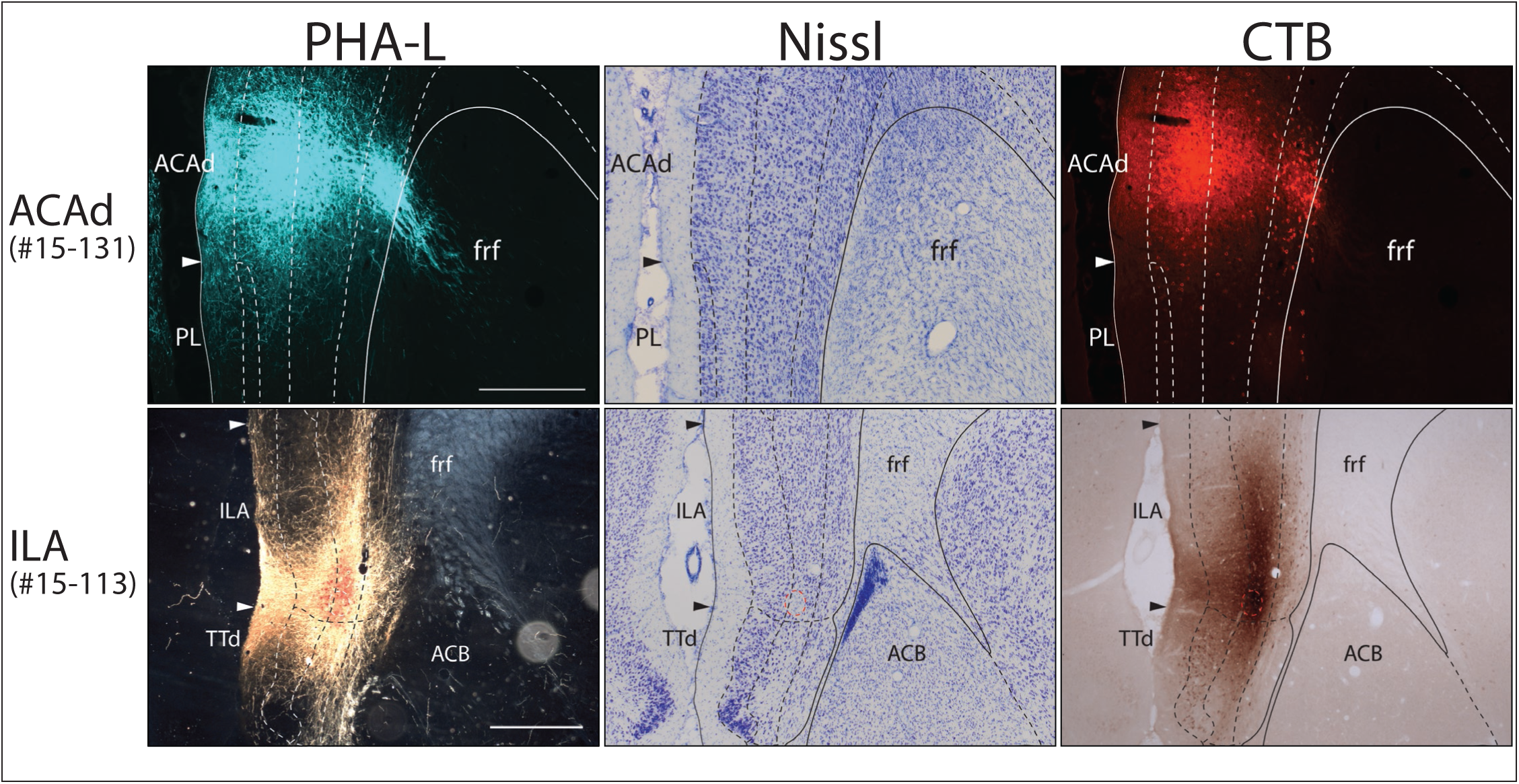
Photomicrographs showing the centers of PHAL and CTB injections into the ILA and ACAd. Regional boundaries derived from adjacent Nissl-stained sections (*middle column*) were superimposed on immunostained images of PHAL (*left column*) and CTB (*right column*). Immunostained images shown here for ACAd and ILA were visualized with epifluorescence and DAB reactions, respectively. Scale bars: 500 μm. See the list of abbreviations on p. 35 for an explanation of those shown in this figure.

### 3.2: Description of fiber systems used by the ILA and ACAd to enter the diencephalon (PHAL anterograde tracing)

ILA axons innervating the diencephalon arrive through two routes. The primary route involves axons forming a lateral segment of the *medial forebrain bundle (Edinger, 1893)* (mfb) (**Figure 3**). This fiber system appears to be the origin of all ILA axons that are found in the HY (Negishi and Khan, 2019). In more caudal parts of HY, a substantial group of mfb collaterals ascend dorsomedially to target caudal parts of TH and the *periaqueductal gray (>1840)* (PAG) of the *midbrain (Baer, 1837)* (**Fig. 3n–s**). ILA axons in the mfb continued as far as the *cerebral peduncles (Tarin, 1753)*. The second route involves corticofugal axons that form fascicles in a ventromedial part of the *caudoputamen (Heimer & Wilson, 1975)* (CP). This group, in our samples, appeared to enter the *terminal stria (Wenzel & Wenzel, 1812)* (st; “stria terminalis”) to innervate the *bed nuclei of terminal stria (Gurdjian, 1925)* (BST) (**Fig. 3n–s**). It is not clear if this route contributes ILA axons to rostral TH. Instead, ILA axons were often noted exiting the mfb in the direction of TH.

In contrast, ACAd axons formed substantial groupings of passing fibers that were observed in the *internal capsule (Burdach, 1822)* (int) until they exited as thalamic radiations through the *reticular thalamic nucleus (>1840)* (RT) at about −1.33 mm from bregma (**Fig. 3h–k**). A subset of ACAd axons continued through the int until exiting through the ZI to arrive at ventral TH (**Fig. 3m**). More caudally, ACAd axons exited the int entirely to innervate the caudal hypothalamus and PAG (**Fig. 3r, s**).

### 3.3: Projections from ILA and ACAd to hypothalamus (PHAL anterograde tracing)

We observed that although ILA axons were present throughout the hypothalamus, they were concentrated in a few regions. In more rostral sections, ILA axons in the *anterior region (Swanson, 2004)* of LHA (LHAa) took the form of passing fibers in the mfb (**Fig. 3i–k**). Once LHAa transitioned to the *dorsal region (Swanson, 2004)* (LHAd), massive collaterals were observed in the amygdala-bound *peduncular loop (Gratiolet, 1857)* (pdl; “ansa peduncularis”; [see Leuret & Gratiolet, 1839–1857]) and towards the LHAd and *suprafornical region (Swanson, 2004)* (LHAs) (**Fig. 3l–n** and **Figure 4**). This collateral system continued until the appearance of the *subthalamic nucleus (>1840)* (STN) (**Fig. 3n, o**). These collaterals occupied an anterior-posterior distance between approximately −2.00 mm to −2.85 mm from bregma. It is important to note that the LHAa and LHAd, as Nissl-defined regions, are distinguished on the basis of cell density (Swanson, 2018). Here, we observed that the cell-sparse LHAd is cospatial with a sizable increase in the density of axon collaterals. In more caudal tissue sections, another major terminal zone was observed in the *parasubthalamic nucleus (Wang & Zhang, 1995)* (PSTN). ILA terminals to PSTN were remarkably restricted within its cytoarchitectonic boundaries (**Fig. 3q, r**). A subset of mfb collaterals formed dense terminals in the cytoarchitectonically distinct *terete part (Petrovich et al., 2001)* (TUte) of the *tuberal nucleus (>1840)* (TU) (**Fig. 5a, b**); this terminal field formed a compact tube shape at the base of the hypothalamus (**Fig. 3o–r**). By far the densest ILA projections were found in a horizontal band immediately ventral to the *posterior hypothalamic nucleus (>1840)* (PH) and *supramammillary nucleus (>1840)* (SUM) (**Fig. 3r, s**). There is no clear cytoarchitecture related to this ILA terminal field, but it is likely the dorsal cap of the *medial mammillary nucleus (Gudden, 1881)* (MM) that includes its *median part (>1840)* (MMme) (**Fig. 5c, d**).

**Table 2.**
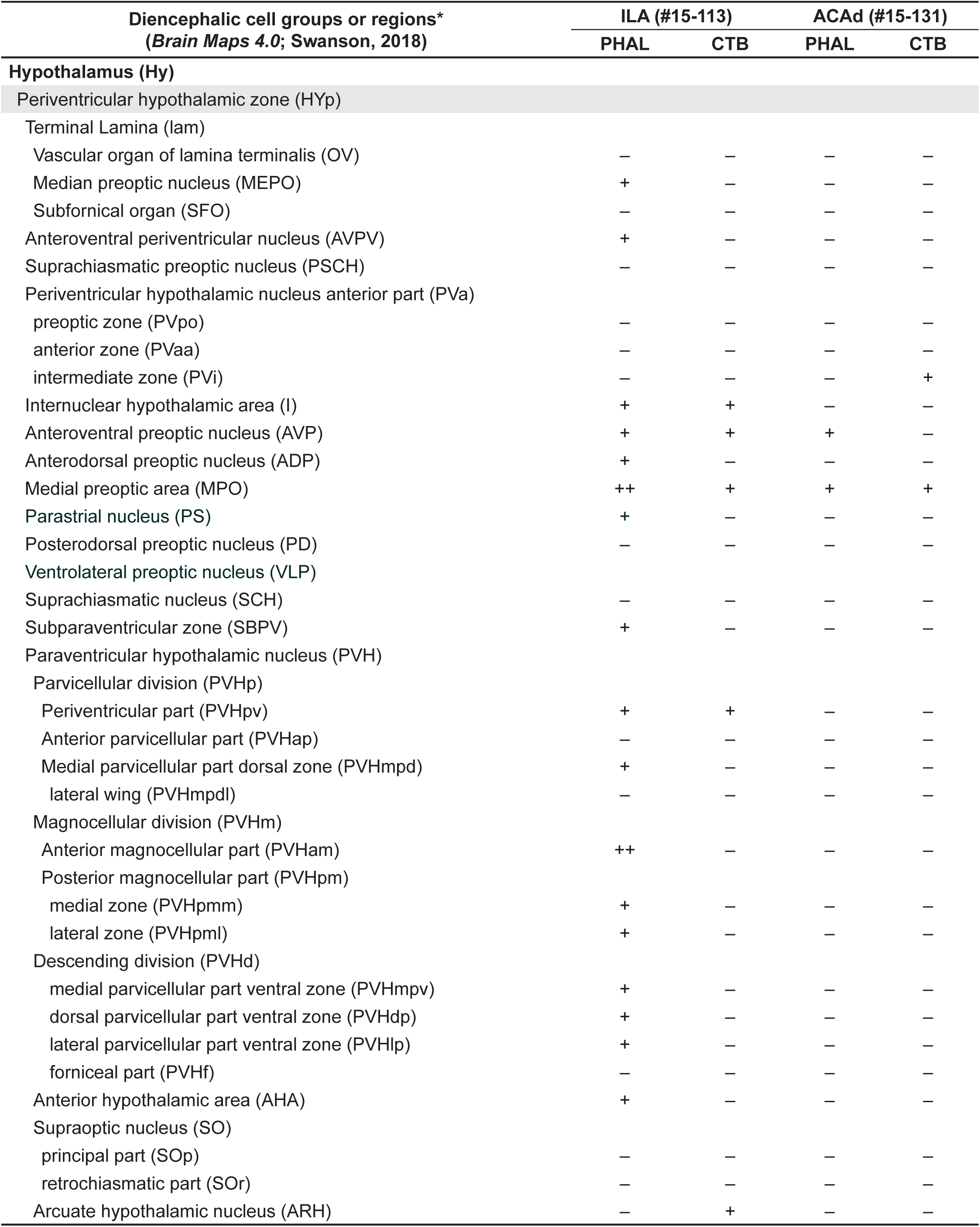
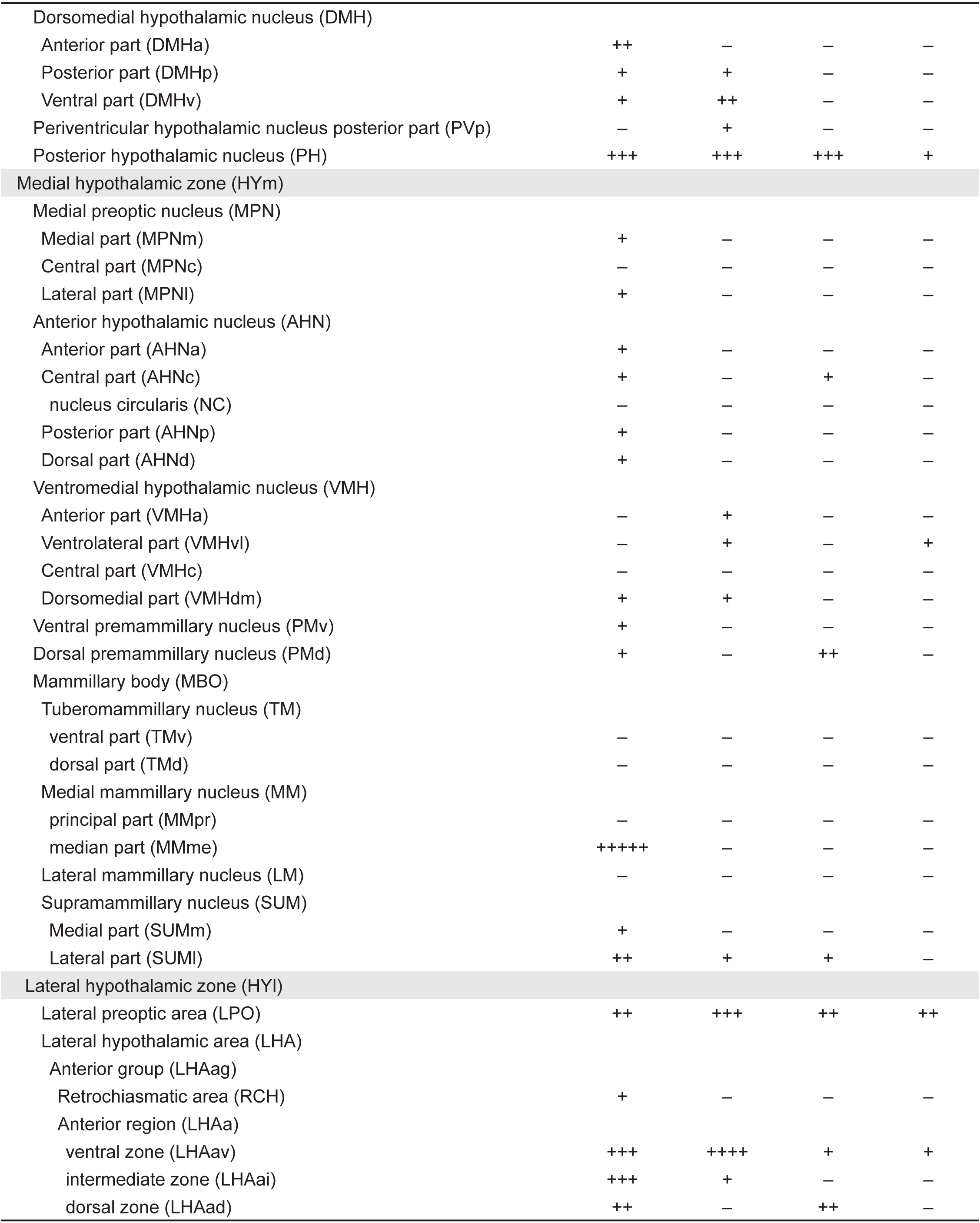
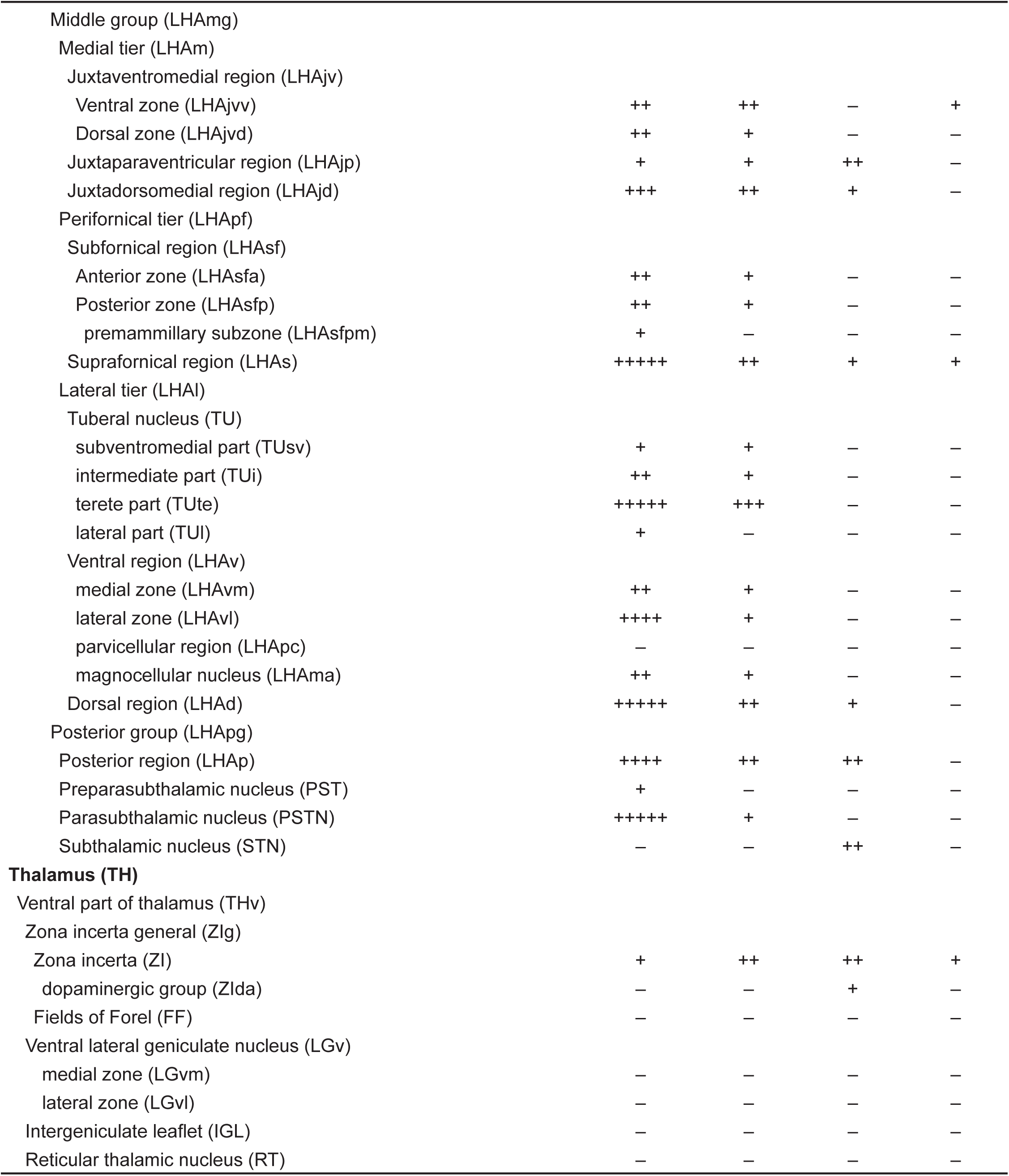
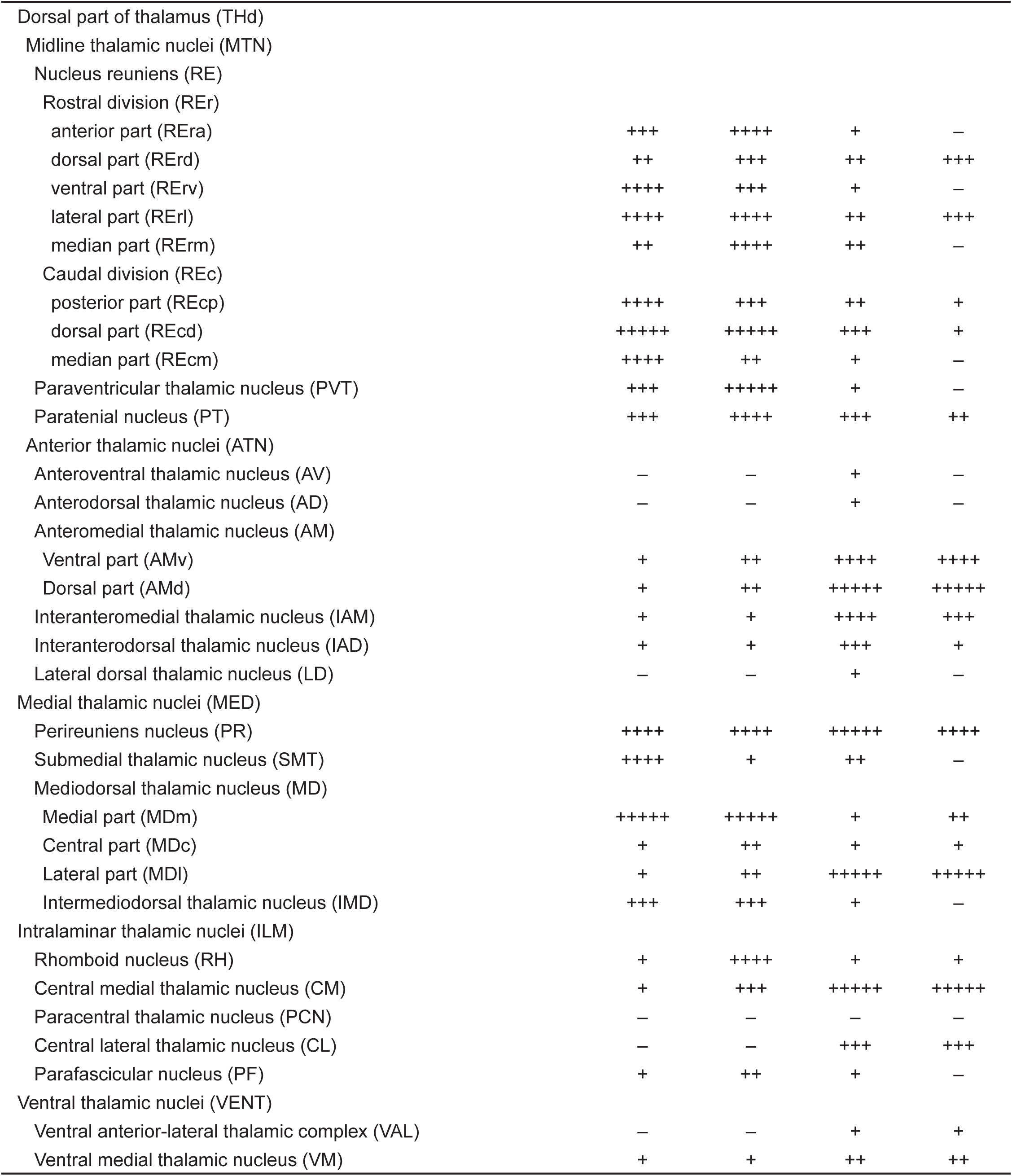
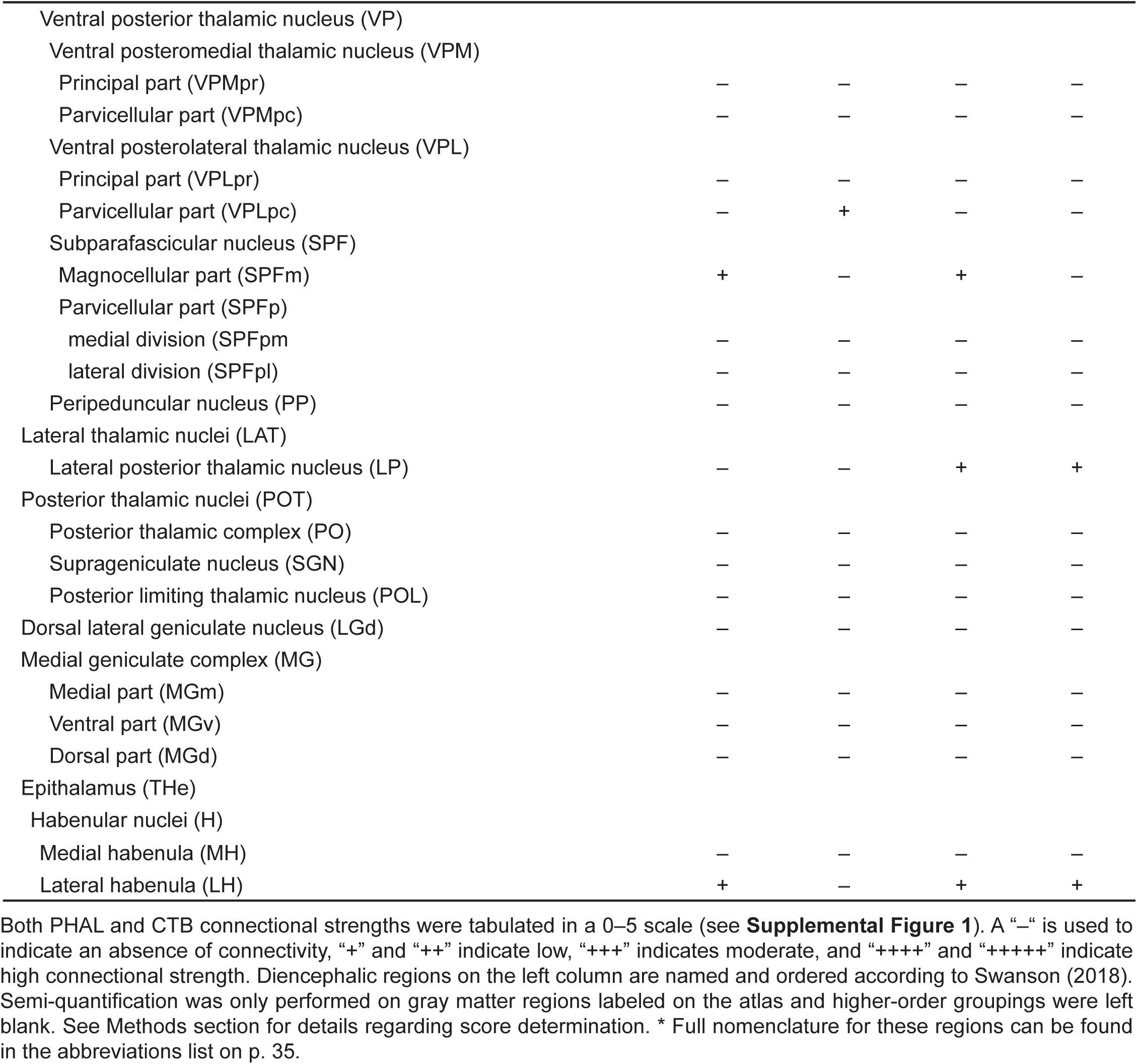
Semi-quantitative analysis of CNG connectional weights with *interbrain (Baer, 1837)* (diencephalon) by region.

**Figure 3.**
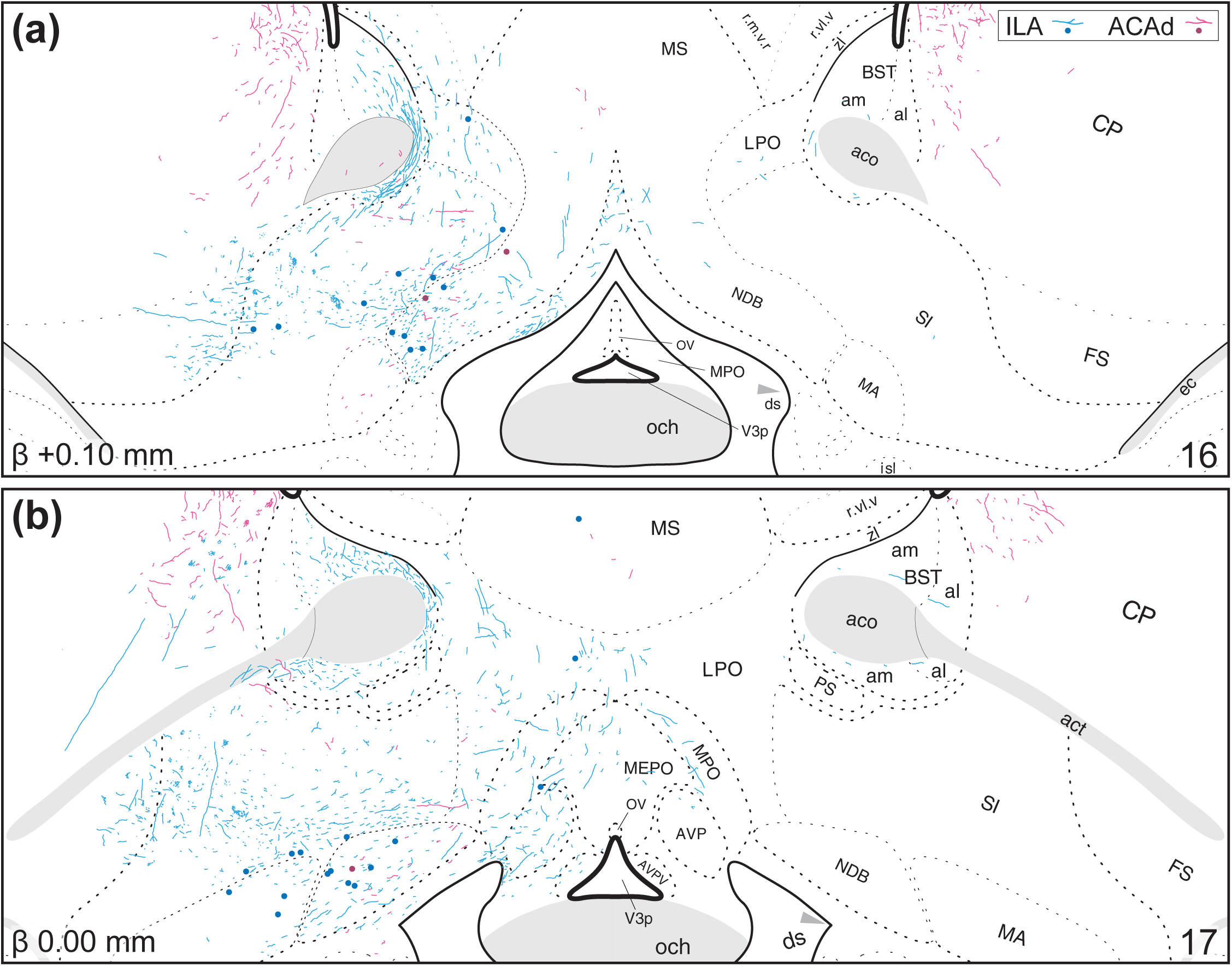
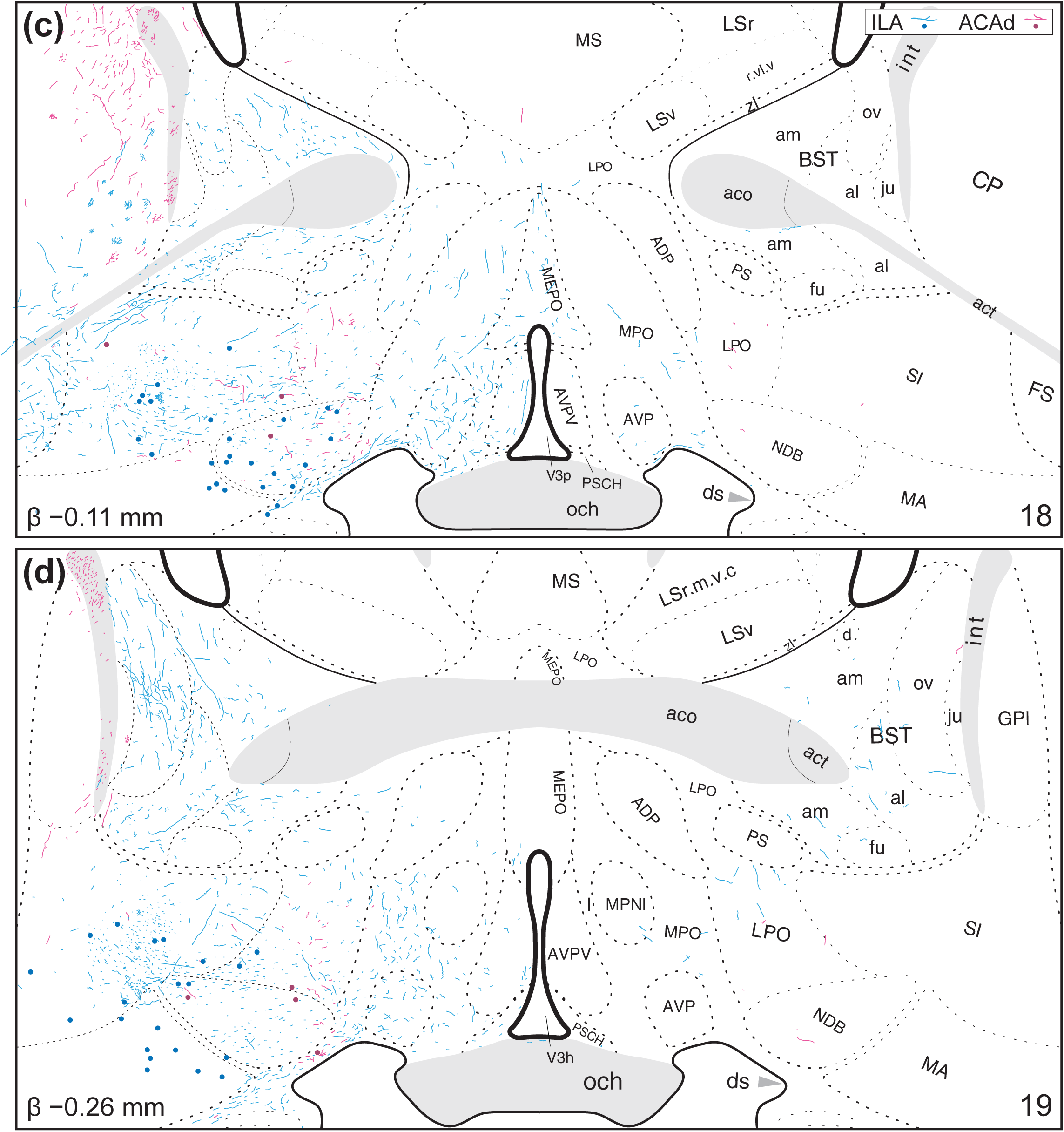
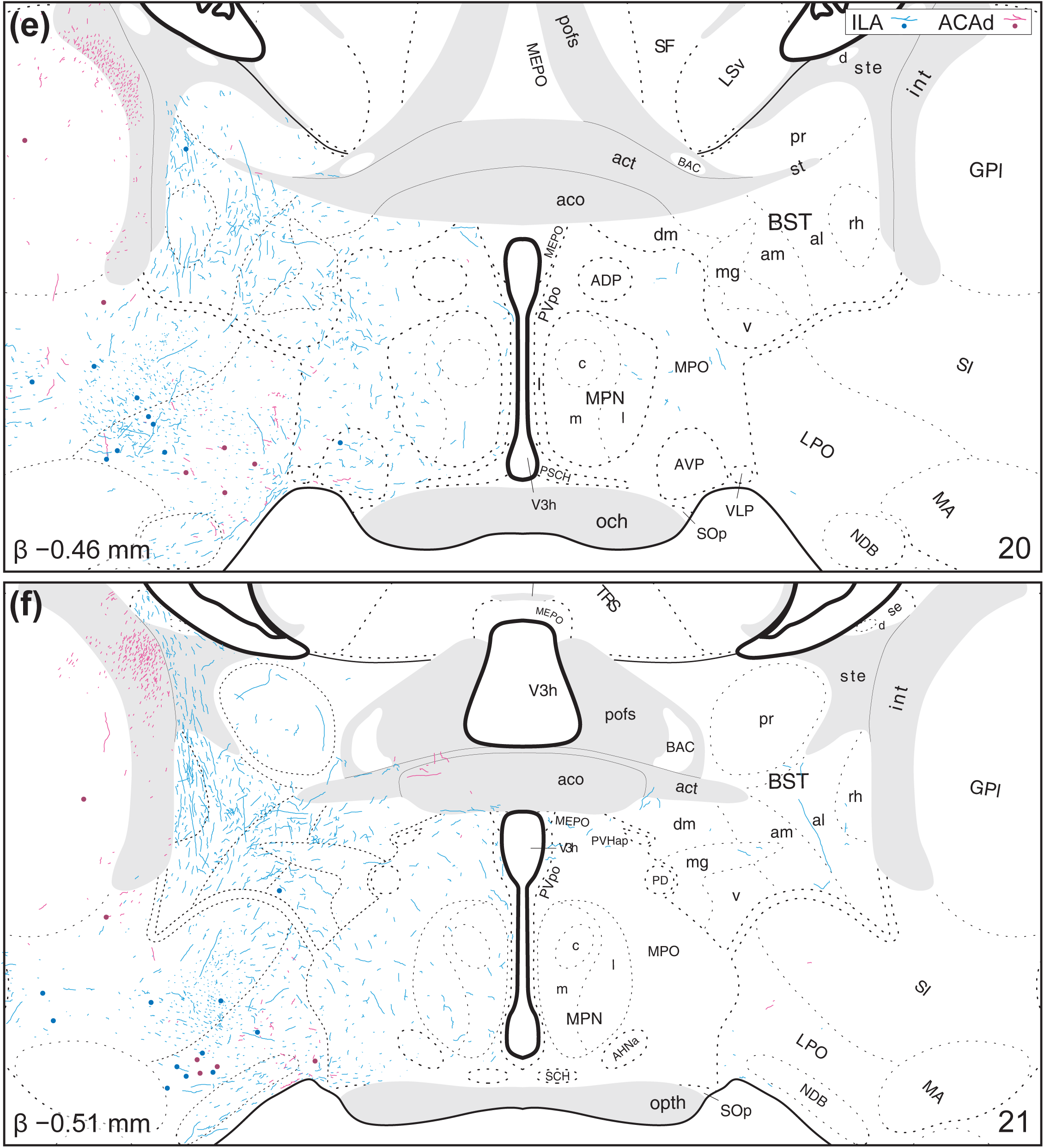
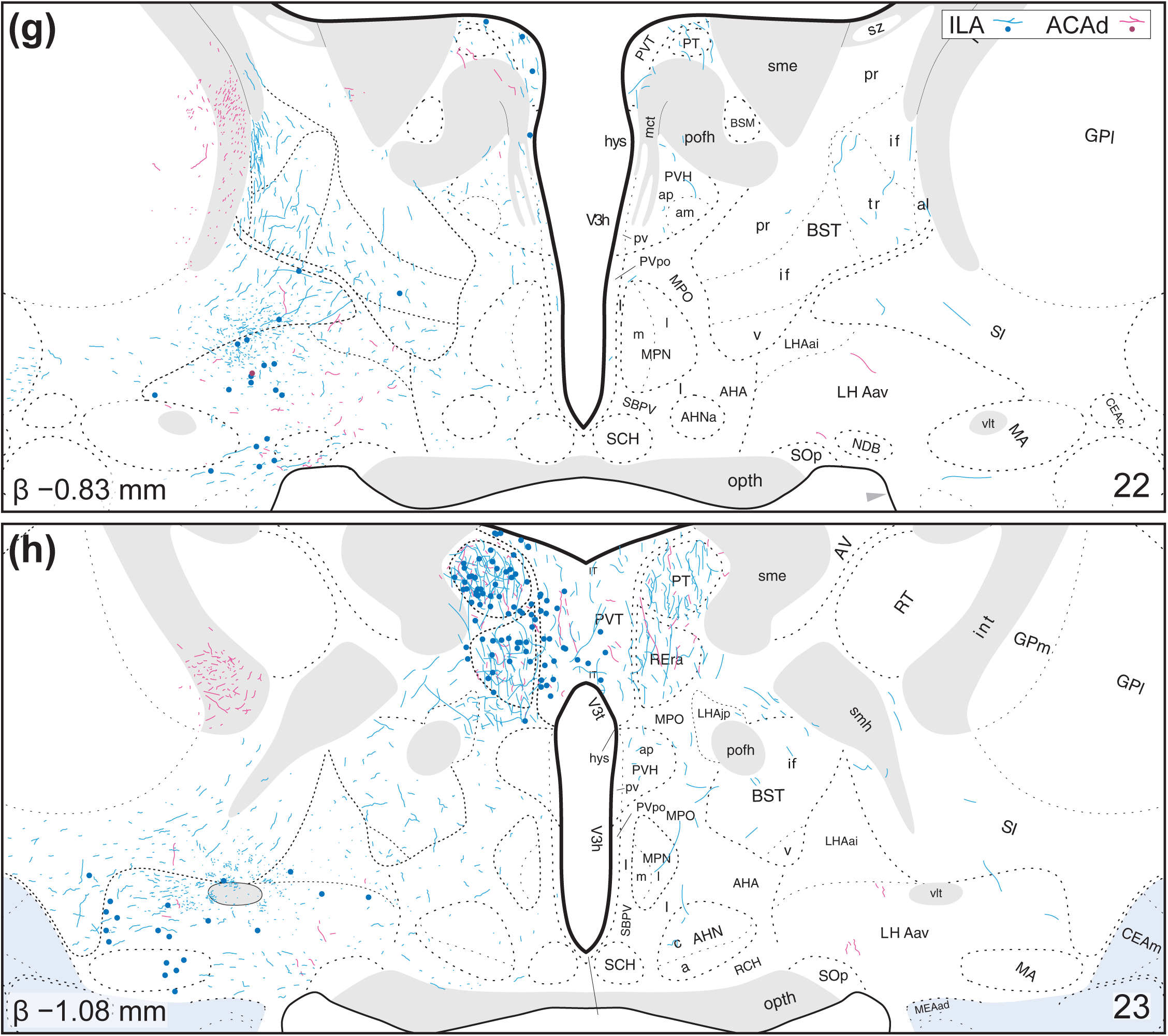
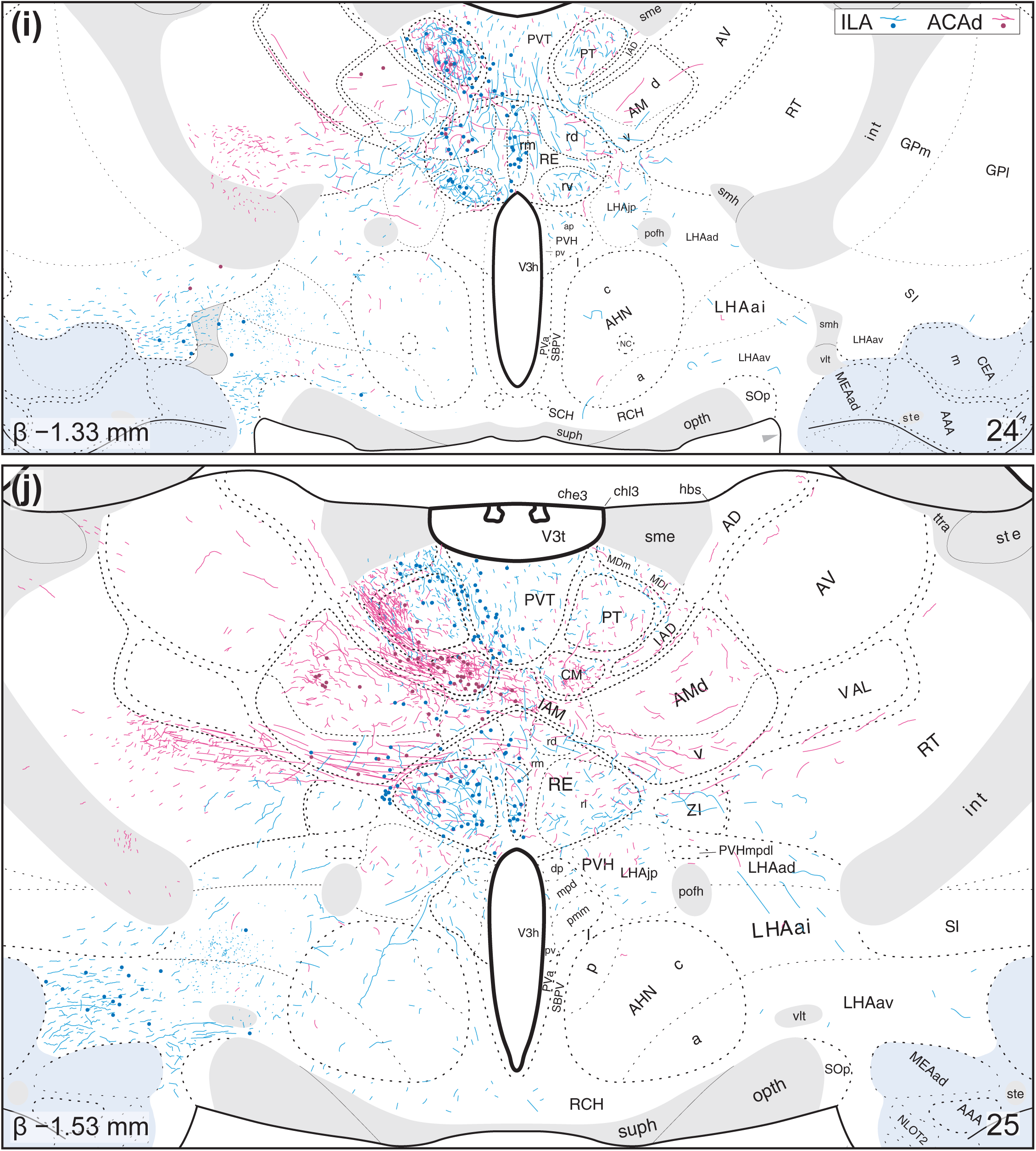
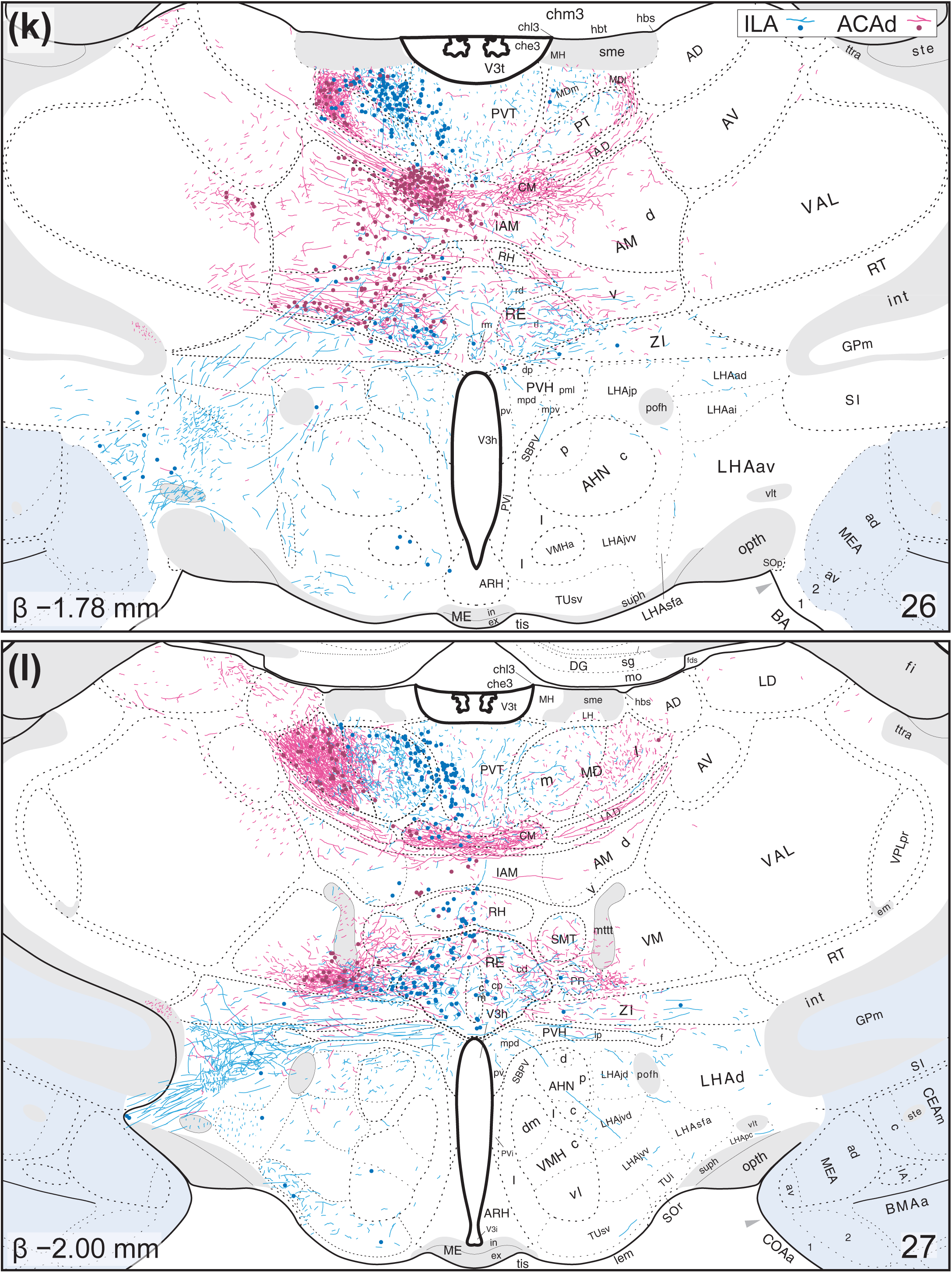
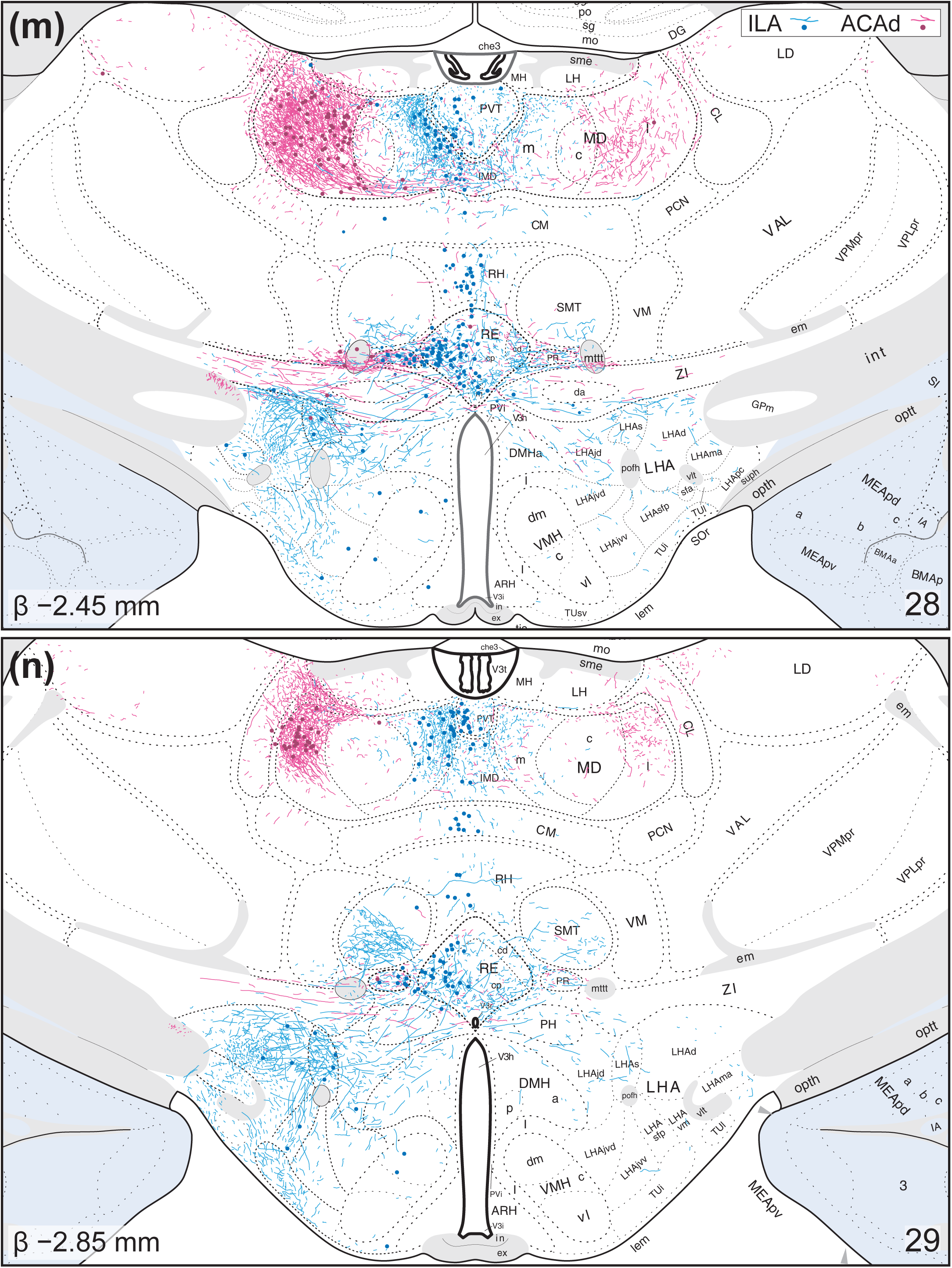
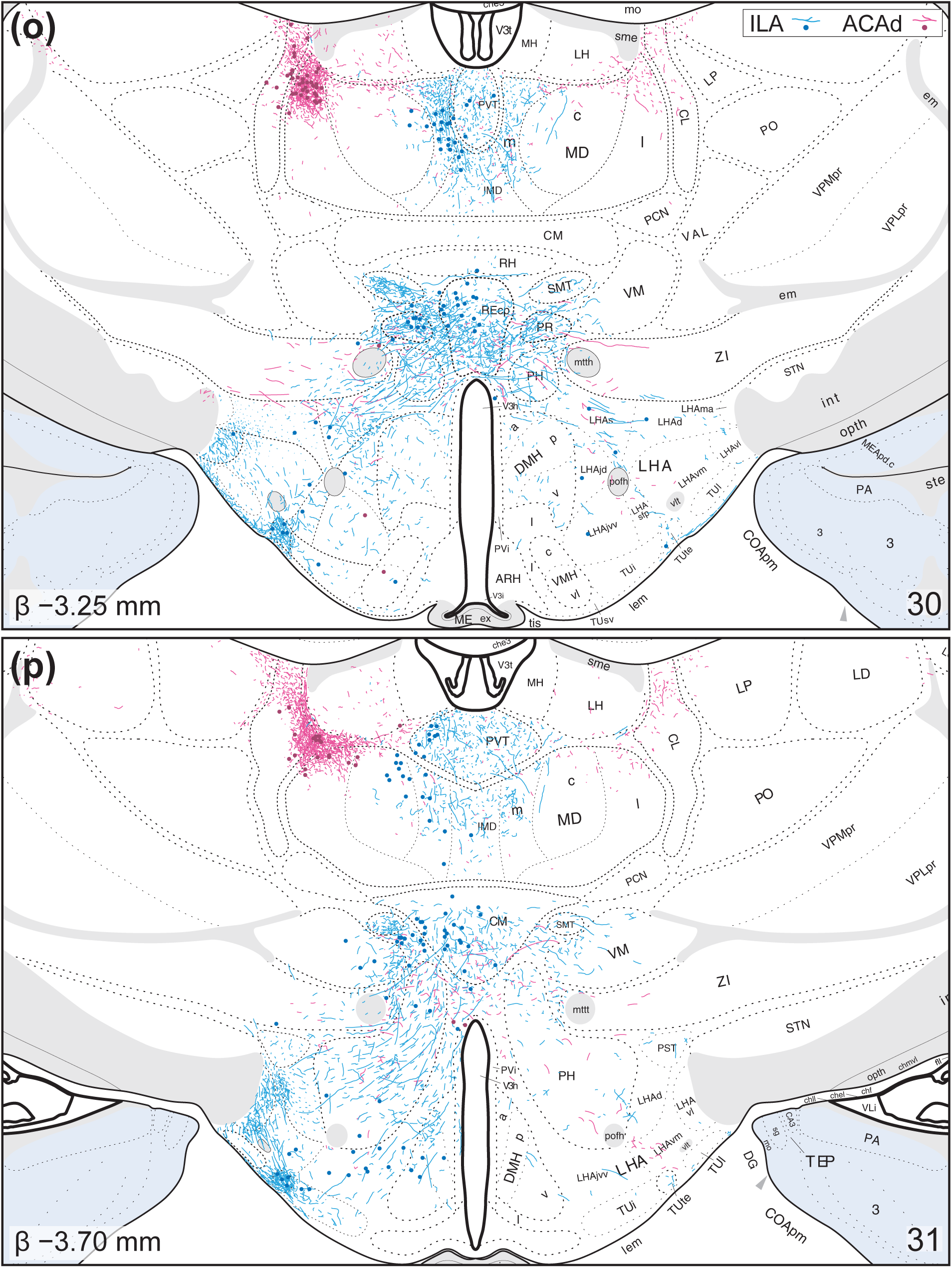
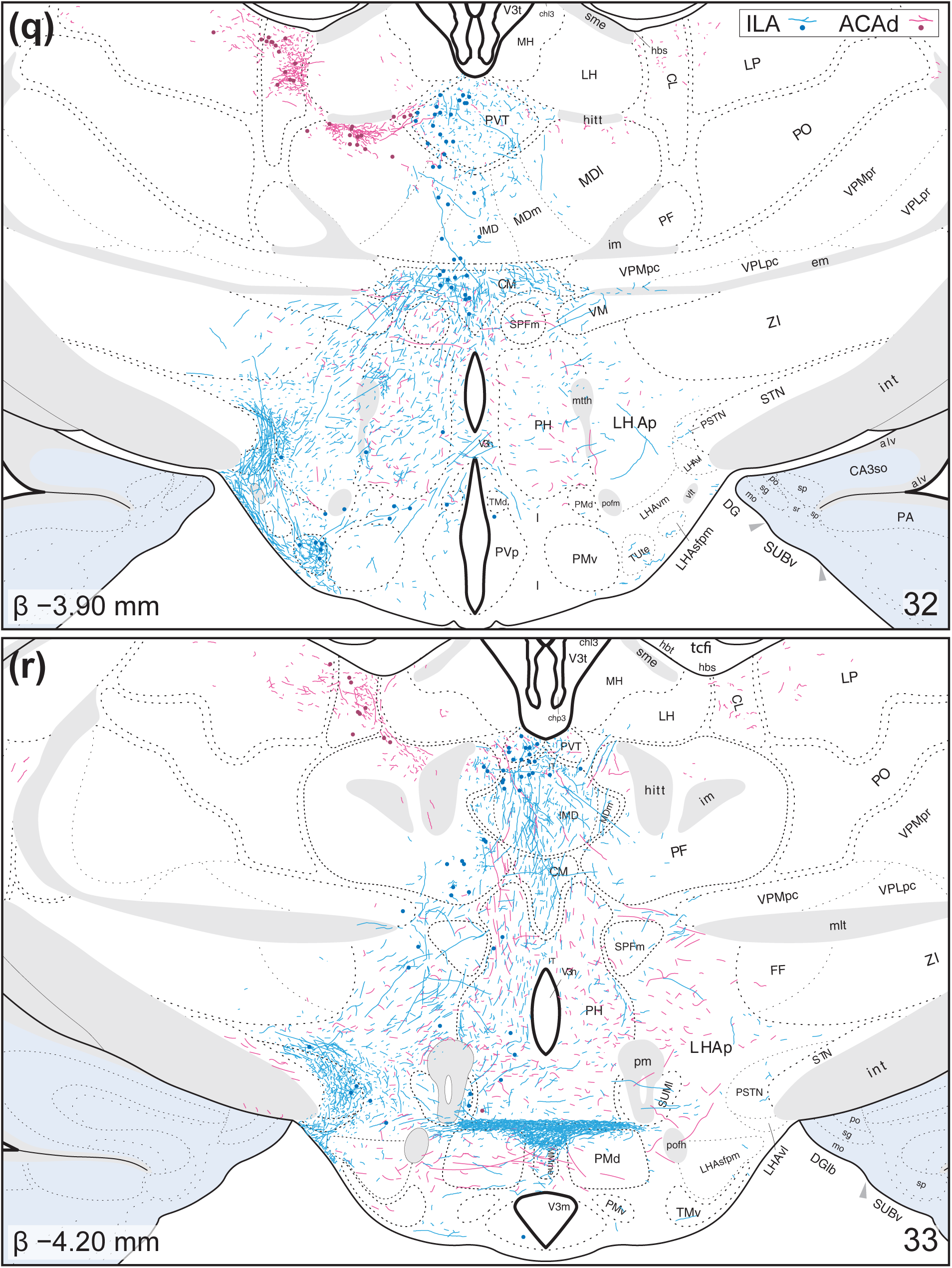
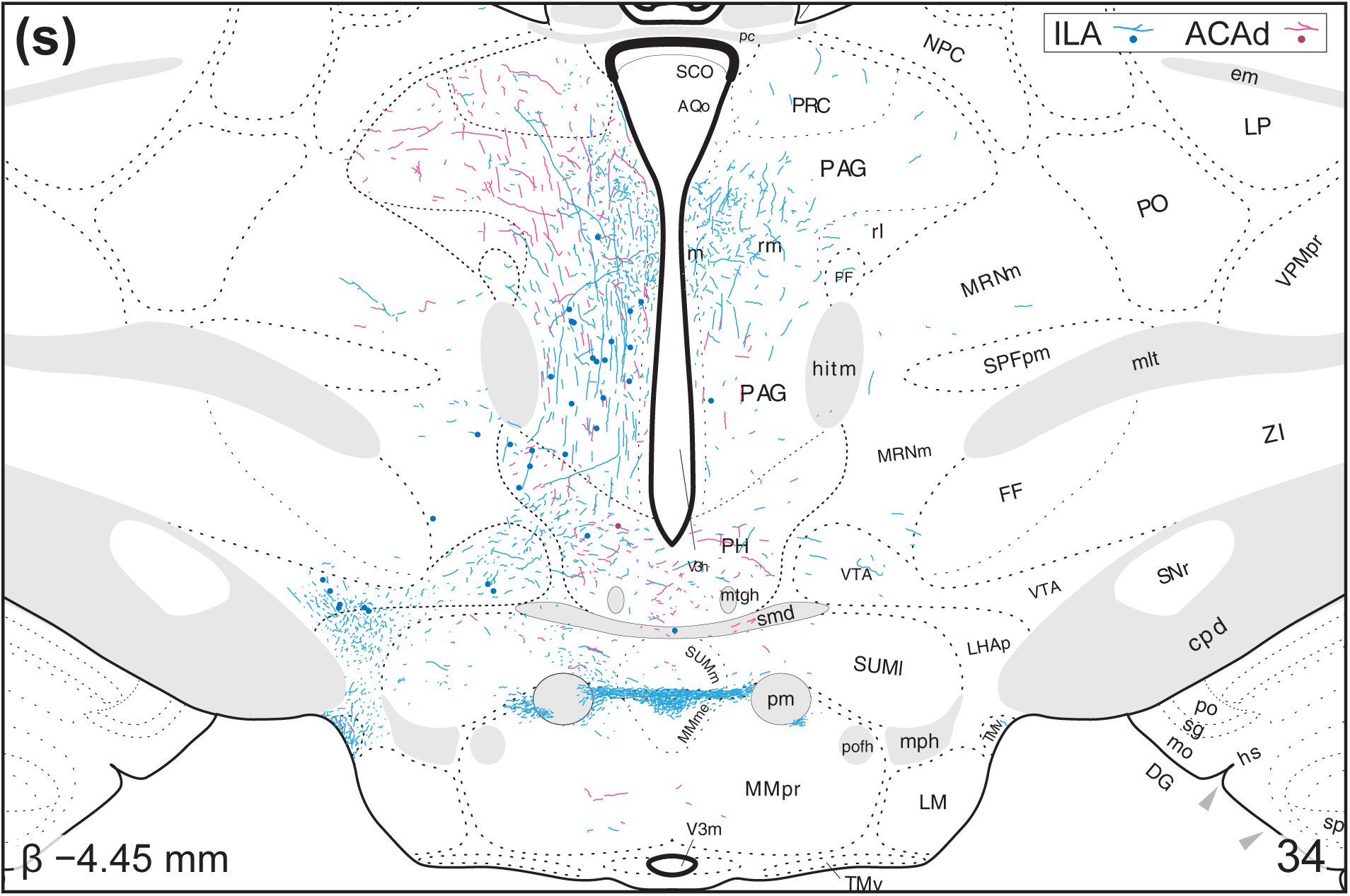
Representative maps showing distributions of immunoreactive PHAL and CTB throughout the diencephalon. Color-coded tracers from ACAd (*purple*) and ILA (*blue*) were localized using boundaries deduced from Nissl-stained sections. Dots mark cell bodies, lines mark axonal fibers. Inferred anterior-posterior positions, indicated as bregma values in the lower-left corners, were derived from *BM4.0* (Swanson, 2018). Atlas levels from *BM4.0* are indicated in the lower-right corners. See the list of abbreviations on p. 35 for an explanation of the abbreviations shown in this figure.

ACAd axons in hypothalamus were sparse. However, a notable increase in axon densities was apparent in the PH (**Fig. 3q–s**). ACAd axons were also present in the *subthalamic nucleus (>1840)* (STN) and *dorsal premammillary nucleus (>1840)* (PMd), regions that were avoided by ILA axons.

### 3.4: Hypothalamic afferents to the ILA and ACAd (CTB retrograde tracing)

CTB-labeled neurons in hypothalamus were far fewer than compared to thalamus. The LHAa and *lateral preoptic area (>1840)* (LPO) contained, by far, the most retrogradely labeled neurons from ILA (**Fig. 3d–k**). Though few in numbers, ILA-projecting neurons were always present in hypothalamic volumes that contained dense axon terminals. These included the TUte, LHAd, LHAs, and PSTN (**Fig. 3l–r**).

Retrograde labeling from the ACAd was virtually undetected in the hypothalamus. The LPO contained the most CTB-ir from the ACAd whereas the adjacent LHAa contained scant labeling (**Fig. 3d–h**).

### 3.5: Bidirectional ILA and ACAd connections with thalamus (distributions of PHAL and CTB)

Two general observations can be made regarding CNG connections with *thalamus (His, 1893a)* (TH). Connectivity between CNG and TH, with few exceptions, tended to form reciprocal connections. PHAL-ir axon terminals and CTB-ir neurons were tightly coupled in space, with the *submedial (>1840)* (SMT), *central medial (Rioch, 1929)* (CM), and *rhomboid (Cajal, 1903)* (RH) *thalamic nuclei* being the only regions that showed unidirectional connections (**Fig. 3h–r**). ILA and ACAd connections were segregated in TH. TH regions that connected with both ILA and ACAd, such as the *mediodorsal thalamic nucleus (>1840)* (MD), tended to occupy non-overlapping compartments (**Fig. 3h–r**). The *nucleus reuniens (Malone, 1910)* (RE) and *paratenial nucleus (>1840)* (PT) were the only regions in which ILA and ACAd connections overlapped (**Fig. 3i–l**).

Most CNG connections with TH, in both volume and density, were formed in the MD (**Fig. 3h–r**). The ILA and ACAd were respectively connected with the *medial (>1840)* (MDm) and *lateral (>1840)* (MDl) *parts* of MD, and both cortical areas appeared to skirt around the *central part (>1840)*. CNG connections in the caudal half of MD appeared to withdraw towards its dorsal boundary (**Fig. 3n–q**). In the most caudal parts of MD, the ACAd connections were densely aggregated in a space between the *central lateral thalamic nucleus (Rioch, 1929)* (CL) and MD, forming the ventrolateral perimeter of the *lateral habenula (>1840)* (LH) (**Fig. 3p–r**).

Dense bidirectional ACAd connectivity was observed in the CM, initially appearing as a small circular cluster beneath the PT (**Fig. 3i–l** and **Figure 6**). High-magnification revealed numerous instances of putative axo-somatic contacts between PHAL-ir axons and CTB-ir neurons (**Fig. 6c–f**). ACAd connections with CM abruptly ended as the SMT emerged (AP approx. −2.45 mm from bregma). The most caudal part of CM was bidirectionally connected with the ILA (**Fig. 3p–r**), and PHAL-ir axons in this segment were passing dorsally towards the paraventricular nucleus (PVT).

A ventral bidirectional cluster of ACAd connections was found in an ill-defined space immediately ventral to the *thalamic mammillothalamic tract (Swanson, 2015)* (mttt) (**Fig. 3l, m**). This cluster was initially thought to be in the ZI, but examination with an adjacent Nissl-stained section revealed a cell-sparse zone separating this cluster from the ZI (**Fig. 7a, b**). Moreover, examination at higher magnification revealed several instances of putative axo-somatic contacts between PHAL-ir axons and CTB-ir neurons (**Fig. 7c–e**). It is unclear whether these ACAd connections are with the *perireuniens nucleus (Brittain, 1988)* (PR) or the *ventral medial thalamic nucleus (>1840)* (VM) as it is centered between them. ACAd projections likely enter this cluster through the group of axons that pass through the RT and *ventral part (Canteras & Swanson, 1992)* (AMv) of the *anteromedial thalamic nucleus (>1840)* (AM). ILA connections did not contribute to this sub-mammillothalamic cluster, but bidirectional connectivity was observed in the adjacent PR and RE.

Moderate retrograde labeling from the ILA was present in every level of the *paraventricular thalamic nucleus (>1840)* (PVT). These connections were bidirectional, although ILA axons were slightly more concentrated in its caudal half (**Fig. 3Gg–r**). ILA and ACAd axons were present in the rostral half of PVT, but they were sparse and had the morphology of axons-of-passage (**Fig. 3g–l**). Taken together, the PVT appears to be largely unidirectional with respect to ILA connectivity. The AM and *interanteromedial nucleus (>1840)* (IAM) were predominantly connected with the ACAd (**Fig. 3i–l**). Some ILA projections were detected in the AMv, but our samples mostly contained retrograde labeling of AM.

There were a few examples of unidirectional connectivity among CNG connections with thalamus. ILA axons to the SMT were densely concentrated in its ventral and caudal halves (**Fig. 3l–p**). Retrograde labeling from ILA was almost absent in this structure. The RH, by contrast, mostly contained retrograde labeling from ILA whereas anterograde labeling was sparse (**Fig. 3k–o**). Finally, ACAd axons were observed in the ZI, primarily appearing to be fibers of passage (**Fig. 3k–o**). ZI afferents to ACAd, however, were rarely observed in our experiments. In contrast to this, ZI showed retrograde labeling from the ILA with little indication of PHAL-ir axon terminals.

## 4. Discussion

In this study, we co-injected anterograde and retrograde tracers to describe the structural organization of bidirectional macro-connections between the CNG and diencephalon. We produced, to our knowledge, the highest spatial resolution maps for ILA and ACAd connectivity to date. This enabled a precise mapping of connectional topography and descriptions within challenging and poorly differentiated structures, such as the lateral hypothalamic zone. Each identified gray matter connection was then compared with peer-reviewed datasets to establish a degree of coherence within the CNG pathway tracing literature. Our maps, in addition to being largely harmonious with the existing literature, emphasize the relevance of connectional topography and reciprocity with respect to the study of CNG connections and motivated behaviors.

**Figure 4.**
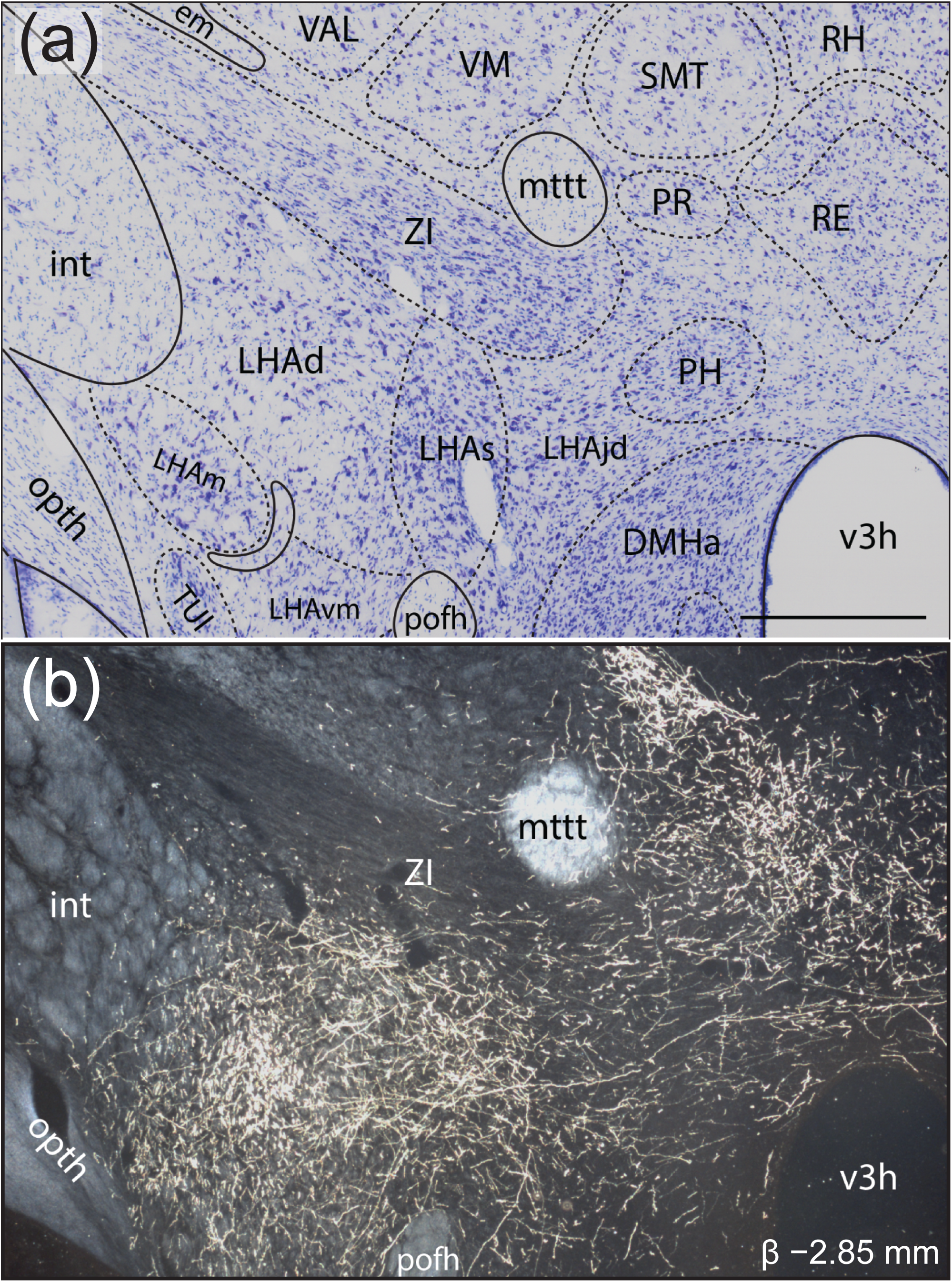
Photomicrographs of PHAL-immunoreactive (-ir) axons from the ILA (experiment # 15-113) in the *dorsal (Swanson, 2004)* (LHAd) and *suprafornical (Swanson, 2004)* (LHAs) *regions* of the *lateral hypothalamic area (Nissl, 1913)*. **(a)** Adjacent Nissl-stained section with boundaries and terminology based on Swanson (2018). **(b)** Darkfield photomicrograph showing PHAL-ir axons. Spatial alignment, labels, and scale bar were derived from the reference Nissl-stained section in **(a)**. Scale bar: 500 µm. See the list of abbreviations on p. 35 for an explanation of the abbreviations shown in this figure.

**Figure 5.**
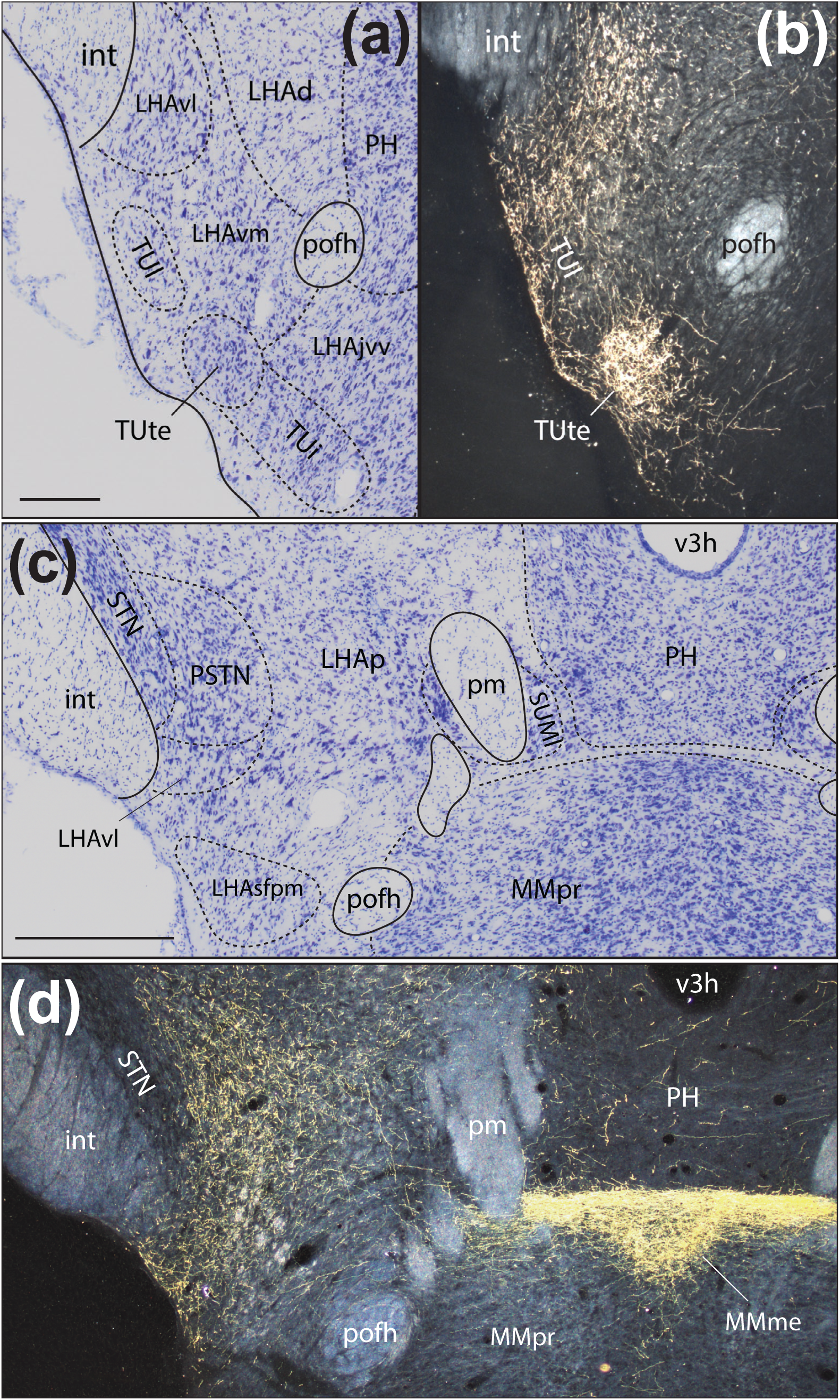
Photomicrographs of PHAL-immunoreactive (-ir) axons from the ILA (experiment # 15-113) in the *terete part (Petrovich et al., 2001)* (TUte) of the *tuberal nucleus (>1840)* **(a, b)**, the *parasubthalamic nucleus (Wang & Zhang, 1995)* (PSTN) and *median part (>1840)* (MMme) of the *medial mammillary nucleus (Gudden, 1881)* (c, d). Adjacent Nissl-stained sections showing the TUte **(a)** and PSTN **(c)**. Boundaries and terminology were based on Swanson (2018) and superimposed on darkfield photomicrographs showing PHAL-ir axons in the TUte **(b)**, PSTN and MMme **(d)**. Scale bars: 200 µm in **(a)**; 500 µm in **(c)**. See the list of abbreviations on p. 35 for an explanation of the abbreviations shown in this figure.

**Figure 6.**
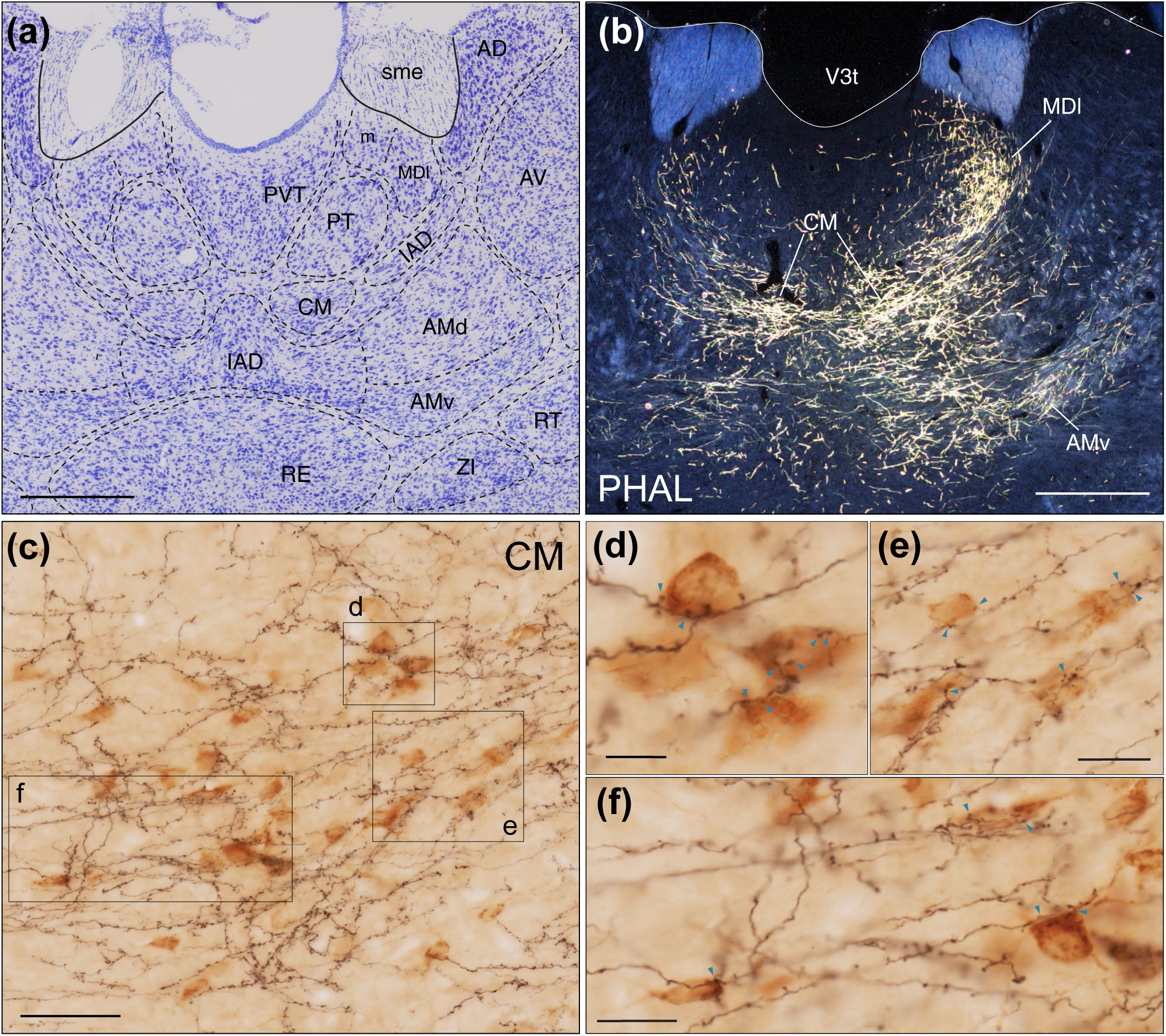
Photomicrographs showing ACAd axons (experiment #15-131) in rostral *thalamus (His, 1893)* and the formation of putative monosynaptic reciprocal connectivity in the c*entral medial thalamic nucleus (Rioch, 1929)* (CM). **(a)** Adjacent Nissl-stained photomicrograph showing superimposed boundaries and regional terms based on Swanson (2018). **(b)** Darkfield photomicrographs showing immunoreactive (-ir) labeling in the CM, *mediodorsal (>1840)* and *anteromedial (>1840) thalamic nuclei* based on the Nissl stain in **(a). (c)** Extended-focus image showing PHAL-ir axons (*black*) and CTB-ir neurons *(brown*) in the CM captured with ×100 objective lenses. **(d, e, f)** Mosaics made with single *z*-plane images to show putative appositions (*blue arrowheads*) from regions indicated in **(c).** Scale bars: 500 µm in **(a)** and **(b)**; 50 µm in **(c);** 10 µm in **(d)**; 20 µm in **(e)** and **(f).** See the list of abbreviations on p. 35 for an explanation of the abbreviations shown in this figure.

### 4.1: Methodological considerations

Pathway tracing can be performed by using a wide variety of tracers, each with different sensitivities, effective time windows, and uptake mechanisms. There is some discussion about the variable efficacies of tracers (Bota et al., 2003; Lanciego and Wouterlood, 2006), but little work has been done to systematically compare and validate them (Ter Horst et al., 1984; Wang et al., 2014; Calabrese et al., 2015). The tracer combination of PHAL and CTB used here was based on their resistance to uptake by fibers-of-passage. PHAL uptake preferentially occurs through dendrites and possibly cell bodies; with rare exceptions, PHAL is only transported in the anterograde direction (Gerfen and Sawchenko, 1984). From all of our experiments, only a single retrogradely labeled PHAL cell body was observed in the field CA1 of the hippocampus (*not shown*). We operated, as customarily done, with the assumption that PHAL-labeled axons detected in the diencephalon originated from CNG cell bodies that were clearly filled with PHAL. CTB transport, by contrast, is known to have both anterograde and retrograde properties (Luppi et al., 1990). Uptake by damaged fibers-of-passage has not been shown to occur for CTB and the necrosis dealt through the injection process is largely avoided with the iontophoretic approach. Nonetheless, the CNG and cortex in general is not recognized as a route for passing fibers. We can therefore conclude that the caveats associated with our anterograde tracers are of less concern in the present study.

**Figure 7.**
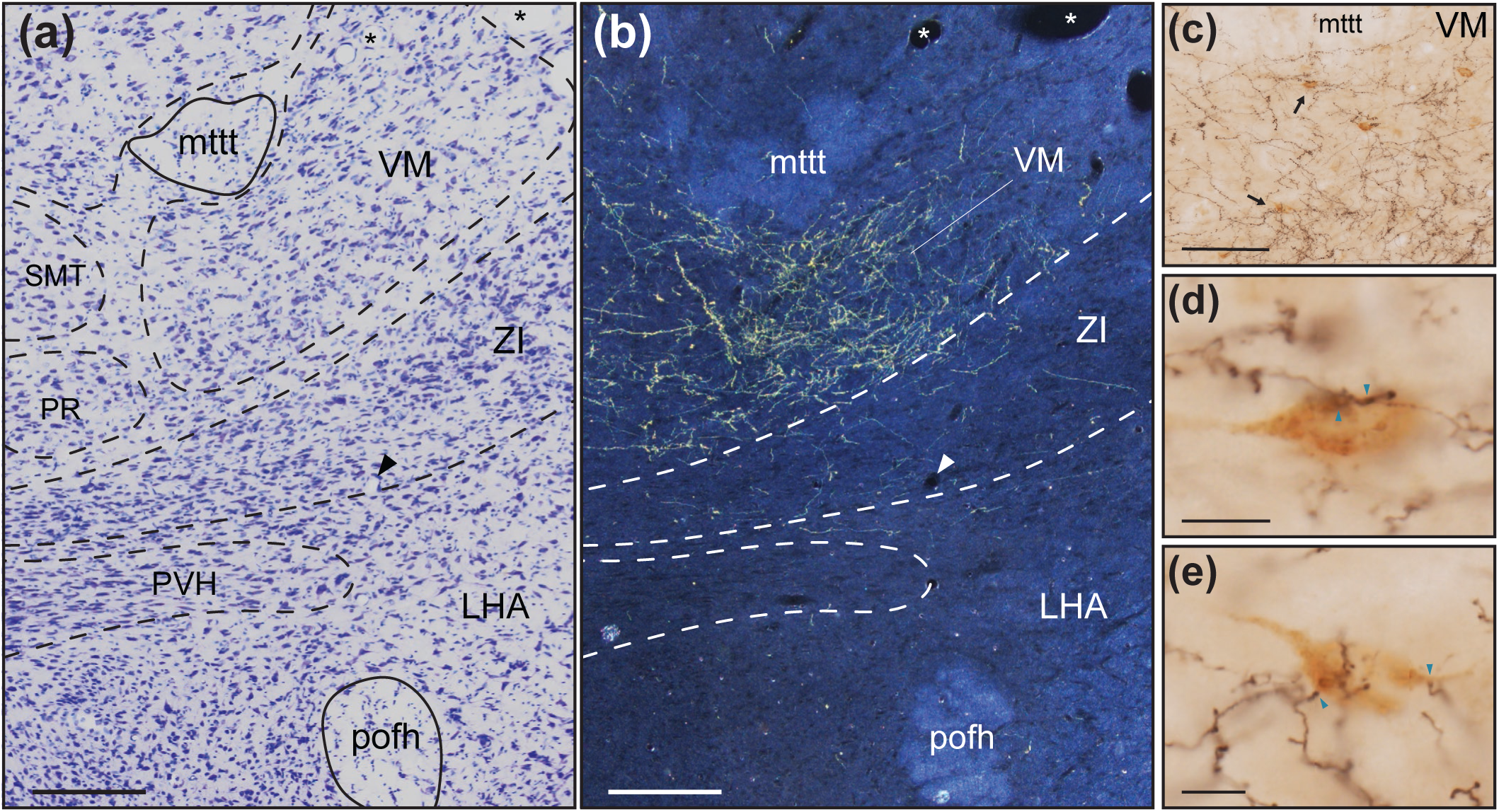
Photomicrographs showing ACAd axon terminals (experiment #15-131) in a rostromedial part of the *ventral medial thalamic nucleus (>1840)* (VM), and not the *zona incerta (>1840)* (ZI). **(a)** Adjacent Nissl-stained photomicrograph showing superimposed boundaries and regional terms based on Swanson (2018). **(b)** Darkfield photomicrographs showing immunoreactive (-ir) labeling in the VM immediately ventral to the *thalamic mammillothalamic tract (Swanson, 2015)* (mttt). *Asterisks* and *arrowheads* in **(a)** and **(b)** show common vasculature between both sections. **(c)** Extended-focus image showing PHAL-ir axons (*black*) and CTB-ir neurons (*brown*) in the VM captured with ×100 objective lenses. **(d, e)** Mosaics made with single *z*-plane images to show putative appositions (*blue arrowheads*) on neurons indicated with *arrows* in **(c).** Scale bars: 200 µm in **(a)** and **(b)**; 100 µm in **(c)**; 10 µm in **(d)** and **(e).** See the list of abbreviations on p. 35 for an explanation of the abbreviations shown in this figure.

Atlas-based mapping was used here to localize injection sites and tracer transport, and we have used this approach to document our anatomical datasets (*e.g.*, Zséli et al., 2016; Santarelli et al., 2018; Martínez et al., 2023; Tapia/Agostinelli et al., 2023) and have also encouraged the use of standardized, atlas-based approaches more generally (Khan, 2013; Khan et al., 2018a,b; Khan et al., 2021). Standardized mapping was achieved using the cytoarchitectonic definitions of gray matter regions described in *Brain Maps 4.0* (Swanson, 2018). Nissl-defined boundaries are advantageous because they facilitate stable and unambiguous representations of histological data, allowing us to locate and precisely relate the same anatomical space from several brains. We leveraged this to represent all of our findings with atlas-level precision. The approach itself has been widely used to represent anatomical information and some recent advances in the use and development of metadata analysis tools should be highlighted. Cytoarchitectonic regularity is such that stereotaxic coordinates can be inferred from them (Khan et al., 2018b) and datasets derived from different brains can be combined (Khan, 2013). The latter feature was well-represented in a study that combined gene expression and connectional data to describe a subregional architecture of the hippocampus (Bienkowski et al., 2018). A combination of anatomical datasets was also instrumental for a recent series of connectomic studies (Bota et al., 2015; Swanson et al., 2016; 2017; 2018; Hahn et al., 2019; Swanson et al., 2019a,b; 2020; 2021; 2022; 2023; 2024a,b). Such metadata analyses necessitate the use of a standardized and hierarchically organized nomenclature system (Swanson, 2015; 2018). The maps shown here satisfy these criteria and they show connectional information at atlas-level resolution.

### 4.2: Comparison of findings with other tract-tracing studies

The input and output connections of the rat CNG are among the most intensely studied. Decades of research involving several successions of pathway-tracing strategies have described the general features of CNG connections (Nauta, 1964; Domesick, 1969; Krettek and Price, 1977; Shiosaka et al., 1980; Brittain, 1988; Sesack et al., 1989; Hoover and Vertes, 2007). However, previous studies did not represent CNG connectivity bidirectionally or achieve a similar level of spatial resolution. Doing so allowed us to observe CNG connections in the diencephalon with high subregional precision.

In the following sections, we describe how our work aligns with other pathway tracing studies that involved injections into the CNG or *interbrain (Baer, 1837)* (IB) (diencephalon). Several reports have examined ILA efferents throughout the diencephalon using the PHAL method (Brittain, 1988; Sesack et al., 1989; Hurley et al., 1991; Vertes, 2004) but fewer PHAL studies have focused on ACAd efferents (Sesack et al., 1989), though there is some coverage achieved with the biotinylated dextran amine method (Balfour et al., 2006). CNG afferents from the diencephalon were best captured by a retrograde study using Fluoro-Gold (Hoover and Vertes, 2007) and an earlier report involving Diamidino Yellow and True Blue (Condé et al., 1995). Our results, despite differing in methodologies from the above reports, are remarkably well-aligned with their findings.

#### 4.2.1: Hypothalamus (Kuhlenbeck, 1927) (HY)

CNG connections with the HY were predominantly observed in our ILA injections and they favored a top-down direction (**Table 2**). As in *thalamus (His, 1893a)* (TH), CNG axons were commonly observed bilaterally but were always less dense on the contralateral side. CNG inputs from HY did show contralateral innervation, a feature not applicable to TH (**Figure 3**). For simplicity, we will discuss our findings for HY across three major divisions: the *periventricular* (HYp), *medial* (HYm), and *lateral* (HYl) *hypothalamic zones (Nauta & Haymaker, 1969)*.

##### 4.2.1 a: Periventricular hypothalamic zone (Nauta & Haymaker, 1969) (HYp)

CNG connections with the HYp tended to be weak in both directions. In general, connectivity in the HYp took the form of diffuse, low density ILA axons with only a few retrogradely labeled cells. Retrograde studies support our observed ILA and ACAd projections for the MEPO (Saper and Levisohn, 1983), PV anterior zone (PVaa) (De Haro, 2015), MPO (Kita and Oomura, 1982; Chiba and Murata, 1985), ARH (Magoul et al., 1993a, 1993b; Martinez, 2017), DMH (Thompson and Swanson, 1998), and PH (Abrahamson and Moore, 2001). Retrograde labeling from the CNG was sparse in the HYp. The few ILA afferents found in a caudal and ventral part of the DMH were not supported by other retrograde studies in the ILA (Condé et al., 1995; Hoover and Vertes, 2007); likewise, PHAL injections into the DMH did not produce labeling of the CNG (ter Horst and Luiten, 1986; Thompson et al., 1996). CNG inputs from the PH are, in part, supported by one PHAL study (Vertes et al., 1995).

The PVH is an especially confounding structure. It is a small, neurochemically diverse region that contains a plurality of subdivisions (Simmons and Swanson, 2009b; Biag et al., 2011). PVH connectivity is likewise difficult to resolve. Our preparations showed ILA axons throughout the PVH (**Figure 3**), a finding that is in agreement with other PHAL studies (Brittain, 1988; Sesack et al., 1989; Vertes, 2004) but others, including several retrograde studies of the PVH, have not reported labeling in the CNG (Silverman et al., 1981; Sawchenko and Swanson, 1983; Hurley et al., 1991).

##### 4.2.1 b: Medial hypothalamic zone (Nauta & Haymaker, 1969) (HYm)

CNG connections in the HYm tended to follow the same diffuse, low densities as in the HYp (**Table 2**). Light ILA innervation of the MPN was consistent with one retrograde tracing study (Simerly and Swanson, 1986). We did not observe MPN subdivision specificity but another investigation showed dense ILA axons in the lateral part of the MPN (Brittain, 1988). Retrograde tracing, in the present and other reports, have not shown transport to the MPN (Condé et al., 1995; Hoover and Vertes, 2007). However, PHAL injections into the MPN showed terminals in layer 6 of the ILA (Simerly and Swanson, 1988). Most anterograde studies showed ILA projections to the AHN (Brittain, 1988; Hurley et al., 1991; Vertes, 2004); there were no retrograde tracing studies in the AHN, to our knowledge, that corroborate these observations. CTB labeling of AHN was not observed in our experiments, the same lack of labeling was observed in one other retrograde tracer study (Condé et al., 1995). There are nonetheless at least two reports that show AHN projections to the ILA (Risold et al., 1994; Hoover and Vertes, 2007). Our work and others have not identified CNG projections to the VMH (Brittain, 1988; Hurley et al., 1991; Vertes, 2004; Toth et al., 2010; Shimogawa et al., 2015). We did, however, observe retrograde labeling in the VMH from CTB injections in ILA and ACAd. This finding has some support (Condé et al., 1995; Canteras et al., 1994) but VMH projections to the CNG were largely not observed in other studies (Saper et al., 1976; Krieger et al., 1979; Hoover and Vertes, 2007; Shimogawa et al., 2015). Retrograde labeling was not detected in the ventral mammillary nucleus (PMv) although its projections to the ILA have been described (Canteras et al., 1992b), this may be due to a slight spillover of injected PHAL into the TUte laterally.

The medial mammillary nucleus (MM) contained the densest ILA terminals observed. We have localized this to a dorsal part of the MM, specifically the MMme, and not the more caudal SUM (**Fig. 5c, d**). Our observations, of both the localization and density of this terminal, were also demonstrated with WGA-HRP injections into the CNG by Allen and Hopkins (1989). Subsequent retrograde tracers deposited into the MM showed that these projections almost exclusively originated from a part of the ILA that corresponded to the “dorsal peduncular cortex” (Paxinos and Watson, 2014). This projection is not described in the majority of ILA pathway tracing studies (Sesack et al., 1989; Vertes, 2004) but the same conclusion was reached by one other group (Hurley et al., 1991).

##### 4.2.1 c: Lateral hypothalamic zone (Nauta & Haymaker, 1969) (HYl)

CNG projections to the LPO have been described in some detail in anterograde tracing studies (Brittain, 1988; Hurley et al., 1991; Vertes, 2004), and at least one retrograde study in the rat (Shiosaka et al., 1980). LPO outputs to the CNG shown here are supported by at least two other reports (Swanson, 1976; Hoover and Vertes, 2007). The LHAa contained the most retrograde labeling in the hypothalamus. This can also be observed in maps produced by Vertes (2004) but here we localized it to the LHAa and, more precisely, to its ventral region (LHAav). Unfortunately, the input and output connections of the LHAa have not been studied in much detail. We were only able to locate one study that reported ILA projections to the LHAav (Thompson and Swanson, 2010) and a PHAL study showed that a medial part of LHAav did not project to CNG (Canteras et al., 2011). One study in mice recently showed bidirectional connectivity between CNG and both LPO and LHAa (Hahn et al., 2022). Their experiments showed preferential connectivity with ventral CNG predominantly with the LHAa.

Connections found in the LHAjv were well-supported by other reports (Gabbot et al., 2005; Hoover and Vertes, 2007; Hahn and Swanson, 2015; Reppucci and Petrovich, 2016). CNG connections with the LHAjp and LHAjd were also in line with other tract tracing studies (Condé et al., 1995; Gabbot et al., 2005; Yoshida et al., 2006; Hahn and Swanson, 2010, 2012). ILA and ACAd projections to the subfornical region were supported in studies that covered a wide portion of the LHA with retrograde injections (Gabbott et al., 2005; Repucci and Petrovich, 2016). Projections to the LHAs were confirmed with small injections of retrograde tracers (Yoshida et al., 2006; Hahn and Swanson, 2010). CNG afferents from the subfornical region were shown in the LHAsfa (Goto et al., 2005) but a small injection to the LHAs failed to produce any anterograde labeling in the CNG (Hahn and Swanson, 2010).

We have demonstrated a dense and restricted ILA projection to the TUte (**Fig. 5a, b**). This projection was noted in an ILA study by Hurley et al. (1991), but it was not ascribed to the TUte (*see their* Figure 5D). CNG projections to the tuberal nucleus have received little attention compared to the rest of the hypothalamus. Only one report incidentally labeled the TUl as part of an injection covering the ventral half of the LHA (Repucci and Petrovich, 2016). CNG afferents from TUi and TUte were demonstrated in one PHAL study (Canteras et al., 1994).

The ventral region of the LHA has received little attention and, consequently, none of the connections described here have LHAv injections to compare with. There is strong agreement with other CNG tracing studies (Brittain, 1988; Hurley et al., 1991; Vertes, 2004) but LHA projections in the CNG have not been described with clear attention to this structure (Condé et al., 1995; Hoover and Vertes, 2007).

We described the morphology of ILA axons in the LHAd, where numerous mfb collaterals were formed and bore the characteristics of axon terminals (**Fig. 3l–n**). Several retrograde tracing studies support this with large injections into LHA (Shiosaka et al., 1980; Hurley et al., 1991; Gabbott et al., 2005; Yoshida et al., 2006) and small PHAL and CTB co-injections restricted to the LHAd revealed retrograde but not anterograde labeling in the CNG (*J. D. Hahn and V. I. Navarro, personal communications*).

Another set of dense CNG collaterals was observed in the PSTN and LHAp (**Fig. 3q, r**). The existence of ILA terminals in the PSTN has been mentioned but not shown in a retrograde study of the PSTN (Chometton et al., 2015). We additionally noted a few retrogradely labeled PSTN neurons from CTB injections to the ILA. These were not detected in other retrograde studies of the CNG (Condé et al., 1995; Hoover and Vertes, 2007) but Goto and Swanson (2004) reported, with anterograde injections into the PSTN, that its projections were remarkably specific to the ILA. We showed ACAd axons in the STN. Canteras et al., (1990) likewise showed ACAd projections to a caudal part of the STN.

Although we demonstrated remarkable specificity in CNG projections to the hypothalamus, it is important to note that a failure to discover CNG axons within a given nucleus should not constitute an absence of connectivity. For example, it is understood that certain hypothalamic neurons can form long dendrites that extend well beyond their nuclear boundaries (Millhouse, 1969).

#### 4.2.2: Thalamus

Our discussion of CNG connections with thalamus is divided according to its two major divisions: the *ventral* and *dorsal parts (Herrick, 1910)*.

##### 4.2.2 a: Ventral part of thalamus (Herrick, 1910) (THv)

ACAd and ILA projections to the ZI were observed in several retrograde studies (Mitrofanis and Mikuletic, 1999; Chometton et al., 2017; Chou et al., 2018). However, our demonstration of dense ACAd and ILA terminals in dorsally and ventrally adjacent areas highlights an important interpretive caveat; specifically, that slight spillover of tracers or infused compounds could result in effects that are not specific to ZI. ACAd terminals in ZI are nevertheless supported by at least two reports that used anterograde tracers (Mitrofanis and Mikuletic, 1999; Balfour et al., 2006). Retrograde labeling from the CNG to the ZI reported here were previously shown (Condé et al., 1995; Hoover and Vertes, 2007); but anterograde studies of the ZI failed to produce CNG labeling (Wagner et al., 1995; Sita et al., 2007), possibly due to the locations of deposited tracers. CNG projections to the RT were demonstrated here with ACAd injections. This connection is corroborated by one retrograde tracer study of the RT (Cornwall et al., 1990).

##### 4.2.2 b: Dorsal part of thalamus (Herrick, 1910) (THd)

The *midline thalamic nuclei (>1840)* are among the most interconnected with the CNG. Every subdivision of the RE was bidirectionally connected with the ILA in moderate or high densities, as shown in other reports (Van der Werf et al., 2002; Vertes, 2002; McKenna and Vertes, 2004; Vertes et al., 2006; Varela et al., 2014). ILA tracing labeled the PVT bidirectionally, consistent with a number of studies (Groenewegen, 1988; Chen and Su, 1990; Berendse and Groenewegen, 1991; Hurley et al., 1991; Moga et al., 1995; Van der Werf et al., 2002; Vertes, 2002; Gabbott et al., 2005; Vertes and Hoover, 2008). However, the same caveat offered for ZI is applicable here as PVT is surrounded by regions that are strongly connected with ILA. Bidirectional connectivity between the PT and both ILA and ACAd was evident from previous work as well (Berendse and Groenewegen, 1991; Chen and Su, 1990; Van der Werf et al., 2002; Vertes, 2002; McDonald et al., 1999; Vertes and Hoover, 2008; Varela et al., 2014).

*Anterior thalamic nuclei (>1840)* were mainly connected with the ACAd. Bidirectional connectivity between the AM and ACAd has been shown previously (Shibata, 1993; van Groen et al., 1999; Shibata and Naito, 2005) with its ventral part being the most strongly connected (Vertes, 2002; Hoover and Vertes, 2007; de Lima et al., 2017). ACAd injections from other reports also revealed slightly less dense connections with the IAM (Vertes, 2002; Hoover and Vertes, 2007) and the IAD (Vertes, 2002).

The *medial thalamic nuclei (>1840)* are also well-connected with the CNG. Bidirectional connectivity with the PR, with a pronounced topographic organization, is in line with other studies (Van Der Werf, 2002; Vertes, 2002; Hoover and Vertes, 2007). Our finding of dense ILA terminals in a ventrocaudal half of the SMT was observed in one other report (Hurley et al., 1991) but was absent in most studies that described ILA outputs (Brittain, 1988; Sesack et al., 1989; Vertes, 2002; 2004). This difference is likely due to our inclusion of a more caudal and ventral portion of the ILA than typically observed in studies of this region. This conclusion is supported by an investigation of SMT that reported retrograde labeling in the ventral half of the caudal ILA but not in the TTd (Yoshida et al., 1992). CNG connections with the MD and IMD have been examined in several earlier reports (Leonard, 1969; Krettek and Price, 1977; Groenewegen, 1988) but none have highlighted the extent of spatial overlap between input and output connections (**Fig. 3j–r**); moreover, the precise topographic organization of MD connections has yet to be characterized.

*Intralaminar thalamic nuclei (>1840)* were variably connected with the CNG and exhibited pronounced spatial topography (**Fig. 3k–r**). In agreement with other studies, the rostral CM showed strong, bidirectional connectivity with the ACAd (**Figure 6**) (Berendse and Groenewegen, 1991; Vertes, 2002; Van der Werf, 2002; Vertes et al., 2012). CM projections to the ILA arose from slightly more caudal parts (Van der Werf, 2002; Hoover and Vertes, 2007; Vertes et al., 2012). ILA retrograde labeling identified RH projections; these were previously shown to only target the ventral ILA (Van der Werf, 2002). A few anterograde studies focused on the CL did not identify projections to the ACAd (Van der Werf, 2002; Wang and Shyu, 2004). This can be explained by the locations of CL injections, which were ventral and rostral to the parts where we observed retrograde labeling (**Fig. 3m–r**; Vertes, 2002; Hoover and Vertes, 2007). Anterograde and retrograde tracing of the PF did not show connectivity with the CNG (Cornwall and Phillipson, 1988; Berendse and Groenewegen, 1991; Van der Werf, 2002). We, and others, have shown low density ILA and ACAd connections with the PF in its more dorsal and medial parts (Condé et al., 1995; Vertes, 2002; Hoover and Vertes, 2007).

A few *ventral thalamic nuclei (>1840)* were shown to be connected with the ACAd. ACAd inputs and outputs to the VAL have been shown by others (Vertes, 2002; Hoover and Vertes, 2007), but our preparations only showed bidirectional ACAd connectivity restricted to a rostral aspect of the VAL (**Fig. 3k, l**). Previously described ACAd inputs from the VM were identified here (Hoover and Vertes, 2007), but were present well within the VM (Vertes, 2002; Balfour et al., 2006). We observed dense ACAd terminals immediately ventral to the mammillothalamic tract (**Fig. 3l, m** and **Figure 7**), this connection was noted in one other study and was interpreted as part of the RE (Vertes, 2002).

Finally, CNG connections with the *epithalamus (His 1893b)* were previously described (Vertes, 2002; Kim and Lee, 2012). We found ILA and mainly ACAd projections terminating in similar parts of the LH.

### 4.3: Networks, second-order connections

Recent work has focused on the assembly of connectomes based on data curated from the tract tracing literature. This project has examined the cerebral cortex (Bota et al., 2015; Swanson et al., 2017, 2018), cerebral nuclei (Swanson et al., 2016, 2018), diencephalon (Hahn et al., 2019; Swanson et al., 2019a, 2019b), midbrain (Swanson et al., 2021), rhombicbrain (Swanson et al., 2022) and spinal cord (Swanson et al., 2024a). Modularity analysis performed on connectomes for cerebral cortex and cerebral nuclei (“basal ganglia”) structures consistently placed the ILA and ACAd in separate modules (Swanson et al., 2018). The ILA was grouped in a module that was associated with somatic and visceral information whereas the ACAd was associated with the default mode network (DMN) (Swanson et al., 2018). In agreement with this, recent work has shown that chemogenetic silencing of the ACAd altered DMN activity, resulting in less time spent sitting still and more time engaging in rearing (Tu et al., 2020). A more recent analysis of the forebrain connectome placed the ILA in a subsystem that supports pheromonal sensorimotor functions while the ACAd contributes to a subsystem that supports voluntary eye and nose movements (Swanson et al., 2020). The forebrain connectome was arranged into two subsystems at the highest level. One subsystem was associated with voluntary control of behaviors and the second with innate survival behaviors and physiology (Swanson et al., 2020). Intriguingly, these subsystems were respectively associated with the lateral forebrain bundle (internal capsule) and mfb. This finding aligns remarkably well with our description of fiber systems used by ACAd and ILA projections.

Connectomes do well to illustrate that regions can leverage second-order connections when direct projections are not available. Several instances of this have been reported for CNG. For example, we and others have shown weak ILA projections to the PVH (Vertes, 2004), a region directly involved in the release of signals regulating hormone secretion from the pituitary gland (Swanson and Sawchenko, 1983). Unexpectedly, lesions delivered to the CNG increased PVH activation following restraint stress, likely due to a reduction in excitatory tone from CNG inputs to PVH-projecting GABA neurons in the BST (Radley et al., 2009). Similarly, CNG connections with the *suprachiasmatic nucleus (Spiegel & Zwieg, 1919)* (SCH) are virtually absent (**Figure 3**). A transneuronal tracing study demonstrated a disynaptic input from the SCH to the ILA that involved a relay through the PVT (Sylvester et al., 2002).

ACAd outputs to the hypothalamus are sparse. They instead target the thalamus in places distinguishable from ILA terminals. The AM is reciprocally connected with the ACAd (**Figs. 3j, k**). This region, in turn, projects to the CNG, *anteromedial* (VISam) and *posteromedial* (VISpm) *visual areas (Espinoza & Thomas, 1983)*, *perirhinal area (Brodmann, 1909)*, *retrosplenial area (Vogt & Peters, 1981)*, *presubiculum (Swanson & Cowan, 1977)*, lateral part of the *entorhinal area (Brodmann, 1909)*, and *temporal association areas (Swanson, 1992)*, all members of the so-called default mode network in rats (Lu et al., 2012; de Lima et al., 2017). Interestingly, each of these cortical areas also receive direct projections from the ACAd (Vogt and Miller, 1983; Jones et al., 2005; Jones and Witter, 2007; Shibata and Naito, 2008). These connectional features suggest that the ACAd is strongly interconnected with members of the default mode network through both direct (cortico-cortical) and indirect (corticothalamic) pathways.

In addition to polysynaptic connections, CNG and other cortical outputs can participate in parallel and convergent pathways (Swanson, 2000). ILA projections directly target the LHAa (**Figure 3**) and a dorsomedial part of the *accumbens nucleus (Ziehen, 1897–1901)* (ACB), which also projects to the LHAa (Thompson and Swanson, 2010). Similar observations were made in the *bed nuclei of terminal stria (Gurdjian, 1925)* (BST; Dong and Swanson, 2003, 2004, 2006a), the *lateral part (Swanson, 1992)* of the *central amygdalar nucleus (Johnston, 1923)* (CEAl; Barbier et al., 2018a, 2018b), the *medial amygdalar nucleus (Johnston, 1923)* (Canteras et al., 1995), and *basomedial amygdalar nucleus (>1840)* (Reppucci and Petrovich, 2016) with respect to axon terminals converging on the LHAs and LHAd. This pattern of descending cortical inputs to striatal and pallidal structures, and the convergence of their projections in the hypothalamus has been hypothesized for the control of motivated behaviors (Swanson, 2000). Upon closer examination, there is a remarkable degree of spatial overlap between projections from the ILA (**Figure 3**), CEA (Barbier et al., 2018a), and BST (Dong and Swanson, 2003, 2004, 2006a). Specifically, they all converged in a small part of the LHA that was consistent with the LHAs and LHAd (approx. −2.45 mm from bregma). It is important to point out that the LHAd is cytoarchitectonically characterized as a cell-sparse region relative to neighboring structures (Swanson, 2018). Given that ILA, CEA, BMA, and BST all formed similarly dense collaterals in the LHAd, it is plausible that cell-sparsity in this region reflects sheer space needed to form collaterals from many regions. Our description matches that of a classic Golgi study of the mfb which showed a massive group of axon collaterals exiting the mfb at roughly the same location, anteroposterior span, and orientation as those in the LHAd (Millhouse, 1969). Previous work has identified the LHAd as the region with the most hypocretin/orexin and melanin-concentrating hormone neurons (Hahn, 2010). Finally, recent functional mapping of LHA found that optic stimulation, specifically in a stereotaxic space corresponding to LHAd, was sufficient to induce feeding and self-stimulation (Urstadt and Berridge, 2020). Therefore, the provisional description of a Nissl-defined boundary between the LHAa and LHAd indicates a transition that is meaningful on connectional, neurochemical, and functional grounds.

### 4.4: Topography of thalamocortical connections

Models for cerebral hemisphere organization often include descriptions of parallel and segregated circuits that include components of cortex, striatum, and pallidum (Alexander et al., 1990; Swanson, 2000). At the level of cortico-striatal pathways, recent work combining tract-tracing and atlas-based mapping in the mouse delineated the dorsal striatum into 29 distinct domains on the basis of cortico-striatal projections (Hinitiryan et al., 2016). Similarly, our spatial framework allows detection of distinct topographic organization within the corticothalamic projectome (**Figure 3**). This is particularly applicable to the MD, where CNG connections occupied minimally overlapping domains that varied across three dimensions. The topography of MD neurons projecting to the PFC has been explored (Alcaraz et al., 2016) but has yet to be carried out with high-spatial-resolution analysis or with an atlas-based framework that would allow comparisons with other neuroanatomical datasets.

Our observed ACAd connections with MDl were strikingly similar to a description of projections from the *nucleus incertus diffuse part (Goto et al., 2001)* (NId) (*see* Fig. 7D, E from Goto et al., 2001). NId terminals, like the ACAd axons (**Figure 7**), were clustered in a space between the mammillothalamic tract and the ZI which they referred to as the rostromedial tip of the VM (Goto et al., 2001). Both the MDl and VM project to a part of the secondary somatomotor area that is considered the “frontal eye field” in the rat (Reep et al., 1984). ACAd outputs can therefore leverage multiple thalamic regions that control visual attention.

Our co-injection studies demonstrated a remarkable degree of spatial overlap between CNG connections in the thalamus; a similar observation was made with HRP implants in the macaque prefrontal cortex (Preuss and Goldman-Rakic, 1987). Analysis of reciprocal connections at high-magnification frequently revealed close appositions of anterogradely labeled axons on retrogradely labeled somata in the thalamus. Although this should not be taken for evidence of monosynaptic loops, it is nonetheless a structural requirement for their existence. Moreover, our observation that CNG inputs and outputs are tightly coupled in space suggests that reciprocal loops may be a general feature of corticothalamic connections. It is still unclear how reciprocal connectivity might contribute to CNG and thalamic functions. There is evidence that the MD contributes to attentional control by amplifying cortical activity during task engagement (Schmitt et al., 2017). Parallel and segregated loops, in this manner, may allow CNG ensembles to self-maintain local activity during tasks. At the level of single-cell morphology, individual MD neurons are known to target multiple and widespread cortical areas (Kuramoto et al., 2016). It was also shown that individual thalamic neurons, at least those in the RH and parafascicular nucleus, formed dense terminals in the striatum in addition to the cortex (Fujiyama et al., 2019). This suggests that reciprocal connectivity between CNG and thalamus is likely coupled with thalamostriatal and corticostriatal pathways.

### 4.5: Functional significance of ILA connections in the context of ingestive behaviors

ILA projections in the *lateral hypothalamic zone (Nauta & Haymaker, 1969)* primarily branched from the *medial forebrain bundle (Edinger, 1893)*. ILA collaterals mainly targeted dorsal parts of the LHAd and LHAs (**Figs. 3** and **4**), both parts of LHA that are connectionally positioned to modulate ingestive behavior (Hahn and Swanson, 2010). In line with this, chemogenetic activation of the ILA was shown to induce feeding in the absence of hunger (Mena et al., 2011), ostensibly by increasing neural activation in a part of the LHA dorsal to the fornix (Mena et al., 2013). The space described roughly corresponds to the LHAs and LHAd, and may recruit H/O neurons among other populations (Mena et al., 2013). This area also corresponds to where injection sites for glutamate-stimulated feeding were migrated from a previous study (Khan et al., 2004), and placed within *Brain Maps 4.0* reference space (Khan et al., 2018b). ILA stimulation is also known to increase arterial blood pressure, an effect that can be reversed by silencing the LHAd/s (Fisk and Wyss, 2000). ILA projections to this area are therefore capable of affecting motivational as well as autonomic outcomes.

More caudally, ILA axons also densely innervated the PSTN (**Figure 3**). Input and output connections of the PSTN are well-described (Goto and Swanson, 2004; Chometton et al., 2015). PSTN projections heavily target hindbrain regions involved in processing gustatory and viscerosensory information as well as preganglionic parasympathetic cell groups that control autonomic responses (Goto and Swanson, 2004). The PSTN stands out as a part of the lateral zone that virtually lacks GABAergic neurons, a marked contrast with the nearby GABA-rich LHAs and LHAd (Chometton et al., 2015). It is still unclear how ILA contributes to PSTN functions. Recent work described a basal ganglia-like circuit motif which included the insular cortex, central nucleus of the amygdala, and substantia innominata (alternatively, the ‘innominate substance’ [Swanson, 2015]) (Barbier et al., 2020). Subsequent chemogenetic inhibition was used to demonstrate that PSTN activity was necessary to suppress feeding during illness and, to some extent, neophobia (Barbier et al., 2020). Here, we showed that ILA contributes substantial innervation to the PSTN. Moreover, our injection experiments and work done by Vertes (2004) show strong ILA projections to the feeding “no-go” circuit. It is therefore worth examining how ILA contributes to PSTN-mediated control of behaviors.

CNG activity is thought to support the associative learning of—and the ability to act on—food-related cues (Petrovich, 2011). Indeed, Pavlovian conditioning depends on an intact CNG to trigger feeding in sated rats (Petrovich et al., 2007). Recent work has also shown that sated rats could be induced to overeat by infusing µ-opioid receptor agonists into the ventral CNG (Mena et al., 2011). Overeating driven by associative learning and the endocannabinoid system clearly engage ventral CNG structures, but the degree to which they share neural substrates is not well understood. Both approaches have demonstrated selective Fos induction in H/O-ir neurons dorsal to the fornix at the level of the DMH (Petrovich et al., 2012; Mena et al., 2013). This region likely corresponds to the LHAs and LHAd, which contain a large proportion of hypothalamic H/O neurons (Swanson et al., 2005; Hahn, 2010). Here we showed that ILA projections, mostly from the caudal part, gave rise to terminal fields in the LHAs (**Fig. 3h**). The possibility of ILA projections forming monosynaptic contacts on H/O neurons has been explored in one study which found that ILA axons formed putative appositions with at least 22% of H/O cells (Yoshida et al., 2006).

### 4.6: Concluding remarks

This work was inspired, in part, by the findings that CNG could potently control feeding behaviors and hedonic processes (Mena et al., 2011; Castro and Berridge, 2017). As part of a structural complement to these findings, we provided detailed maps of CNG connections in the diencephalon that highlight potential pathways that mediate these effects. Our maps provide sub-millimeter precision for targeting CNG terminals. Given that optogenetic and chemogenetic manipulation of axon terminals actually affect fiber systems rather than individual terminals, CNG “tractography” allows scrutiny of pathways that are inadvertently affected during such manipulations. A major problem in the study of prefrontal cortex is confusion in the nomenclature and boundary definitions across and within models (Laubach et al., 2018). We demonstrated that connectivity between CNG and thalamus was predominantly reciprocal and non-overlapping in the case of ILA and ACAd. One can imagine a redefinition of CNG boundaries based on degree of overlap or using connectional “fingerprints.” In this sense, brain atlases would function more as stable coordinate systems in which boundaries are iteratively redrawn as we continue to understand the brain in its own language (Buzsáki, 2019).

## Author Contributions

Experimental conception and design: AMK, KN. Project management: KN, AMK. Surgeries and *in vivo* experiments: KN. Perfusions and histology: KN, LPM, VN. Nissl staining and immunohistochemistry: KN, LPM, CO. Microscopy and imaging: KN, LPM, VN. Cytoarchitectonic analysis and atlas-based mapping: KN, LPM, VN, LSA. Research funding procurement: AMK. Research supervision and training: AMK, KN, VN. Conference presenters: KN, LPM. Manuscript preparation in current form: KN, AMK with final checks from all co-authors. All authors have read and agreed to this version of the manuscript.

## Data Availability Statement

All relevant protocols and data are contained within the article and supplemental materials.

## Acknowledgments

All *BM4.0* atlas maps were modified and/or reproduced with permission under the conditions outlined by a Creative Commons BY-NC 4.0 license. Adobe Illustrator and Adobe Photoshop are either registered trademarks or trademarks of Adobe in the United States and/or other countries.

## Abbreviations

Abbreviations and standard terms follow those of Swanson (2018). When available, the references at the ends of standard terms refer to the first use of the terms as defined. See *Section 2.1* of this study for details.

AAA: anterior amygdalar area (Gurdjian 1928)
ACAd: anterior cingulate area dorsal part (Krettek & Price, 1977)
ACB: nucleus accumbens (Ziehen, 1897-1901)
aco: olfactory limb of the anterior commissure (>1840)
act: temporal limb of the anterior commissure (>1840)
AD: anterodorsal thalamic nucleus (>1840)
ADP: anterodorsal preoptic nucleus (Simerly et al., 1984)
AHA: anterior hypothalamic area (>1840)
AHNa: anterior hypothalamic area, anterior part (>1840)
AHNc: anterior hypothalamic area, central part (>1840)
AHNd: anterior hypothalamic area, dorsal part (>1840)
AHNp: anterior hypothalamic area, posterior part (>1840)
AM: anteromedial thalamic nucleus (>1840)
AMd: anteromedial thalamic nucleus dorsal part (Canteras & Swanson, 1992)
AMv: anteromedial thalamic nucleus ventral part (Canteras & Swanson, 1992)
APN: anterior pretectal nucleus (>1840)
AQo: opening of cerebral aqueduct (>1840)
ARH: arcuate hypothalamic nucleus (>1840)
ATN: anterior thalamic nuclei (>1840)
AV: anteroventral thalamic nucleus (>1840)
AVP: anteroventral preoptic nucleus (>1840)
AVPV: anteroventral periventricular nucleus (>1840)
BA: bed nucleus of accessory olfactory tract (Scalia & Winans, 1975)
BAC: bed nucleus of anterior commissure (Gurdjian, 1925)
BSM: bed nucleus of medullary stria (Risold & Swanson, 1995)
BST: bed nuclei of terminal stria (Gurdjian, 1925)
BSTa: bed nuclei of terminal stria anterior division (Ju & Swanson, 1989)
BSTal: bed nuclei of terminal stria anterolateral area (Swanson, 2004)
BSTam: bed nuclei of terminal stria anteromedial area (Dong & Swanson, 2006b)
BSTdm: bed nuclei of terminal stria dorsomedial nucleus (Ju & Swanson, 1989)
BSTfu: bed nuclei of terminal stria fusiform nucleus (Ju & Swanson, 1989)
BSTif: bed nuclei of terminal stria interfascicular nucleus (Ju & Swanson, 1989)
BSTju: bed nuclei of terminal stria juxtacapsular nucleus (McDonald, 1983)
BSTmg: bed nuclei of terminal stria magnocellular nucleus (Ju & Swanson, 1989)
BSTp: bed nuclei of terminal stria posterior division (Ju & Swanson, 1989)
BSTpr: bed nuclei of terminal stria principal nucleus (Ju & Swanson, 1989)
BSTrh: bed nuclei of terminal stria rhomboid nucleus (Ju & Swanson, 1989)
BSTse: bed nuclei of terminal stria striatal extension (Ju & Swanson, 1989)
BSTsz: bed nuclei of terminal stria principal nucleus cell-sparse zone (Ju & Swanson, 1989)
BSTtr: bed nuclei of terminal stria transverse nucleus (Ju & Swanson, 1989)
BSTv: bed nuclei of terminal stria ventral nucleus (Ju & Swanson, 1989)
CEAc: central amygdalar nucleus capsular part (McDonald, 1982)
CEAm: central amygdalar nucleus medial part (McDonald, 1982)
CL: central lateral thalamic nucleus (Rioch, 1929)
CM: central medial thalamic nucleus (Rioch, 1929)
CNG: cingulate region (Brodmann, 1909)
COAa: cortical amygdalar area anterior part (>1840)
COApm: cortical amygdalar area posterior part medial zone (>1840)
CP: caudoputamen (Heimer & Wilson, 1975)
cpd: cerebral peduncle (Tarin, 1753)
DMH: dorsomedial hypothalamic nucleus (>1840)
DMHa: dorsomedial hypothalamic nucleus anterior part (>1840)
DMHp: dorsomedial hypothalamic nucleus posterior part (>1840)
DMHv: dorsomedial hypothalamic nucleus ventral part (>1840)
ec: external capsule (Burdach, 1822)
em: external medullary lamina (>1840)
FF: fields of Forel (>1840)
frf: radiation of corpus callosum frontal forceps (>1840)
FS: striatal fundus (>1840)
GPl: lateral globus pallidus (>1840)
GPm: medial globus pallidus (>1840)
hbc: habenular commissure (>1840)
hitt: thalamic habenulo-interpeduncular tract (Swanson, 2015)
HY: hypothalamus (Kuhlenbeck, 1927)
HYa: hypothalamus anterior level (>1840)
HYl: lateral hypothalamic zone (Nauta & Haymaker, 1969)
HYm: medial hypothalamic zone (Nauta & Haymaker, 1969)
HYp: periventricular hypothalamic zone (Nauta & Haymaker, 1969)
HYpr: hypothalamic preoptic level (>1840)
hys: hypothalamic sulcus (>1840)
I: internuclear hypothalamic area (Swanson, 2004)
IAD: interanterodorsal thalamic nucleus (>1840)
IAM: interanteromedial thalamic nucleus (>1840)
IB: interbrain (Baer, 1837)
ILA: infralimbic area (Rose & Woolsey, 1948)
ILM: intralaminar thalamic nuclei (>1840)
IMD: intermediodorsal thalamic nucleus (>1840)
im: thalamic internal medullary lamina (>1840)
int: internal capsule (Burdach, 1822)
IT: interthalamic adhesion (>1840)
LD: lateral dorsal thalamic nucleus (>1840)
LH: lateral habenula (>1840)
LHA: lateral hypothalamic area (Nissl, 1913)
LHAad: lateral hypothalamic area anterior group anterior region dorsal zone (Swanson, 2004)
LHAag: lateral hypothalamic area anterior group (Swanson et al., 2005)
LHAai: lateral hypothalamic area anterior region anterior region intermediate zone (Swanson, 2004)
LHAav: lateral hypothalamic area anterior group anterior region ventral zone (Swanson, 2004)
LHAd: lateral hypothalamic area middle group lateral tier dorsal region (Swanson, 2004)
LHAjd: lateral hypothalamic area middle group medial tier juxtadorsomedial region (Swanson, 2004)
LHAjp: lateral hypothalamic area middle group medial tier juxtaparaventricular region (Swanson, 2004)
LHAjv: lateral hypothalamic area middle group medial tier juxtaventromedial region (Swanson, 2004)
LHAjvd: lateral hypothalamic area middle group medial tier juxtaventromedial region dorsal zone (Swanson, 2004)
LHAjvv: lateral hypothalamic area middle group medial tier juxtaventromedial region ventral zone (Swanson, 2004)
LHAl: lateral hypothalamic area middle group lateral tier (Swanson et al., 2005)
LHAm: lateral hypothalamic area middle group medial tier (Swanson et al., 2005)
LHAma: lateral hypothalamic area middle group lateral tier ventral region magnocellular nucleus (Paxinos & Watson, 1986)
LHAmg: lateral hypothalamic area middle group (Swanson et al., 2005)
LHAp: lateral hypothalamic area posterior group posterior region (Swanson, 2004)
LHApc: lateral hypothalamic area middle group lateral tier ventral region parvicellular region (Swanson, 2004)
LHApf: lateral hypothalamic area middle group perfironical tier (Swanson et al., 2005)
LHApg: lateral hypothalamic area posterior group (Swanson et al., 2005)
LHAs: lateral hypothalamic area middle group perifornical tier suprafornical region (Swanson, 2004)
LHAsf: lateral hypothalamic area middle group perifornical tier subfornical region (Swanson, 2004)
LHAsfa: lateral hypothalamic area middle group perifornical tier subfornical region anterior zone (Swanson, 2004)
LHAsfp: lateral hypothalamic area middle group perifornical tier subfornical region posterior zone (Swanson, 2004)
LHAsfpm: lateral hypothalamic area middle group perifornical tier subfornical region posterior zone premammillary subzone (Swanson, 2004)
LHAv: lateral hypothalamic area middle group lateral tier ventral region (Swanson et al., 2005)
LHAvl: lateral hypothalamic area middle group lateral tier ventral region lateral zone (Swanson, 2004)
LHAvm: lateral hypothalamic area middle group lateral tier ventral region medial zone (Swanson, 2004)
LM: lateral mammillary nucleus (Gudden, 1881)
LP: lateral posterior thalamic nucleus (>1840)
LPO: lateral preoptic area (>1840)
LSr: lateral septal nucleus rostral (rostroventral) part (Risold & Swanson, 1997)
LSr.m.v.c: lateral septal nucleus rostral (rostroventral) part medial zone ventral region caudal domain (Risold & Swanson, 1997)
LSr.vl.v: lateral septal nucleus rostral (rostroventral) part ventrolateral zone ventral region (Risold & Swanson, 1997)
LSv: lateral septal nucleus ventral part (Risold & Swanson, 1997)
MA: magnocellular nucleus (Swanson, 2004)
mct: medial corticohypothalamic tract (Gurdjian, 1927)
MDc: mediodorsal thalamic nucleus central part (>1840)
MDl: mediodorsal thalamic nucleus lateral part (>1840)
MDm: mediodorsal thalamic nucleus medial part (>1840)
ME: median eminence (Tilney, 1936)
MEex: median eminence external lamina (>1840)
MEin: median eminence internal lamina (>1840)
MEAad: medial amygdalar nucleus anterodorsal part (>1840)
MEAav: medial amygdalar nucleus anteroventral part (>1840)
MEApv: medial amygdalar nucleus posteroventral part (>1840)
MEApd: medial amygdalar nucleus posterodorsal part(>1840)
MEPO: median preoptic nucleus (Loo, 1931)
mfb: medial forebrain bundle (Edinger, 1893)
mlt: thalamic medial lemniscus (Swanson, 2015)
MM: medial mammillary nucleus (Gudden, 1881)
MMme: medial mammillary nucleus median part (>1840)
MMpr: medial mammillary nucleus principal part (Swanson, 2018)
MOs: secondary somatomotor areas (>1840)
mph: hypothalamic mammillary peduncle (Swanson, 2018)
MPN: medial preoptic nucleus (Gurdjian, 1927)
MPNc: medial preoptic nucleus central part (Simerly et al., 1984)
MPNl: medial preoptic nucleus lateral part (Simerly et al., 1984)
MPNm: medial preoptic nucleus medial part (Simerly et al., 1984)
MPO: medial preoptic area (>1840)
MRNm: midbrain reticular nucleus magnocellular part (Swanson, 2004)
MS: medial septal nucleus (>1840)
mtgh: hypothalamic mammillotegmental tract (Swanson, 2015)
mtt: mammillothalamic tract (Kölliker, 1896)
mtth: hypothalamic mammillothalamic tract (Swanson, 2018)
mttt: thalamic mammillothalamic tract (Swanson, 2015)
NC: nucleus circularis (>1840)
NDB: diagonal band nucleus (>1840)
NId: nucleus incertus diffuse part (Goto et al., 2001)
NLOT: nucleus of the lateral olfactory tract (Swanson & Petrovich, 1998)
NPC: nucleus of posterior commissure (>1840)
och: optic chiasm (Galen, c173)
opth: hypothalamic optic tract (Swanson, 2015)
optt: thalamic optic tract (Swanson, 2015)
ORBv: ventral orbital area (Krettek & Price, 1977)
OT: olfactory tubercle (Calleja, 1893)
OV: vascular organ of lamina terminalis (>1840)
PA: posterior amygdalar nucleus (Canteras et al., 1992a)
PAG: periaqueductal gray (>1840)
PAGdl: periaqueductal gray dorsolateral column (Carrive et al., 1997)
PAGm: periaqueductal gray medial division (Beitz, 1985)
PAGrl: periaqueductal gray rostrolateral division (Swanson, 1998)
PAGrm: periaqueductal gray rostromedial division (Swanson, 1998)
PAGvl: periaqueductal gray ventrolateral column (Carrive et al., 1997)
PCN: paracentral thalamic nucleus (Gurdjian, 1927)
PD: posterodorsal preoptic nucleus (Simerly et al., 1984)
pdl: peduncular loop (Gratiolet, 1857)
PF: parafascicular nucleus (Vogt, 1909)
PH: posterior hypothalamic nucleus (>1840)
PL: prelimbic area (Brodmann, 1909)
pm: principal mammillary tract (Kölliker, 1896)
PMd: dorsal mammillary nucleus (>1840)
PMv: ventral premammillary nucleus (>1840)
PO: posterior thalamic complex (>1840)
pofs: septal postcommissural fornix (Swanson, 2015)
pofh: hypothalamic postcommissural fornix (Swanson, 2015)
PR: perireuniens nucleus (Brittain, 1988)
PRC: periaqueductal gray precommissural nucleus (Paxinos & Watson, 1986)
PS: parastrial nucleus (Simerly et al., 1984)
PSCH: suprachiasmatic preoptic nucleus (Simerly et al., 1984)
PST: preparasubthalamic nucleus (Swanson, 2004)
PSTN: parasubthalamic nucleus (Wang & Zhang, 1995)
PT: paratenial nucleus (>1840)
PVa: periventricular hypothalamic nucleus anterior part (Swanson, 2018)
PVaa: periventricular hypothalamic nucleus anterior part anterior zone (Swanson, 2018)
PVH: paraventricular hypothalamic nucleus (>1840)
PVHam: paraventricular hypothalamic nucleus magnocellular division anterior magnocellular part (Swanson & Kuypers, 1980)
PVHap: paraventricular hypothalamic nucleus parvicellular division anterior parvicellular part (>1840)
PVHdp: paraventricular hypothalamic nucleus descending division dorsal parvicellular part (>1840)
PVHf: paraventricular hypothalamic nucleus descending division forniceal part (>1840)
PVHlp: paraventricular hypothalamic nucleus descending division lateral parvicellular part (>1840)
PVHmpd: paraventricular hypothalamic nucleus parvicellular division medial parvicellular part dorsal zone (Simmons & Swanson, 2008)
PVHmpdl: paraventricular hypothalamic nucleus parvicellular medial parvicellular part dorsal zone lateral wing (Simmons & Swanson, 2008)
PVHmpv: paraventricular hypothalamic nucleus descending division medial parvicellular part ventral zone (>1840)
PVHpml: paraventricular hypothalamic nucleus magnocellular division posterior magnocellular part lateral zone (>1840)
PVHpmm: paraventricular hypothalamic nucleus magnocellular division posterior magnocellular part medial zone (>1840)
PVHpv: paraventricular hypothalamic nucleus parvicellular division periventricular part (>1840)
PVi: paraventricular hypothalamic nucleus anterior part intermediate zone (Swanson, 2018)
PVp: periventricular hypothalamic nucleus posterior part (>1840)
PVpo: periventricular hypothalamic nucleus anterior part preoptic zone (Swanson, 2018)
PVT: paraventricular thalamic nucleus (>1840)
RCH: lateral hypothalamic area anterior group retrochiasmatic area (>1840)
REcd: nucleus reuniens caudal division dorsal part (Risold et al., 1997)
REcm: nucleus reuniens caudal division median part (Risold et al., 1997)
REcp: nucleus reuniens caudal division posterior part (Risold et al., 1997)
REr: nucleus reuniens rostral division (Risold et al., 1997)
REra: nucleus reuniens rostral division anterior part (Risold et al., 1997)
RErd: nucleus reuniens rostral division dorsal part (Risold et al., 1997)
RErl: nucleus reuniens rostral division lateral part (Risold et al., 1997)
RErm: nucleus reuniens rostral division median part (Risold et al., 1997)
RErv: nucleus reuniens rostral division ventral part (Risold et al., 1997)
RH: rhomboid nucleus (Cajal, 1904)
ri: rhinal incisure (>1840)
RT: reticular thalamic nucleus (>1840)
SBPV: subparaventricular zone (Watts et al., 1987)
SCH: suprachiasmatic nucleus (Spiegel & Zwieg, 1919)
SCO: subcommissural organ (>1840)
SEZ: subependymal zone (>1840)
SF: septofimbrial nucleus (>1840)
SI: innominate substance (Schwalbe, 1881)
sm: medullary stria (Wenzel & Wenzel, 1812)
sme: epithalamic medullary stria (Swanson, 2015)
smd: supramammillary decussation (>1840)
smh: hypothalamic medullary stria (Swanson, 2015)
SMT: submedial thalamic nucleus (>1840)
SNr: substantia nigra reticular part (Sano, 1910)
SO: supraoptic nucleus (Lenhossék, 1887)
SOp: supraoptic nucleus principal part (Swanson, 2018)
SOr: supraoptic nucleus retrochiasmatic part (>1840)
SPFm: subparafascicular nucleus magnocellular part (>1840)
SPFpl: subparafascicular nucleus parvicellular part lateral division (>1840)
SPFpm: subparafascicular nucleus parvicellular part medial division (>1840)
st: terminal stria (Wenzel & Wenzel, 1812)
ste: endbrain terminal stria (Swanson, 2015)
STN: subthalamic nucleus (>1840)
SUM: supramammillary nucleus (>1840)
SUMl: supramammillary nucleus lateral part (Swanson, 1982)
SUMm: supramammillary nucleus medial part (Swanson, 1982)
sup: supraoptic decussations (>1840)
suph: hypothalamic supraoptic decussations (Swanson, 2018)
TEP: temporal pole (Broca, 1878)
TH: thalamus (His, 1893a)
THe: epithalamus (His, 1893b)
THv: ventral part of thalamus (Herrick, 1910)
TMd: tuberomammillary nucleus dorsal part (Köhler et al., 1985)
TMv: tuberomammillary nucleus ventral part (Köhler et al., 1985)
TRS: triangular septal nucleus (>1840)
TTd: tenia tecta dorsal part (Swanson, 1992)
TTv: tenia tecta ventral part (Swanson, 1992)
TU: lateral hypothalamic area middle group lateral tier tuberal nucleus (>1840)
TUi: lateral hypothalamic area middle group lateral tier tuberal nucleus intermediate part (Swanson, 2004)
TUl: lateral hypothalamic area middle group lateral tier tuberal nucleus lateral part (Swanson, 2004)
TUsv: lateral hypothalamic area middle group lateral tier tuberal nucleus subventromedial part (Swanson, 2004)
TUte: lateral hypothalamic area middle group lateral tier tuberal nucleus terete part (Petrovich et al., 2001)
V3h: hypothalamic part of third ventricle principal part (Swanson, 2018)
V3i: third ventricle infundibular recess (>1840)
V3m: third ventricle mammillary recess (>1840)
V3p: third ventricle preoptic recess (>1840)
V3r: roof of third ventricle (>1840)
V3t: thalamic part of third ventricle principal part (Swanson, 2018)
VAL: ventral anterior-lateral thalamic complex (>1840)
VENT: ventral thalamic nuclei (>1840)
VLP: ventrolateral preoptic nucleus (Sherin et al., 1998)
vlt: ventrolateral hypothalamic tract (Swanson, 2004)
VM: ventral medial thalamic nucleus (>1840)
VMHa: ventromedial hypothalamic nucleus anterior part (>1840)
VMHc: ventromedial hypothalamic nucleus central part (>1840)
VMHdm: ventromedial hypothalamic nucleus dorsomedial part (>1840)
VMHvl: ventromedial hypothalamic nucleus ventrolateral part (>1840)
VPLpc: ventral posterolateral thalamic nucleus parvicellular part (>1840)
VPLpr: ventral posterolateral thalamic nucleus principal part (Swanson, 2004)
VPMpc: ventral posteromedial thalamic nucleus parvicellular part (>1840)
VPMpr: ventral posteromedial thalamic nucleus principal part (Swanson, 2004)
VTA: ventral tegmental area (Tsai, 1925)
ZI: zona incerta (>1840)
ZIda: zona incerta dopaminergic group (>1840)

## References

Abrahamson, E. E., & Moore, R. Y. (2001). The posterior hypothalamic area: chemoarchitecture and afferent connections. Brain Res, 889(1–2), 1–20. 10.1016/s0006-8993(00)03015-8

Akhter, F., Haque, T., Sato, F., Kato, T., Ohara, H., Fujio, T., Tsutsumi, K., Uchino, K., Sessle, B. J., & Yoshida, A. (2014). Projections from the dorsal peduncular cortex to the trigeminal subnucleus caudalis (medullary dorsal horn) and other lower brainstem areas in rats. Neuroscience, 266, 23–37. 10.1016/j.neuroscience.2014.01.046

Alcaraz, F., Marchand, A. R., Courtand, G., Coutureau, E., & Wolff, M. (2016). Parallel inputs from the mediodorsal thalamus to the prefrontal cortex in the rat. Eur J Neurosci, 44(3), 1972–1986. 10.1111/ejn.13316

Alexander, G. E., Crutcher, M. D., & DeLong, M. R. (1990). Basal ganglia-thalamocortical circuits: parallel substrates for motor, oculomotor, “prefrontal” and “limbic” functions. Prog Brain Res, 85, 119–146. 10.1016/S0079-6123(08)62678-3

Allen, G. V., & Hopkins, D. A. (1989). Mamillary body in the rat: topography and synaptology of projections from the subicular complex, prefrontal cortex, and midbrain tegmentum. J Comp Neurol, 286(3), 311–336. 10.1002/cne.902860303

Baer, K.E. von (1828–1837) Über Entwickelungsgeschichte der Thiere. Beobachtung und Reflexion, (Bornträger; Koch, Königsberg). Available online: https://archive.org/details/berentwickelun01baer/page/n5/mode/2up.

Balfour, M. E., Brown, J. L., Yu, L., & Coolen, L. M. (2006). Potential contributions of efferents from medial prefrontal cortex to neural activation following sexual behavior in the male rat. Neuroscience, 137(4), 1259– 1276. 10.1016/j.neuroscience.2005.11.013

Barbier, M., Fellmann, D., & Risold, P.-Y. (2018a). Characterization of McDonald’s intermediate part of the central nucleus of the amygdala in the rat. J Comp Neurol, 526(14), 2165–2186. 10.1002/cne.24470

Barbier, M., Fellmann, D., & Risold, P.-Y. (2018b). Morphofunctional organization of the connections from the medial and intermediate parts of the central nucleus of the amygdala into distinct divisions of the lateral hypothalamic area in the rat. Front Neurol, 9, 688. 10.3389/fneur.2018.00688

Barbier, M., Chometton, S., Pautrat, A., Miguet-Alfonsi, C., Datiche, F., Gascuel, J., Fellman, D., Peterschmitt, Y., Coizet, V., & Risold, P.-Y. (2020). A basal ganglia-like cortical-amygdalar-hypothalamic network mediates feeding behavior. Proc Natl Acad Sci U S A, 117(27), 15967–15976. 10.1073/pnas.2004914117

Beitz, A. J. (1985). The midbrain periaqueductal gray in the rat. I. Nuclear volume, cell number, density, orientation, and regional subdivisions. J Comp Neurol, 237(4), 445–459. 10.1002/cne.902370403

Berendse, H. W., & Groenewegen, H. J. (1991). Restricted cortical termination fields of the midline and intralaminar thalamic nuclei in the rat. Neuroscience, 42(1), 73–102. 10.1016/0006-8993(90)91570-7

Biag, J., Huang, Y., Gou, L., Hintiryan, H., Askarinam, A., Hahn, J. D., Toga, A. W., & Dong, H.-W. (2011). Cyto- and chemoarchitecture of the hypothalamic paraventricular nucleus in the C57BL/6J male mouse: A study of immunostaining and multiple fluorescent tract tracing. J Comp Neurol, 520(1), 6–30. 10.1002/cne.22698

Bienkowski, M. S., Bowman, I., Song, M. Y., Gou, L., Ard, T., Cotter, K., Zhu, M., Benavidez, N. L., Yamashita, S., Abu-Jaber, J., Azam, S., Lo, D., Foster, N. N., Hintiryan, H., & Dong, H.-W. (2018). Integration of gene expression and brain-wide connectivity reveals the multiscale organization of mouse hippocampal networks. Nat Neurosci, 21(11), 1628–1643. 10.1038/s41593-018-0241-y

Bota, M., Dong, H.-W., & Swanson, L. W. (2003). From gene networks to brain networks. Nat Neurosci, 6(8), 795–799. 10.1038/nn1096

Bota, M., Sporns, O., & Swanson, L. W. (2015). Architecture of the cerebral cortical association connectome underlying cognition. Proc Natl Acad Sci U S A, 112(16), E2093–E2101. 10.1073/pnas.1504394112

Brittain, D. A. (1988). The efferent connections of the infralimbic cortex in the rat. Doctoral dissertation. University of California, San Diego.

Broca, P. P. (1878). Nomenclature cérébrale: dénomination et subdivision des hémisphères et des anfractuosités de la surface. Revue d’Anthropologie, 1, 193–236.

Brodmann, K. (1909). *Vergleichende Lokalisationslehre der Grosshirnrinde in ihren Prinzipien Dargestellt auf Grund des Zellenbaues* (Barth, Leipzig).

Burdach, K. F. (1819, 1822, 1826). Vom Baue und Leben des Gehirns (Dyk’schen Buchhandlung, Leipzig).

Buzsáki, G. (2019). The brain from inside out. Oxford: Oxford University Press. https://psycnet.apa.org/doi/10.1093/oso/9780190905385.001.0001

Cajal, S.R. y (1899–1904). Textura del Sistema Nervioso del Hombre y de los Vertebrados: Estudios sobre el Plan Estructural y Composición de los Centros Serviosos Adicionados de Consideraciones Fisiológicas Fundadas en los Nuevos Descubrimientos, 2 Vols. (Moya, Madrid).

Calabrese, E., Badea, A., Cofer, G., Qi, Y., & Johnson, G. A. (2015). A diffusion MRI tractography connectome of the mouse brain atlas and comparison with neuronal tracer data. Cereb Cortex, 25(11), 4628–37. 10.1093/cercor/bhv121

Calleja, C. (1893). *La Región Olfatoria del Cerebro* (Moya, Madrid).

Canteras, N. S., Ribeiro-Barbosa, E. R., Goto, M., Cipolla-Neto, J., & Swanson, L. W. (2011). The retinohypothalamic tract: comparison of axonal projection patterns from four major targets. Brain Res Rev, 65(2), 150–183. 10.1016/j.brainresrev.2010.09.006

Canteras, N. S., Shammah-Lagnado, S. J., Silva, B. A., & Ricardo, J. A. (1990). Afferent connections of the subthalamic nucleus: a combined retrograde and anterograde horseradish peroxidase study in the rat. Brain Res, 513(1), 43–59. 10.1016/0006-8993(90)91087-w

Canteras, N.S., Simerly, R. B., & Swanson, L.W. (1992a). The connections of the posterior nucleus of the amygdala. J Comp Neurol, 324(2), 143–179. 10.1002/cne.903240203

Canteras, N. S., Simerly, R. B., & Swanson, L. W. (1992b). Projections of the ventral premammillary nucleus. J Comp Neurol, 324(2), 195–212. 10.1002/cne.903240205

Canteras, N. S., Simerly, R. B., & Swanson, L. W. (1994). Organization of projections from the ventromedial nucleus of the hypothalamus: a *Phaseolus vulgaris*-leucoagglutinin study in the rat. J Comp Neurol, 348(1), 41–79. 10.1002/cne.903480103

Canteras, N. S., Simerly, R. B., & Swanson, L. W. (1995). Organization of projections from the medial nucleus of the amygdala: a PHAL study in the rat. J Comp Neurol, 360(2), 213–245. 10.1002/cne.903600203

Canteras, N.S. & Swanson, L.W. (1992). The dorsal premammillary nucleus: an unusual component of the mammillary body. Proc Natl Acad Sci U S A, 89(21), 10089–10093. 10.1073/pnas.89.21.10089

Carrive, P., Leung, P., Harris, J., & Paxinos, G. (1997). Conditioned fear to context is associated with increased Fos expression in the caudal ventrolateral region of the midbrain periaqueductal gray. Neuroscience, 78(1), 165–177. 10.1016/s0306-4522(97)83047-3

Castro, D. C., & Berridge, K. C. (2017). Opioid and orexin hedonic hotspots in rat orbitofrontal cortex and insula. Proc Natl Acad Sci U S A, 114(43), E9125–E9134. 10.1073/pnas.1705753114

Chen, S., & Su, H. S. (1990). Afferent connections of the thalamic paraventricular and parataenial nuclei in the rat—a retrograde tracing study with iontophoretic application of Fluoro-Gold. Brain Res, 522(1), 1–6. 10.1016/0006-8993(90)91570-7

Chiba, T., & Murata, Y. (1985). Afferent and efferent connections of the medial preoptic area in the rat: a WGA-HRP study. Brain Res Bull, 14(3), 261–272. 10.1016/0361-9230(85)90091-7

Chometton, S., Pedron, S., Peterschmitt, Y., Van Waes, V., Fellmann, D., & Risold, P.-Y. (2015). A premammillary lateral hypothalamic nuclear complex responds to hedonic but not aversive tastes in the male rat. Brain Struct Funct, 221(4), 2183–2208. 10.1007/s00429-015-1038-3

Chometton, S., Charrière, K., Bayer, L., Houdayer, C., Franchi, G., Poncet, F., Fellmann, D., & Risold, P.-Y. (2017). The rostromedial zona incerta is involved in attentional processes while adjacent LHA responds to arousal: c-Fos and anatomical evidence. Brain Struct Funct, 222(6), 2507–2525. 10.1007/s00429-016-1353-3

Chou, X.-L., Wang, X., Zhang, Z.-G., Shen, L., Zingg, B., Huang, J., Zhong, W., Mesik, L., Zhang, L. I., & Tao, H. W. (2018). Inhibitory gain modulation of defense behaviors by zona incerta. Nat Commun, 9(1), 1151. 10.1038/s41467-018-03581-6

Condé, F., Maire-Lepoivre, E., Audinat, E., & Crépel, F. (1995). Afferent connections of the medial frontal cortex of the rat. II. Cortical and subcortical afferents. J Comp Neurol, 352(4), 567–593. 10.1002/cne.903520407

Cornwall, J., & Phillipson, O. T. (1988). Afferent projections to the parafascicular thalamic nucleus of the rat, as shown by the retrograde transport of wheat germ agglutinin. Brain Res Bull, 20(2), 139–150. 10.1016/0361-9230(88)90171-2

Cornwall, J., Cooper, J. D., & Phillipson, O. T. (1990). Projections to the rostral reticular thalamic nucleus in the rat. Exp Brain Res, 80(1), 157–171. 10.1007/bf00228857

De Haro, B. (2015). *Structural organization of the connections between neurons of the paraventricular and lateral hypothalamic regions in the adult* *male* *rat*. Master’s Thesis. The University of Texas at El Paso. *Open Access Theses & Dissertations*, 1028. https://scholarworks.utep.edu/open_etd/1028

de Lima, M. A., Baldo, M. V., & Canteras, N. S. (2017). A role for the anteromedial thalamic nucleus in the acquisition of contextual fear memory to predatory threats. Brain Struct Funct, 222(1), 113–129. 10.1007/s00429-016-1204-2

Domesick, V. B. (1969). Projections from the cingulate cortex in the rat. Brain Res, 12(2):296–320. 10.1016/0006-8993(69)90002-x

Dong, H.-W., & Swanson, L. W. (2003). Projections from the rhomboid nucleus of the bed nuclei of the stria terminalis: implications for cerebral hemisphere regulation of ingestive behaviors. J Comp Neurol, 463(4), 434–472. 10.1002/cne.10758

Dong, H.-W., & Swanson, L. W. (2004). Organization of axonal projections from the anterolateral area of the bed nuclei of the stria terminalis. J Comp Neurol, 468(2), 277–298. 10.1002/cne.10949

Dong, H.-W., & Swanson, L. W. (2006a). Projections from bed nuclei of the stria terminalis, dorsomedial nucleus: implications for cerebral hemisphere integration of neuroendocrine, autonomic, and drinking responses. J Comp Neurol, 494(1), 75–107. 10.1002/cne.20790

Dong, H.-W. & Swanson, L.W. (2006b). Projections from bed nuclei of the stria terminalis, anteromedial area: cerebral hemisphere integration of neuroendocrine, autonomic, and behavioral aspects of energy balance. J Comp Neurol, 494(1), 142–178. 10.1002/cne.20788

Edinger, L. (1893). *Vorlesungen über den Bau der nervösen Centralorgane des Menschen und der Thier*e (Vogel, Leipzig).

Eickhoff, S. B., Constable, R. T., & Thomas Yeo, B. T. (2018). Topographic organization of the cerebral cortex and brain cartography. NeuroImage, 170, 332–347.

Espinoza, S.G. & Thomas, T.C. (1983). Retinotopic organization of striate and extrastriate visual cortex in the hooded rat. Brain Research, 272, 137–144.

Fisk, G. D., & Wyss, J. M. (2000). Descending projections of infralimbic cortex that mediate stimulation-evoked changes in arterial pressure. Brain Res, 859(1), 83–95. 10.1016/s0006-8993(00)01935-1

Fujiyama, F., Unzai, T., & Karube, F. (2019). Thalamostriatal projections and striosome-matrix compartments. Neurochem Int, 125, 67–73. 10.1016/j.neuint.2019.01.024

Gabbott, P. L., Warner, T. A., Jays, P. R., Salway, P., & Busby, S. J. (2005). Prefrontal cortex in the rat: projections to subcortical autonomic, motor, and limbic centers. J Comp Neurol, 492(2), 147–177. 10.1002/cne.20738

55. Galen, C. (c173). On the Usefulness of the Parts of the Body. Translated from the Greek with an Introduction and Commentary by M. T. May (Cornell University Press, Ithaca), 1968.

Gerfen, C. R., & Sawchenko, P. E. (1984). An anterograde neuroanatomical tract tracing method that shows the detailed morphology of neurons, their axons and terminals: immunohistochemical localization of an axonally transported plant lectin, *Phaseolus vulgaris* leucoagglutinin (PHAL). Brain Res, 1644, 42–45. 10.1016/j.brainres.2015.12.040

Goto, M., Swanson, L. W., & Canteras, N. S. (2001). Connections of the nucleus incertus. J Comp Neurol, 438(1), 86–122. 10.1002/cne.1303

Goto, M., & Swanson, L. W. (2004). Axonal projections from the parasubthalamic nucleus. J Comp Neurol, 469(4), 581–607. 10.1002/cne.11036

Goto, M., Canteras, N. S., Burns, G., & Swanson, L. W. (2005). Projections from the subfornical region of the lateral hypothalamic area. J Comp Neurol, 493(3), 412–438. 10.1002/cne.20764

Gratiolet (1857): see Leuret, F. & Gratiolet, P.

Grill, H. J., & Norgren, R. (1978). Neurological tests and behavioral deficits in chronic thalamic and chronic decerebrate rats. Brain Res, 143(2), 299–312.

Groenewegen, H. J. (1988). Organization of the afferent connections of the mediodorsal thalamic nucleus in the rat, related to the mediodorsal-prefrontal topography. Neuroscience, 24(2), 379–431. 10.1016/0306-4522(88)90339-9

Gudden, B. von (1881). Beitrag zur Kentnis des Corpus mammillare und der sogenannten Schenkel des Fornix. Archiv für Psychiatrie und Nervenkrankheiten, 11, 428–452.

Gurdjian, E. S. (1925). Olfactory connections in the albino rat, with special reference to the stria medullaris and the anterior commissure. J Comp Neurol, 38, 127–163.

Gurdjian, E. S. (1927). The diencephalon of the albino rat. J Comp Neurol, 43, 1–114.

Gurdjian, E. S. (1928). The corpus striatum of the rat. J Comp Neurol, 45, 249–281.

Hahn, J. D. (2010). Comparison of melanin-concentrating hormone and hypocretin/orexin peptide expression patterns in a current parceling scheme of the lateral hypothalamic zone. Neurosci Lett, 468(1), 12–17. 10.1016/j.neulet.2009.10.047

Hahn, J. D., & Swanson, L. W. (2010). Distinct patterns of neuronal inputs and outputs of the juxtaparaventricular and suprafornical regions of the lateral hypothalamic area in the male rat. Brain Res Rev, 64(1), 14–103. 10.1016/j.brainresrev.2010.02.002

Hahn, J. D., & Swanson, L. W. (2012). Connections of the lateral hypothalamic area juxtadorsomedial region in the male rat. J Comp Neurol, 520(9), 1831–1890. 10.1002/cne.23064

Hahn, J. D., & Swanson, L. W. (2015). Connections of the juxtaventromedial region of the lateral hypothalamic area in the male rat. Front Syst Neurosci, 9, 66. 10.3389/fnsys.2015.00066

Hahn, J. D., Sporns, O., Watts, A. G., & Swanson, L. W. (2019). Macroscale intrinsic network architecture of the hypothalamus. Proc Natl Acad U S A, 116(16), 8018–8027. 10.1073/pnas.1819448116

Hahn, J. D., Gao, L., Boesen, T., Gou, L., Hintiryan, H., & Dong, H.-W. (2022). Macroscale connections of the mouse lateral preoptic area and anterior lateral hypothalamic area. J Comp Neurol, 530(13), 2254–2285. 10.1002/cne.25331

Heimer, L. & Wilson, R.D. (1975). The subcortical projections of the allocortex: similarities in the neural associations of the hippocampus, the piriform cortex, and the neocortex. In: Santini, M. (Ed.) Golgi Centennial Symposium Proceedings (Raven Press, New York), pp. 177–193.

Herrick, C.J. (1910). The morphology of the forebrain in amphibia and reptilia. J Comp Neurol Psychol, 20, 413–547. 10.1002/cne.920200502

Hintiryan, H., Foster, N. N., Bowman, I., Bay, M., Song, M. Y., Gou, L., Yamashita, S., Bienkowski, M. S., Zingg, B., Zhu, M., Yang, X. W., Shih, J. C., Toga, A. W., & Dong, H.-W. (2016). The mouse cortico-striatal projectome. Nat Neurosci 19(8), 1100–1114. 10.1038/nn.4332

His, W. (1893a). Ueber das frontale Ende des Gehirnrohres. *Archiv für Anatomie und Entwickelungsgeschichte* [no volume indicated] pp. 157–171.

His, W. (1893b). Vorschläge zur Eintheilung des Gehirns. *Archiv für Anatomie und Entwickelungsgeschichte* [no volume indicated] pp. 172–179.

Hoover, W. B., & Vertes, R. P. (2007). Anatomical analysis of afferent projections to the medial prefrontal cortex in the rat. Brain Struct Funct, 212(2), 149–179. 10.1007/s00429-007-0150-4

Hurley, K. M., Herbert, H., Moga, M. M., & Saper, C. B. (1991). Efferent projections of the infralimbic cortex of the rat. J Comp Neurol, 308(2), 249–276. 10.1002/cne.903080210

Jones, B. F., Groenewegen, H. J., & Witter, M. P. (2005). Intrinsic connections of the cingulate cortex in the rat suggest the existence of multiple functionally segregated networks. Neuroscience, 133(1), 193–207. 10.1016/j.neuroscience.2005.01.063

Jones, B. F., & Witter, M. P. (2007). Cingulate cortex projections to the parahippocampal region and hippocampal formation in the rat. Hippocampus, 17(10), 957–976. 10.1002/hipo.20330

Ju, G. & Swanson, L.W. (1989). Studies on the cellular architecture of the bed nuclei of the stria terminalis in the rat: I. Cytoarchitecture. J Comp Neurol, 280(4), 587–602. 10.1002/cne.902800409

Khan, A. M. (2013). Controlling feeding behavior by chemical or gene-directed targeting in the brain: what’s so spatial about our methods? Front Syst Neurosci, 7, 1–49. 10.3389/fnins.2013.00182

Khan A. M., Cheung, H. H., Gillard, E. R., Palarca, J. A., Welsbie, D. S., Gurd, J. W., & Stanley, B. G. (2004). Lateral hypothalamic signaling mechanisms underlying feeding stimulation: differential contributions of Src family tyrosine kinases to feeding triggered either by NMDA injection or by food deprivation. J Neurosci., 24(47):10603–10615. 10.1523/JNEUROSCI.3390-04.2004.

Khan, A. M., D’Arcy, C. E., & Olimpo, J. T. (2021). A historical perspective on training students to create standardized maps of novel brain structure: Newly-uncovered resonances between past and present research-based neuroanatomy curricula. Neurosci Lett, 759, 136052. 10.1016/j.neulet.2021.136052.

Khan, A. M., Grant, A. H., Martinez, A., Burns, G. A. P. C., Thatcher, B. S., Anekonda, V. T., Thompson, B. W., Roberts, Z. S., Moralejo, D. H., & Blevins, J. E. (2018a). Mapping molecular datasets back to the brain regions they are extracted from: Remembering the native countries of hypothalamic expatriates and refugees. Adv Neurobiol, 21, 101–193. 10.1007/978-3-319-94593-4_6.

Khan, A. M., Perez, J. G., Wells, C. E., & Fuentes, O. (2018b). Computer vision evidence supporting craniometric alignment of rat brain atlases to streamline expert-guided, first-order migration of hypothalamic spatial datasets related to behavioral control. Front Syst Neurosci, 12, 7. 10.3389/fnsys.2018.00007

Kim, U., & Lee, T. (2012). Topography of descending projections from anterior insular and medial prefrontal regions to the lateral habenula of the epithalamus in the rat. Eur J Neurosci, 35(8), 1253–1269. 10.1111/j.1460-9568.2012.08030.x

Kita, H., & Oomura, Y. (1982). An HRP study of the afferent connections to rat medial hypothalamic region. Brain Res Bull, 8(1), 53–62. 10.1016/0361-9230(82)90027-2

Köhler, C., Swanson, L.W., Haglund, L., & Wu, Y.-Y. (1985). The cytoarchitecture, histochemistry and projections of the tuberomammillary nucleus in the rat. Neuroscience, 16(1), 85–110. 10.1016/0306-4522(85)90049-1

Kölliker, A. (1896). *Handbuch der Gewebelehre des Menschen, 6th Edn*., *Zweiter Band: Nervensystem des Menschen und der Thiere* (Engelmann, Leipzig).

Krettek, J. E., & Price, J. L. (1977). The cortical projections of the mediodorsal nucleus and adjacent thalamic nuclei in the rat. J Comp Neurol, 171(2), 157–191. 10.1002/cne.901710204

Krieger, M. S., Condrad, L. C., & Pfaff, D. W. (1979). An autoradiographic study of the efferent connections of the ventromedial nucleus of the hypothalamus. J Comp Neurol, 183(4), 785–815. 10.1002/cne.901830408

Kuhlenbeck, H. (1927) *Vorlesungen über das Zentralnervensystem der Wirbeltiere* (Fischer, Jena).

Kuramoto, E., Pan, S., Furuta, T., Tanaka, Y. R., Iwai, H., Yamanaka, A., Ohno, S., Kaneko, T., Goto, T., & Hioki, H. (2016). Individual mediodorsal thalamic neurons project to multiple areas of the rat prefrontal cortex: A single neuron-tracing study using virus vectors. J Comp Neurol, 525(1),166–185. 10.1002/cne.24054

Lanciego, J. L., & Wouterlood, F. G. (2006). Multiple Neuroanatomical Tract-Tracing: Approaches for Multiple Tract-Tracing. In: Zaborszky, L., Wouterlood, F. G., Lanciego, J. L. (eds.) Neuroanatomical Tract-Tracing 3. New York: Springer. pp. 336–365.

Laubach, M., Amarante, L. M., Swanson, K., & White, S. R. (2018). What, if anything, is rodent prefrontal cortex? eNeuro, 5(5), ENEURO.0315-18.2018. 10.1523/eneuro.0315-18.2018

Lenhossék, M. (1887). Beobachtungen am Gehirn des Menschen. Anatomischer Anzeiger, 2, 450–461.

Leonard, C. M. (1969). The prefrontal cortex of the rat. I. Cortical projection of the mediodorsal nucleus. II. Efferent connections. Brain Res, 12(2), 321–343. 10.1016/0006-8993(69)90003-1

Leuret, F. & Gratiolet, P. (1839–1857). Anatomie Comparée du Système Nerveux Considéré dans ses Rapports avec l’Intelligence, 2 Vols. & Atlas (Baillière et Fils, Paris).

Loo, Y. T. (1931). The forebrain of the opossum, *Didelphis virginiana*. J Comp Neurol, 52(1), 1–148. 10.1002/cne.900520102

Lu, H., Zou, Q., Gu, H., Raichle, M. E., Stein, E. A., & Yang, Y. (2012). Rat brains also have a default mode network. Proc Natl Acad Sci U S A, 109(10), 3979–3984. 10.1073/pnas.1200506109

Luppi, P. H., Fort, P., & Jouvet, M. (1990). Iontophoretic application of unconjugated cholera toxin B subunit (CTB) combined with immunohistochemistry of neurochemical substances: a method for transmitter identification of retrogradely labeled neurons. Brain Res, 534(1–2), 209–224. 10.1016/0006-8993(90)90131-t

Magoul, R., Dubourg, P., Benjelloun, W., & Tramu, G. (1993a). Direct and indirect enkephalinergic synaptic inputs to the rat arcuate nucleus studied by combination of retrograde tracing and immunocytochemistry. Neuroscience, 55(4), 1055–1066. 10.1016/0306-4522(93)90319-b

Magoul, R., Onteniente, B., Benjelloun, W., & Tramu, G. (1993b). Tachykinergic afferents to the rat arcuate nucleus. A combined immunohistochemical and retrograde tracing study. Peptides, 14(2), 275–286. 10.1016/0196-9781(93)90042-f

Martinez A. (2017). The metabolically sentient arcuate nucleus: A functional, chemoarchitectural and connectional study in the adult male rat. Doctoral dissertation. University of Texas, El Paso.

Mandelbaum, G., Taranda, J., Haynes, T. M., Hochbaum, D. R., Huang, K. W., Hyun, M., Umadevi Venkataraju, K., Straub, C., Wang, W., Robertson, K., Osten, P., & Sabatini, B. L. (2019). Distinct cortical-thalamic-striatal circuits through the parafascicular nucleus. Neuron, 102(3), 636–652.e7. 10.1016/j.neuron.2019.02.035

Martínez, M., Espinoza, V. E., Garcia, V., Uribe, K. P., Negishi, K., Estevao, I.L., Carcoba, L. M., O’Dell, L. E., Khan, A. M., & Mendez, I. A. (2023). Withdrawal from repeated nicotine vapor exposure increases somatic signs of physical dependence, anxiety-like behavior, and brain reward thresholds in adult male rats. Neuropharmacology, 240, 109681. 10.1016/j.neuropharm.2023.109681.

McDonald, A. J. (1982). Cytoarchitecture of the central amygdaloid nucleus of the rat. J Comp Neurol, 208(4), 401–418. 10.1002/cne.902080409

McDonald, A. J. (1983). Neurons of the bed nucleus of the stria terminalis: a Golgi study in the rat. Brain Res Bull, 10(1), 111–120. 10.1016/0361-9230(83)90082-5

McDonald, A. J., Shammah-Lagnado, S. J., Shi, C., & Davis, M. (1999). Cortical afferents to the extended amygdala. Ann N Y Acad Sci, 877, 309–338. 10.1111/j.1749-6632.1999.tb09275.x

McKenna, J. T., & Vertes, R. P. (2004). Afferent projections to nucleus reuniens of the thalamus. J Comp Neurol, 480(2), 115–142. 10.1002/cne.20342

Mena, J. D., Sadeghian, K., & Baldo, B. A. (2011). Induction of hyperphagia and carbohydrate intake by µ-opioid receptor stimulation in circumscribed regions of frontal cortex. J Neurosci, 31(9), 3249–3260. 10.1523/jneurosci.2050-10.2011

Mena, J. D., Selleck, R. A., & Baldo, B. A. (2013). Mu-opioid stimulation in rat prefrontal cortex engages hypothalamic orexin/hypocretin-containing neurons, and reveals dissociable roles of nucleus accumbens and hypothalamus in cortically driven feeding. J Neurosci, 33(47), 18540–18552. 10.1523/jneurosci.3323-12.2013

Mickelsen, L. E., Flynn, W. F., Springer, K., Wilson, L., Beltrami, E. J., Bolisetty, M., Robson, P., & Jackson, C. (2020) Cellular taxonomy and spatial organization of the murine ventral posterior hypothalamus. Elife, 9, e58901. 10.7554/elife.58901

Miller, E. K., & Cohen, J. D. (2001). An integrative theory of prefrontal cortex function. Annu Rev Neurosci, 24, 167–202. 10.1146/annurev.neuro.24.1.167

Millhouse, O. E. (1969). A Golgi study of the descending medial forebrain bundle. Brain Res, 15(2), 341–363. 10.1016/0006-8993(69)90161-9

Mitrofanis, J., & Mikuletic, L. (1999). Organisation of thecortical projection to thezonaincerta of the thalamus. J Comp Neurol, 412(1),173–185. 10.1002/(SICI)1096-9861(19990913)412:1%3C173::AID-CNE13%3E3.0.CO;2-Q

Moga, M. M., Weis, R. P., & Moore, R. Y. (1995). Efferent projections of the paraventricular thalamic nucleus in the rat. J Comp Neurol, 359(2), 221–238. 10.1002/cne.903590204

Nauta, W. J. H. (1964). Some efferent connections of the prefrontal cortex in the monkey. In: Warren, J. M., Akert, K. (eds.) The Frontal Granular Cortex and Behavior. New York: McGraw Hill. pp. 397–409.

Nauta, W.J.H. & Haymaker, W. (1969). Hypothalamic nuclei and fiber connections. In: Haymaker, W., Anderson, E., & Nauta, W.J.H. (Eds.) The Hypothalamus (Charles C. Thomas, Springfield IL), pp. 136–209.

Negishi, K. (2016). Macro- and mesoscale analysis of connections between the cingulate region and the lateral hypothalamic area: tracer co-injection and chemoarchitectural studies in the adult male rat. Master’s Thesis. 56 pp. The University of Texas at El Paso. Open Access Theses & Dissertations, 709. https://scholarworks.utep.edu/open_etd/709.

Negishi, K. (2023). Connectional analysis of brain regions associated with feeding. Doctoral dissertaion. The University of Texas at El Paso. *Open Access Theses & Dissertations*, 3835. https://scholarworks.utep.edu/open_etd/3835.

Negishi, K., Hamdan, J., & Khan, A. M. (2015). Initial chemoarchitectural and connectional characterization of polymodal association cortical structures with the diencephalon: Immunohistochemical and tract tracing studies. Program No. 616.14. 2015 *Neuroscience Meeting Planner*. Chicago, IL. Society for Neuroscience, 2015. Online.

Negishi, K., & Khan, A. M. (2017). Mapping the connections between the medial prefrontal cortex and the diencephalon: A combined anterograde and retrograde tract-tracing study in the adult male rat. Program No. 604.07. 2017 *Neuroscience Meeting Planner*. Washington, DC. Society for Neuroscience, 2017. Online.

Negishi, K., & Khan, A. M. (2019). The contribution of axonal projections from infralimbic area neuronal populations to the medial forebrain bundle: analysis of morphology and interactions with hypocretin/orexin neurons in the rat. Program No. 149.24. 2019 *Neuroscience Meeting Planner.* Chicago, IL: Society for Neuroscience, 2019. Online.

Nieuwenhuys, R., Geeraedts, L. M. G., & Veening, J. G. (1982). The medial forebrain bundle of the rat. I. General introduction. J Comp Neurol, 206:49–81.

Nissl, F. (1913). Die Grosshirnanteile des Kaninchens. Archiv für Psychiatrie und Nervenkrankheiten, 52, 867–953.

NRC [National Research Council (US) Committee for the Update of the Guide for the Care and Use of Laboratory Animals]. (2011). Guide for the care and use of laboratory animals*, 8th edition*. Washington, D.C.: National Academies Press.

Paus, T., Zijdenbos, A., Worsley, K., Collins, D. L., Blumenthal, J., Giedd, J. N., Rapoport, J. A., & Evans, C. (1999). Structural maturation of neural pathways in children and adolescents: in vivo study. Science, 283(5409),1908–11. 10.1126/science.283.5409.1908

Paxinos, G. & Watson, C. (1986). The Rat Brain in Stereotaxic Coordinates, 2nd Edn. New York: Academic Press.

Paxinos, G., & Watson, C. (2014). The Rat Brain in Stereotaxic Coordinates, 7th Edn. San Diego: Academic Press.

Petrovich, G. D. (2011). Forebrain circuits and control of feeding by learned cues. Neurobiol Learn Mem, 95(2),152–158. 10.1016/j.nlm.2010.10.003

Petrovich, G. D., Canteras, N. S., & Swanson, L. W. (2001). Combinatorial amygdalar inputs to hippocampal domains and hypothalamic behavior circuits. Brain Res Rev, 38(1–2), 247–289. 10.1016/s0165-0173(01)00080-7

Petrovich, G. D., Hobin, M. P., & Reppucci, C. J. (2012). Selective Fos induction in hypothalamic orexin/ hypocretin, but not melanin-concentrating hormone neurons, by a learned food-cue that stimulates feeding in sated rats. Neuroscience, 224,70–80. 10.1016/j.neuroscience.2012.08.036

Petrovich, G. D., Ross, C. A., Holland, P. C., & Gallagher, M. (2007). Medial prefrontal cortex is necessary for an appetitive contextual conditioned stimulus to promote eating in sated rats. J Neurosci, 27(24),6436–41. 10.1523/jneurosci.5001-06.2007

Preuss, T. M., & Goldman-Rakic, P. S. (1987). Crossed corticothalamic and thalamocortical connections of macaque prefrontal cortex. J Comp Neurol, 257(2),269–81. 10.1002/cne.902570211

Radley, J. J., Gosselink, K. L., & Sawchenko, P. E. (2009). A discrete GABAergic relay mediates medial prefrontal cortical inhibition of the neuroendocrine stress response. J Neurosci, 29(22),7330–7340. 10.1523/jneurosci.5924-08.2009

Reep, R. L., Corwin, J. V., Hashimoto, A., & Watson, R. T. (1984). Afferent connections of medial precentral cortex in the rat. Neurosci Lett, 44,247–252. 10.1016/0304-3940(84)90030-2

Reppucci, C. J. & Petrovich, G. D. (2016). Organization of connections between the amygdala, medial prefrontal cortex, and lateral hypothalamus: a single and double retrograde tracing study in rats. Brain Struct Funct, 221(6),2937–62. 10.1007/s00429-015-1081-0

Rioch, D.M. (1929). Studies on the diencephalon of Carnivora. Part II: Certain nuclear configurations and fiber connections of the subthalamus and midbrain of the dog and cat. J Comp Neurol, 49, 121–153.

Risold, P.Y. & Swanson, L.W. (1995). Cajal’s nucleus of the stria medullaris: characterization by in situ hybridization and immunohistochemistry for enkephalin. J Chem Neuroanat, 9(4), 235–240. 10.1016/0891-0618(95)00083-6

Risold, P.Y. & Swanson, L.W. (1997). Chemoarchitecture of the rat lateral septal nucleus. Brain Res Rev, 24(2–3), 91–113. 10.1016/s0165-0173(97)00008-8

Risold, P.Y., Thompson, R.H., & Swanson, L.W. (1997). The structural organization of connections between hypothalamus and cerebral cortex. Brain Res Rev, 24(2–3), 197–254. 10.1016/s0165-0173(97)00007-6

Rose, J.E. & Woolsey, C.N. (1948). Structure and relations of limbic cortex and anterior thalamic nuclei in rabbit and cat. J Comp Neurol, 89(3), 279–347. 10.1002/cne.900890307

Sano, T. (1910). Beitrag zur vergleichenden Anatomie der Substantia nigra, des Corps Luysii und der Zona incerta. Monatsschrift für Psychiatrie und Neurologie, 28, 26–31.

Risold, P.-Y., Canteras, N. S., & Swanson, L. W. (1994). Organization of projections from the anterior hypothalamic nucleus: a *Phaseolus vulgaris*-leucoagglutinin study in the rat. J Comp Neurol, 348(1),1–40. 10.1002/cne.903480102

Santarelli, A. J., Khan, A. M., & Poulos, A. M. (2018). Contextual fear retrieval-induced Fos expression across early development in the rat: An analysis using established nervous system nomenclature ontology. Neurobiol Learn Mem, 155, 42–49. 10.1016/j.nlm.2018.05.015.

Saper, C. B., Swanson, L. W., & Cowan, W. M. (1976). The efferent connections of the ventromedial nucleus of the hypothalamus of the rat. J Comp Neurol, 169(4),409–42. 10.1016/j.neures.2014.10.016

Saper, C. B., & Levisohn, D. (1983). Afferent connections of the median preoptic nucleus in the rat: anatomical evidence for a cardiovascular integrative mechanism in the anteroventral third ventricular (AV3V) region. Brain Res, 288(1–2),21–31. 10.1016/0006-8993(83)90078-1

Sapolsky, R. M. (2004). The frontal cortex and the criminal justice system. Phil Trans R Soc Lond B Biol Sci, 359(1451),1787–96. 10.1098/rstb.2004.1547

Sawchenko, P. E., & Swanson, L. W. (1983). The organization of forebrain afferents to the paraventricular and supraoptic nuclei of the rat. J Comp Neurol, 218(2),121–44. 10.1002/cne.902180202

Scalia, F. & Winans, S.S. (1975). The differential projections of the olfactory bulb and accessory olfactory bulb in mammals. J Comp Neurol, 161(2), 31–56. 10.1002/cne.901610105

Schmitt, LI., Wimmer, R. D., Nakajima, M., Happ, M., Mofokham, S., & Halassa, M. M. (2017). Thalamic amplification of cortical connectivity sustains attentional control. Nature, 545,219–223. 10.1038/nature22073

Schwalbe, G. (1881). *Lehrbuch der Neurology* (Besold, Erlangen).

Sesack, S. R., Deutch, A. Y., Roth, R. H., & Bunney, B. S. (1989). Topographical organization of the efferent projections of the medial prefrontal cortex in the rat: an anterograde tract-tracing study with *Phaseolus vulgaris* leucoagglutinin. J Comp Neurol, 290(2),213–42. 10.1002/cne.902900205

Sherin, J. E., Elmquist, J. K., Torrealba, F., & Saper, C.B. (1998). Innervation of histaminergic tuberomammillary neurons by GABAergic and galaninergic neurons in the ventrolateral preoptic nucleus of the rat. J Neurosci, 18(12), 4705–4721. 10.1523/jneurosci.18-12-04705.1998

Shibata, H. (1993). Efferent projections from the anterior thalamic nuclei to the cingulate cortex in the rat. J Comp Neurol, 330(4),533–42. 10.1002/cne.903300409

Shibata, H., & Naito, J. (2005). Organization of anterior cingulate and frontal cortical projections to the anterior and laterodorsal thalamic nuclei in the rat. Brain Res, 1059(1),93–103. 10.1016/j.brainres.2005.08.025

Shibata, H., & Naito, J. (2008). Organization of anterior cingulate and frontal cortical projections to the retrosplenial cortex in the rat. J Comp Neurol, 506(1),30–45. 10.1002/cne.21523

Shimogawa, Y., Sakuma, Y., & Yamanouchi, K. (2015). Efferent and afferent connections of the ventromedial hypothalamic nucleus determined by neural tracer analysis: implications for lordosis regulation in female rats. Neurosci Res, 91,19–33. 10.1016/j.neures.2014.10.016

Shiosaka, S., Tohyama, M., Takagi, H., Takahashi, Y., Saitoh, Y., Sakumoto, T., Nakagawa, H., & Shimizu, N. (1980). Ascending and descending components of the medial forebrain bundle in the rat as demonstrated by the horesradish peroxidase-blue reaction. Exp Brain Res, 39, 377–388.

Silverman, A. J., Hoffman, D. L., & Zimmerman, E. A. (1981). The descending afferent connections of the paraventricular nucleus of the hypothalamus (PVN). Brain Res Bull, 6(1),47–61. 10.1016/s0361-9230(81)80068-8

Simerly, R. B., & Swanson, L. W. (1986). The organization of neural inputs to the medial preoptic nucleus of the rat. J Comp Neurol, 246(3), 312–342. 10.1002/cne.902460304

Simerly, R. B., & Swanson, L. W. (1988). Projections of the medial preoptic nucleus: a *Phaseolus vulgaris* leucoagglutinin anterograde tract-tracing study in the rat. J Comp Neurol, 270(2), 209–242. 10.1002/cne.902700205

Simerly, R.B., Swanson, L.W., & Gorski, R.A. (1984). Demonstration of a sexual dimorphism in the distribution of serotonin-immunoreactive fibers in the medial preoptic nucleus of the rat. J Comp Neurol, 225(2), 151–166. 10.1002/cne.902250202

Simmons, D.M. & Swanson, L.W. (2008). High resolution paraventricular nucleus serial section model constructed within a traditional rat brain atlas. Neurosci Lett, 438(1), 85–89. 10.1016/j.neulet.2008.04.057

Simmons, D. M., & Swanson, L. W. (2009a). Comparing histological data from different brains: sources of error and strategies for minimizing them. Brain Res Rev, 60(2),349–367. 10.1016/j.brainresrev.2009.02.002

Simmons, D. M., & Swanson, L. W. (2009b). Comparison of the spatial distribution of seven types of neuroendocrine neurons in the rat paraventricular nucleus: Toward a global 3D model. J Comp Neurol, 516(5),423–41. 10.1002/cne.22126

Sita, L. V., Elias, C. F., & Bittencourt, J. C. (2007). Connectivity pattern suggests that incerto-hypothalamic area belongs to the medial hypothalamic system. Neuroscience, 148(4),949–969. 10.1016/j.neuroscience.2007.07.010

Spiegel, E.A. & Zwieg, H. (1919). Zur Cytoarchitektonic des Tuber cinereum. Arbeiten aus dem Neurologischen Institute an der Wiener Universiteit, 22, 278–295.

Swanson, L. W. (1976). An autoradiographic study of the efferent connections of the preoptic region in the rat. J Comp Neurol, 167(2), 227–256. 10.1002/cne.901670207

Swanson, L.W. (1982). The projections of the ventral tegmental area and adjacent regions: a combined fluorescent retrograde tracer and immunofluorescence study in the rat. Brain Res Bull, 9(1–6), 321–353. 10.1016/0361-9230(82)90145-9

Swanson, L. W. (1992). Brain Maps: Structure of the Rat Brain. Amsterdam: Elsevier.

Swanson, L. W. (1998). Brain Maps: Structure of the Rat Brain. A Laboratory Guide with Printed and Electronic Templates for Data, Models and Schematics (2nd ed.). Amsterdam: Elsevier.

Swanson, L. W. (2000). Cerebral hemisphere regulation of motivated behavior. Brain Res, 886,113–164. 10.1016/s0006-8993(00)02905-x

Swanson, L.W. (2004). Brain Maps: Structure of the Rat Brain. A Laboratory Guide with Printed and Electronic Templates for Data, Models and Schematics, 3rd Edn. (Elsevier, Amsterdam), 215 pp. with CD-ROM, *Brain Maps: Computer Graphics Files 3.0*.

Swanson, L. W. (2015). Neuroanatomical Terminology: A lexicon of classical origins and historical foundations. Oxford: Oxford University Press.

Swanson, L. W. (2018). *Brain maps 4.0—Structure of the rat brain*: An open access atlas with global nervous system nomenclature ontology and flatmaps. J Comp Neurol, 526(6), 935–943. 10.1002/cne.24381

Swanson, L. W. & Kuypers, H. G. J. M. (1980). A direct projection from the ventromedial nucleus and retrochiasmatic area of the hypothalamus to the medulla and spinal cord of the rat. Neurosci Lett, 17(3), 307–312. 10.1016/0304-3940(80)90041-5

Swanson, L.W. & Petrovich, G.D. (1998). What is the amygdala? Trends Neurosci, 21(8), 323–331. 10.1016/s0166-2236(98)01265-x

Swanson, L. W., Sanchez-Watts, G., & Watts, A. G. (2005). Comparison of melanin-concentrating hormone and hypocretin/orexin mRNA expression patterns in a new parceling scheme of the lateral hypothalamic zone. Neurosci Lett, 387(2), 80–84. 10.1016/j.neulet.2005.06.066

Swanson, L. W., & Sawchenko, P. E. (1983). Hypothalamic integration: Organization of the paraventricular and supraoptic nuclei. Ann Rev Neurosci, 6, 269–324. 10.1146/annurev.ne.06.030183.001413

Swanson, L.W. & Simmons, D. M. (1989). Differential steroid hormone and neural influences on peptide mRNA levels in CRH cells of the paraventricular nucleus: a hybridization histochemical study in the rat. J Comp Neurol, 285(4), 413–435. 10.1002/cne.902850402

Swanson, L. W., Sporns, O., & Hahn, J. D. (2016). Network architecture of the cerebral nuclei (basal ganglia) association and commissural connectome. Proc Natl Acad Sci U S A, 113(40), E5971–E5981. 10.1073/pnas.1613184113

Swanson, L. W., Hahn, J. D., & Sporns, O. (2017). Organizing principles for the cerebral cortex network of commissural and association connections. Proc Natl Acad Sci U S A, 114(45), E9692–E9701. 10.1073/pnas.1712928114

Swanson, L. W., Hahn, J. D., Jeub, L. G. S., Fortunato, S., & Sporns, O. (2018). Subsystem organization of axonal connections within and between the right and left cerebral cortex and cerebral nuclei (endbrain). Proc Natl Acad Sci U S A, 115(29), E6910–E16919. 10.1073/pnas.1807255115

Swanson, L. W., Sporns, O., & Hahn, J. D. (2019a). The network architecture of rat intrinsic interbrain (diencephalic) macroconnections. Proc Natl Acad Sci U S A, 116(52), 26991–27000. 10.1073/pnas.1915446116

Swanson, L. W., Sporns, O., & Hahn, J. D. (2019b). The network organization of rat intrathalamic macroconnections and a comparison with other forebrain divisions. Proc Natl Acad Sci U S A, 116(27), 13661–13669. 10.1073/pnas.1905961116

Swanson, L. W., Hahn, J. D., & Sporns, O. (2020). Structure-function subsystem models of female and male forebrain networks integrating cognition, affect, behavior, and bodily functions. Proc Natl Acad Sci U S A, 117(49), 31470–31481. 10.1073/pnas.2017733117

Swanson, L. W., Hahn, J. D., & Sporns, O. (2021). Subsystem macroarchitecture of the intrinsic midbrain neural network and its tectal and tegmental subnetworks. Proc Natl Acad Sci U S A, 118(20), e2101869118. 10.1073/pnas.2101869118

Swanson, L. W., Hahn, J. D., & Sporns, O. (2022). Structure-function subsystem model and computational lesions of the central nervous system’s rostral sector (forebrain and midbrain). Proc Natl Acad Sci U S A. 119(45):e2210931119. 10.1073/pnas.2210931119.

Swanson, L. W., Hahn, J. D., & Sporns, O. (2023). Intrinsic circuitry of the rhombicbrain (central nervous system’s intermediate sector) in a mammal. Proc Natl Acad Sci U S A, 120(52):e2313997120. 10.1073/pnas.2313997120.

Swanson, L. W., Hahn, J. D., & Sporns, O. (2024a). Network architecture of intrinsic connectivity in a mammalian spinal cord (the central nervous system’s caudal sector). Proc Natl Acad Sci U S A, 121(5):e2320953121. 10.1073/pnas.2320953121.

Swanson, L. W., Hahn, J. D., & Sporns, O. (2024b). Neural network architecture of a mammalian brain. Proc Natl Acad Sci U S A, 121(39):e2413422121. 10.1073/pnas.2413422121.

Sylvester, C. M., Krout, K. E., & Loewy, A. D. (2002). Suprachiasmatic nucleus projection to the medial prefrontal cortex: a viral transneuronal tracing study. Neuroscience, 114(4), 1071–1080. 10.1016/s0306-4522(02)00361-5

Tapia, G. P.*, Agostinelli, L. J.*, Chenausky, S. D., Salcido Padilla, J. V., Navarro, V. I., Alagh, A., Si, G., Thompson, R. H., Balivada, S., & Khan, A. M. (2023). Glycemic challenge is associated with the rapid cellular activation of the locus ceruleus and nucleus of solitary tract: Circumscribed spatial analysis of phosphorylated MAP kinase immunoreactivity. J Clin Med, 12, 2483. 10.3390/jcm12072483. *co-first authors

Tarin, P. (1753). *Dictionnaire Anatomique suivi d’une Bibliotheque Anatomique et Physiologique* (Briasson, Paris).

Ter Horst, G. J., Groenewegen, H. J., Karst, H., & Luiten, P. G. (1984). *Phaseolus vulgaris* leuco-agglutinin immunohistochemistry. A comparison between autoradiographic and lectin tracing of neuronal efferents. Brain Res 307(1–2), 379–383. 10.1016/0006-8993(84)90500-6

Ter Horst, G. J., & Luiten, P. G. (1986). The projections of the dorsomedial hypothalamic nucleus in the rat. Brain Res Bull, 16(2), 231–248. 10.1016/0361-9230(86)90038-9

Thompson, R. H., Canteras, N. S., & Swanson, L. W. (1996). Organization of projections from the dorsomedial nucleus of the hypothalamus: a PHAL study in the rat. J Comp Neurol, 376(1), 143–173. https://doi.org/10.1002/(sici)1096-9861(19961202)376:1%3C143::aid-cne9%3E3.0.co;2-3

Thompson, R. H., & Swanson, L. W. (1998). Organization of inputs to the dorsomedial nucleus of the hypothalamus: a reexamination with Fluorogold and PHAL in the rat. Brain Res Rev, 27(2), 89–118. 10.1016/s0165-0173(98)00010-1

Thompson, R. H., & Swanson, L. W. (2010). Hypothesis-driven structural connectivity analysis supports network over hierarchical model of brain architecture. Proc Natl Acad Sci U S A, 107, 15235–15239. 10.1073/pnas.1009112107

Tilney, F. (1936). The development and constituents of the human hypophysis. Bulletin of the Neurological Institute of New York, 5, 387–436.

Toth, M., Fuzesi, T., Halasz, J., Tulogdi, A., & Haller, J. (2010). Neural inputs of the hypothalamic “aggression area” in the rat. Behav Brain Res, 215(1), 7–20. 10.1016/j.bbr.2010.05.050

Tsai, C. (1925). The optic tracts and centers of the opossum, *Didelphis virginiana*. J Comp Neurol, 39(2), 173–216. 10.1002/cne.900390202

Tu, W., Ma, Z., Ma, Y., Dopfel, D., & Zhang, N. (2021). Suppressing anterior cingulate cortex modulates default mode network and behavior in awake rats. Cereb Cortex, 31(1), 312–323. 10.1093/cercor/bhaa227

Urstadt, K. R., & Berridge, K. C. (2020). Optogenetic mapping of feeding and self-stimulation within the lateral hypothalamus of the rat. PLoS One, 15(1), e0224301. 10.1371/journal.pone.0224301

van der Werf, Y. D., Witter, M. P., & Groenewegen, H. J. (2002). The intralaminar and midline nuclei of the thalamus. Anatomical and functional evidence for participation in processes of arousal and awareness. Brain Res Rev, 39, 107–140. 10.1016/s0165-0173(02)00181-9

van Groen, T., Kadish, I., & Wyss, J. M. (1999). Efferent connections of the anteromedial nucleus of the thalamus of the rat. Brain Res Rev, 30(1), 1–26. 10.1016/s0165-0173(99)00006-5

Vanderwolf, C. H., Kolb, B., & Cooley, R. K. (1978). Behavior of the rat after removal of the neocortex and hippocampal formation. J Comp Physiol Psychol, 92(1), 156–175. 10.1037/h0077447

Varela, C., Kumar, S., Yang, J. Y., & Wilson, M. A. (2014). Anatomical substrates for direct interactions between hippocampus, medial prefrontal cortex, and the thalamic nucleus reuniens. Brain Struct Funct, 219(3), 911–929. 10.1007/s00429-013-0543-5

Vertes, R. P. (2002). Analysis of projections from the medial prefrontal cortex to the thalamus in the rat, with emphasis on nucleus reuniens. J Comp Neurol, 442(2), 163–187. 10.1002/cne.10083

Vertes, R. P. (2004). Differential projections of the infralimbic and prelimbic cortex in the rat. Synapse, 51(1), 32–58. 10.1002/syn.10279

Vertes, R. P., Crane, A. M., Colom, L. V., & Bland, B. H. (1995). Ascending projections of the posterior nucleus of the hypothalamus: PHAL analysis in the rat. J Comp Neurol, 359(1), 90–116. 10.1002/cne.903590107

Vertes, R. P., Hoover, W. B., Do Valle, A. C., Sherman, A., & Rodriguez, J. J. (2006). Efferent projections of reuniens and rhomboid nuclei of the thalamus in the rat. J Comp Neurol, 499(5), 768–796. 10.1002/cne.21135

Vertes, R. P., & Hoover, W. B. (2008). Projections of the paraventricular and paratenial nuclei of the dorsal midline thalamus in the rat. J Comp Neurol, 508(2), 212–237. 10.1002/cne.21679

Vertes, R. P., Hoover, W. B., & Rodriguez, J. J. (2012). Projections of the central medial nucleus of the thalamus in the rat: Node in cortical, striatal and limbic forebrain circuitry. Neuroscience, 219, 120–136. 10.1016/j.neuroscience.2012.04.067

Vogt, B. A., & Peters, A. (1981). Form and distribution of neurons in the rat cingulate cortex: areas 32, 24, and 29. J Comp Neurol, 195(4), 603–625. 10.1002/cne.901950406

Vogt, B. A., & Miller, M. W. (1983). Cortical connections between rat cingulate cortex and visual, motor, and postsubicular cortices. J Comp Neurol, 216(2), 192–210. 10.1002/cne.902160207

Vogt, C. (1909) La myéloarchitecture du thalamus du cercopithèque. Journal für Psychologie und Neurologie, 12 (Suppl.), 285–324.

Wagner, C. K., Eaton, M. J., Moore, K. E., & Lookingland, K. J. (1995). Efferent projections from the region of the medial zona incerta containing A13 dopaminergic neurons: a PHA-L anterograde tract-tracing study in the rat. Brain Res, 677(2), 229–237. 10.1016/0006-8993(95)00128-d

Wang, C. C., & Shyu, B. C. (2004). Differential projections from the mediodorsal and centrolateral thalamic nuclei to the frontal cortex in rats. Brain Res, 995(2), 226–235. 10.1016/j.brainres.2003.10.006

Wang, P. Y. & Zhang, F. C. (1995). Outlines and Atlas of Learning Rat Brain Slices (Westnorth University Press, China).

Wang, Q., Henry, A. M., Harris, J. A., Oh, S. W., Joines, K. M., Nyhus, J., Hirokawa, K. E., Dee, N., Mortrud, M., Parry, S., Ouellette, B., Caldejon, S., Bernard, A., Jones, A. R., Zeng, H., & Hohmann, J. G. (2014). Systematic comparison of adeno-associated virus and biotinylated dextran amine reveals equivalent sensitivity between tracers and novel projection targets in the mouse brain. J Comp Neurol, 522(9), 1989– 2012. 10.1002/cne.23567

Watts, A. G., Kanoski, S. E., Sanchez-Watts, G., & Langhans, W. (2022). The physiological control of eating: signals, neurons, and networks. Physiol Rev, 102(2), 689–813. 10.1152/physrev.00028.2020

Watts, A.G., Swanson, L.W, & Sanchez-Watts, G. (1987). Efferent projections of the suprachiasmatic nucleus: I. Studies using anterograde transport of *Phaseolus vulgaris* leucoagglutinin in the rat. J Comp Neurol, 258(2), 204–229. 10.1002/cne.902580204

Wenzel, J. & Wenzel, K. (1812). *De Penitiori Structura Cerebri Hominis et Brutorum* (Cotta, Tübingen).

Yoshida, A., Dostrovsky, J. O., & Chiang, C. Y. (1992). The afferent and efferent connections of the nucleus submedius in the rat. J Comp Neurol, 324(1), 115–133. 10.1002/cne.903240109

Yoshida, K., McCormack, S., España, R. A., Crocker, A., & Scammell, T. E. (2006). Afferents to the orexin neurons of the rat brain. J Comp Neurol, 494(5), 845–861. 10.1002/cne.20859

Ziehen, T. (1897–1901). Das Centralnervensystem der Monotremen und Marsupialier. Denkschriften der Medicinisch-naturwissenschaftlichen Gesellschaft zu Jena, 6, 1–187, 677–728.

Zséli, G., Vida, B., Martinez, A., Lechan, R. M., Khan, A. M., & Fekete, C. (2016). Elucidation of the anatomy of a satiety network: Focus on connectivity of the parabrachial nucleus in the adult rat. J Comp Neurol, 524(14), 2803–2827. 10.1002/cne.23992.

