## Supplemental Figure 1 for "Topographic organization of bidirectional connections between the cingulate region (infralimbic area and anterior cingulate area, dorsal part) and the interbrain (diencephalon) of the adult male rat"

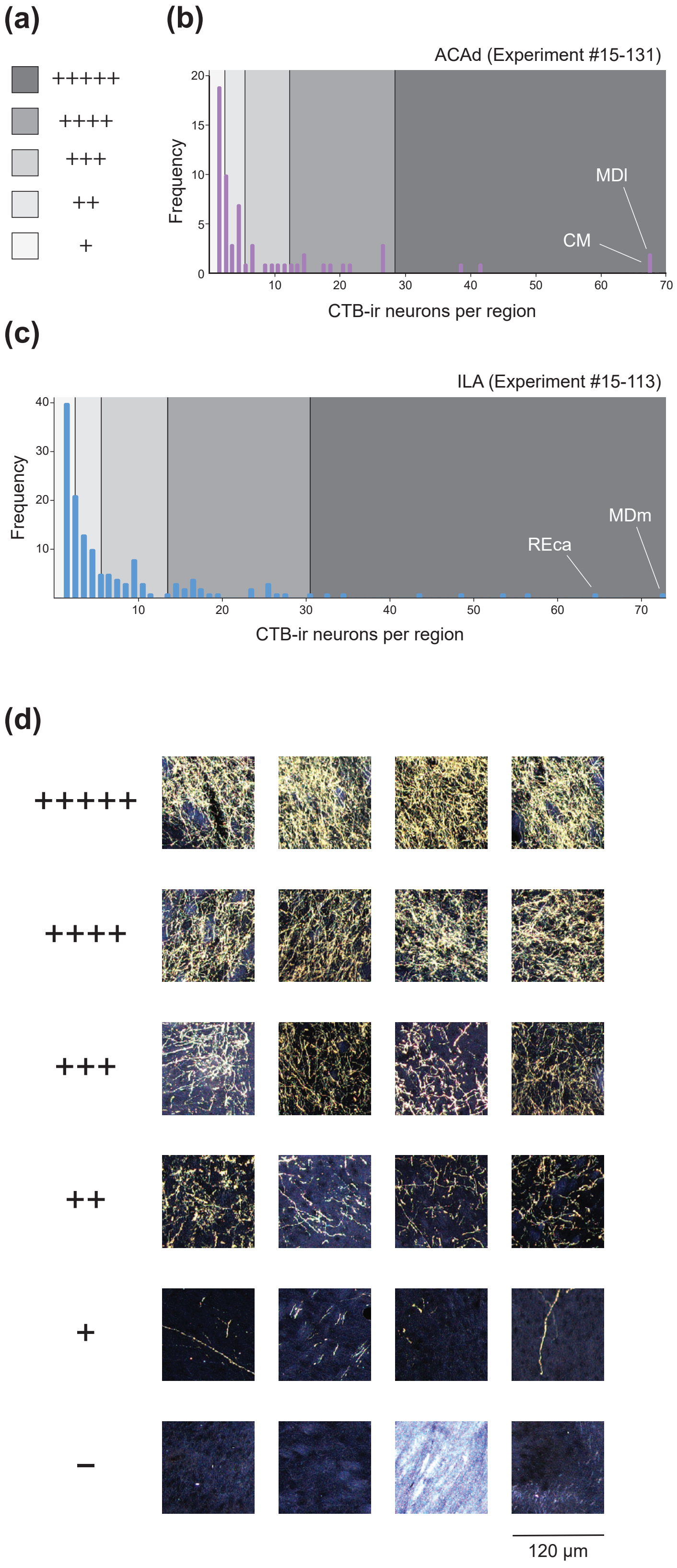


**Supplemental Figure 1.** Strategies for obtaining semi-quantitative scores (0–5 scale) for CTB **(a–c)** and PHAL **(d)**. **(a)** Shading code corresponding to symbols used in **Table 2**. **(b, c)** Histograms showing frequency of CTB-ir neuron counts per region across each atlas level for CTB injections in the ACAd **(b)** and ILA **(c)**. Shaded zones follow numerical ranges for scores in **(a)**. **(d)** Representative darkfield photomicrographs showing examples for each axon density score. Images were sampled using 120 × 120 μm bins.
